# Multiple merger genealogies in outbreaks of *Mycobacterium tuberculosis*

**DOI:** 10.1101/2019.12.21.885723

**Authors:** F. Menardo, S. Gagneux, F. Freund

**Affiliations:** Department of Medical Parasitology and Infection Biology, Swiss Tropical and Public Health Institute, Basel, Switzerland; University of Basel, Basel, Switzerland; Institute of Plant Breeding, Seed Science and Population Genetics, University of Hohenheim, Stuttgart, Germany

**Author notes:** Equal contribution, corresponding authors.

**Keywords:** *Mycobacterium tuberculosis*, demographic inference, multiple merger coalescent, Approximate Bayesian Computation, Random Forest

## Abstract

The Kingman coalescent and its developments are often considered among the most important advances in population genetics of the last decades. Demographic inference based on coalescent theory has been used to reconstruct the population dynamics and evolutionary history of several species, including *Mycobacterium tuberculosis* (MTB), an important human pathogen causing tuberculosis. One key assumption of the Kingman coalescent is that the number of descendants of different individuals does not vary strongly, and violating this assumption could lead to severe biases caused by model misspecification. Individual lineages of MTB are expected to vary strongly in reproductive success because 1) MTB is potentially under constant selection due to the pressure of the host immune system and of antibiotic treatment, 2) MTB undergoes repeated population bottlenecks when it transmits from one host to the next, and 3) some hosts show much higher transmission rates compared to the average (“super-spreaders”).

Here we used an Approximate Bayesian Computation approach to test whether multiple merger coalescents (MMC), a class of models that allow for large variation in reproductive success among lineages, are more appropriate models to study MTB populations. We considered eleven publicly available whole genome sequence data sets sampled from local MTB populations and outbreaks, and found that MMC had a better fit compared to the Kingman coalescent for ten of the eleven data sets. These results indicate that the null model for analyzing MTB outbreaks should be reassessed, and that past findings based on the Kingman coalescent need to be revisited.

## Introduction

The coalescent is a stochastic mathematical model that formally describes the shapes of the expected genealogies in a population (Kingman 1982a). The original formulation of Kingman has been extended to include different evolutionary processes such as fluctuations in population size (Griffith and Tavaré 1994), population subdivision and migration (Wilkinson-Herbots 1998), recombination (Hudson 1983), and selection (Kaplan et al. 1988, Neuhauser and Krone 1997).

Although the genealogy of a sample is typically unknown, mutational models can be superimposed onto the coalescent to describe DNA sequence polymorphisms. These are generally easy to obtain from natural populations, thus opening the possibility of data-based statistical inference.

Applications of the coalescent include the study of the evolutionary histories and population dynamics of a variety of taxa (Kuhner 2009), including humans (Li and Durban 2001, Excoffier et al. 2013) and pathogens (Pybus et al. 2001, Joy et al. 2003), and the identification of genetic loci under selection (Biswas and Akey 2006, Hernandez et al. 2011).

One of the assumptions of the Kingman coalescent is that the variation in reproductive success among individuals is small, such as at most one pair of sampled lineages can find a common ancestor for any single time point on the coalescent time scale.

This assumption is relaxed in a more general class of models, the so-called multiple merger coalescents (MMC). MMC have been developed to model scenarios in which the variance in reproductive success is large enough to cause the coalescence of more than two lineages at the same time point on the coalescent time scale (Möhle and Sagitov 2001, see Tellier and Lemaire 2014 for a review). Some of the underlying discrete generations models leading to MMC allow for very large offspring numbers of one or more individuals in a single generation (Schweinsberg 2003, Eldon and Wakeley 2006). However, MMC genealogies can also arise if one individual has many descendants in a relatively small number of generations, so that this family leads to multiple mergers after collapsing discrete generations to arrive at the timescale of the continuous time coalescent. In this manuscript, we will refer to “skewed offspring distribution” to indicate variation in reproductive success that lead to multiple merger genealogies. For details on how coalescent models arise from discrete generation models we refer to the relevant mathematical literature (Kingman 1982b, Griffith and Tavaré 1994, Möhle and Sagitov 2001, Freund 2020).

Compared to the Kingman coalescent, MMC have been proposed to be more appropriate models to investigate marine organisms with sweepstakes reproduction (Sargsyan and Wakeley 2008), agricultural pathogens with recurrent seasonal bottlenecks (Tellier and Lemaire 2014), loci under positive selection (Durrett and Schweinsberg 2005), and rapidly adapting pathogens (Neher and Hallatschek 2013).

Despite a growing interest in MMC, there are few studies that used genetic polymorphisms to test whether MMC are indeed better fitting models compared to the Kingman coalescent. Signatures of MMC have been detected at the *creatin kinase muscle type A* locus of the Atlantic cod *(Gadus morhua;* Árnason and Halldórsdóttir 2015), in the mitochondrial genome of Japanese sardines *(Sardinops melanostictus;* Niwa et al. 2016), in populations of breast cancer cells (Kato et al. 2017), and in the B-cell repertoire response to viruses such as HIV-1 and influenza (Nourmmohammad et al. 2019, Horns et al. 2019). While MMC are theoretically appealing genealogy models for pathogen samples (Irwin et al. 2016, Rocha 2018, Neher and Walczak 2018), their fit to observed data in pathogen populations has not been investigated so far. Only very recently, MMC have been used to study the within-host genetic diversity of *Mycobacterium tuberculosis* (MTB), a major human pathogen causing tuberculosis (Morales-Arce et al. 2020).

Here we look for evidence of MMC in between-host populations of *Mycobacterium tuberculosis.* Between-host populations of MTB are expected to have a skewed offspring distribution because of three reasons: 1) MTB is an obligate pathogen, and therefore potentially constantly adapting under the pressure of the host immune system and of antibiotic treatment (Gagneux 2018). 2) Super-spreaders; these are patients responsible for a very large number of secondary infections compared to the average (Gardy et al. 2011, Walker et al. 2013, Ypma et al. 2013, Stucki et al. 2015, Lee et al. 2015a, Lee et al. 2020), thus causing a large variance of the pathogen’s offspring size. 3) MTB undergoes repeated bottlenecks when transmitting from one host to the next, with a few bacteria, and potentially as few as one, founding the entire population infecting the new host (Lin et al. 2014).

Additionally, a low genetic diversity and an excess of rare variants (singletons) have been reported in MTB (Hershberg et al. 2008, Pepperell et al. 2013), and both are known signatures of MMC genealogies (Tellier and Lemaire 2014).

Methods based on the Kingman coalescent are often used in population genetic analyses of MTB. For example: 1) The Bayesian Skyline Plot (BSP, Drummond et al. 2005) has been used to infer past population dynamics in tuberculosis outbreaks, finding evidence for constant effective population size (Bainomugisa et al. 2018), rapid effective population growth (Eldholm et al. 2015, Folkvardsen et al. 2017, Brown et al. 2019) or slow effective population decline (Lee et al. 2015b). 2) Different methods have been used to infer the demographic history of the global MTB population (Pepperell et al. 2013, Comas et al. 2013, Bos et al. 2014) and of individual MTB lineages (Kay et al. 2015, Luo et al. 2015, Merker et al. 2015, Merker et al. 2018, Liu et al. 2018, Huang et al. 2019, O’Neill et al. 2019, Mulholland et al. 2019), finding evidence for effective population growth or for complex fluctuations that have been correlated with major events in human history such as the introduction of antibiotic treatment. 3) The strength of purifying selection was estimated with a simulation based approach, finding a genome-wide selection coefficient several order of magnitude higher compared to other prokaryotes and eukaryotes (Pepperell et al. 2013).

While some of these results might be biased by unaccounted population structure (Heller et al. 2013) or sampling biases (Lapierre et al. 2016), potentially they are all impacted by the violation of the Kingman’s assumption described above, and their conclusions could be affected by model misspecification (Tellier and Lemaire 2014).

Given the undergoing efforts in controlling and stopping the spread of tuberculosis, and the global impact of this pathogen that causes more than 1.4 million deaths each year (WHO 2019), it is important to evaluate the adequacy of the population genetic models used to study tuberculosis epidemics. To this end, we considered eleven MTB whole genome sequence (WGS) data sets, and used an Approximate Bayesian Computation (ABC) approach based on simulations to find the best fitting model among Kingman’s coalescent, and two MMC models, the Beta coalescent (Schweinsberg 2003) and the Dirac coalescent (Eldon and Wakeley 2006). We found that MMC were the best fitting model for ten of the eleven data sets (nine fitted best to the Beta, one to the Dirac coalescent). In addition, we investigated the consequences of violating the assumption on the offspring distribution when performing demographic inference with the BSP, and found that it leads to the inference of false population dynamics. Consequently, demographic inference based on models assuming non-skewed offspring distribution (i.e. Kingman’s coalescent) likely leads to inaccurate results when applied to MTB epidemics, and potentially to the epidemics of other pathogens with similar life histories.

## Results

### Models and data sets

MTB is thought to be strictly clonal, with lateral gene flow completely absent, or very rare (Hershberg et al. 2008, Gagneux 2018, Chiner-Oms et al. 2019). Therefore, the MTB genome can be considered as a single genetic locus, and one single genealogy describes the relationships among all MTB strains in any data set. The shape of the genealogy of a sample is influenced by many factors, such as the underlying offspring distribution, sampling scheme, population subdivision, geographic population structure, migration and changes in population size. To avoid these confounding effects, we considered only populations that were unlikely to be affected by population structure, sampling biases, population subdivision and migration. We searched the literature for whole genome sequence (WGS) data sets of MTB where all strains were sampled from a single phylogenetic clade that was restricted to a particular geographic region, and identified eleven studies. Most of these data sets represent single outbreaks (Methods). For each data set, we downloaded the raw Illumina sequences (Sup. Table 1) and used a bioinformatic pipeline described in the Methods to identify high confidence SNPs (Table 1). To test the robustness of our analyses to different SNP call procedures, we performed an additional SNP call altering one key parameter: the minimum proportion of reads supporting a SNP call (from 90% to 75%, see Methods). We found that the allele frequency spectrum (one of the most important statistics, see below) was robust to the different SNP call settings (Sup. Figs. 1-2). We performed the main analyses (see below) on both data set variants. Since the results were similar, and we consider the SNP call with the 75% threshold less stringent, in the manuscript we report the results for the data sets based on the 90% threshold. The results for the data sets based on the 75% threshold and the comparison between the two different sets are reported in Sup. Table 2.

**Table 1.** Data sets used in this study

| Data set<sup>1</sup> | Number of strains | Number of polymorphic positions | Locality of sampling |
| --- | --- | --- | --- |
| Eldholm 2015 | 248 | 497 | Buenos Aires (Argentina) |
| Lee 2015 | 147 | 454 | Nunavit (Canada) |
| Stucki 2016 | 175 | 6264 | Central African countries |
| Shitikov 2017 | 176 | 1164 | Russia and Belarus |
| Roetzer 2013 | 61 | 74 | Hamburg (Germany) |
| Comas 2015 | 21 | 1334 | Ethiopia |
| Bainomugisa 2018 | 81 | 401 | Daru Island (PNG) |
| Bjorn-Mortensen 2016 | 121 | 128 | East Greenland |
| Folkvardsen 2017 | 702 | 214 | Copenhagen (Denmark) |
| Stucki 2014 | 60 | 128 | Bern (Switzerland) |
| Eldholm 2016 | 25 | 17 | Oslo (Norway) |

<sup>1</sup> We identified the data sets with the first author's name and year of the original publication

Excluding population structure, two factors that can shape the diversity of these data sets are changes in population size, and whether offspring distributions are skewed. We modeled changes in population size assuming exponential population growth, as has often been done in previous studies (Eldholm et al. 2015, Merker et al. 2015, Eldholm et al. 2016, O’Neill et al. 2019).

We modeled skewed offspring distributions with two MMC models deriving from explicit population models: 1) the Beta coalescent, in which the probability of each individual to coalesce in a multiple merger event is regulated by a Beta distribution with parameters α (between 1 and 2) and 2-α. The Beta coalescent was originally introduced to model populations with sweepstakes reproduction (Schweinsberg 2003), but it was also proposed to capture the genealogies of populations undergoing recurrent bottlenecks and of epidemics characterized by super-spreaders (Tellier and Lemaire 2014, Hoscheit and Pybus 2019). Lower values of α (closer to one) correspond to larger multiple mergers events, and for α=1 the Beta coalescent corresponds to the Bolthausen-Sznitman (BSZ) coalescent. The BSZ coalescent is an explicit model for genealogies of populations evolving under rapid positive selection, which lead certain families of selected genotypes to have strongly increased sizes compared to the average (Bolthausen and Sznitman 1998, Brunet and Derrida 2013, Neher and Hallatschek 2013).

2) The Dirac coalescent, also known as psi coalescent, is defined by a single parameter (ψ). The parameter ψ represents the average proportion of sampled lineages that coalesce in a single multiple merger event. The Dirac coalescent was derived from a modified Moran population model, where at each generation a proportion of the population is replaced by the offspring of a single individual, thus also modeling skewed offspring distribution (Eldon and Wakeley 2006).

Importantly, none of these MMC models was derived from a population model specific for MTB. Nevertheless, they are useful to investigate whether processes leading to skewed offspring distribution (on the coalescent time scale) are important in MTB, and we will discuss this further below. In a first analysis, we tested whether modeling skewed offspring distributions alone explained the observed genetic diversity better than modeling variable population sizes (with an exponential growth model) and standard offspring distributions. Therefore, we considered MMC models with constant population sizes. It was previously shown that even for a single locus, these hypotheses can be I distinguished for moderate sample sizes and high enough mutation rates (Eldon et al. 2015, Freund and Siri-Jégousse 2019).

Subsequently, we explored whether modeling skewed offspring distribution together with variable population size (exponential growth) further improved the fit to the data.

### Model selection and parameter estimation with Approximate Bayesian Computation

For model selection and parameter estimation, we used an ABC approach based on random forests (RF), as reported in detail in the Methods section and represented in Figure 1. We considered four models: Kingman’s coalescent with constant population size (KM), Kingman’s coalescent with exponential population growth (KM+exp), Beta coalescent with constant population size (BETA), and Dirac coalescent with constant population size (Dirac). Briefly, for each data set, we collected the SNPs identified with the bioinformatic analysis, reconstructed the genotype of the most recent ancestor (MRCA) and used it to polarize the SNPs. We then calculated a set of 24 summary statistics measuring genetic diversity and phylogenetic properties. For each model, we performed 125,000 simulations of a sample of size n, where n is the number of individuals in the data set, drawing the scaled population size from a prior distribution spanning one order of magnitude around the Watterson estimator (θ_obs_). As described in Pudlo et al. (2015), we performed model selection via ABC using a random forest of 1,000 decision trees. For parameter estimation within a model class, we followed the approach of Raynal et al. (2018).

**Figure 1.**
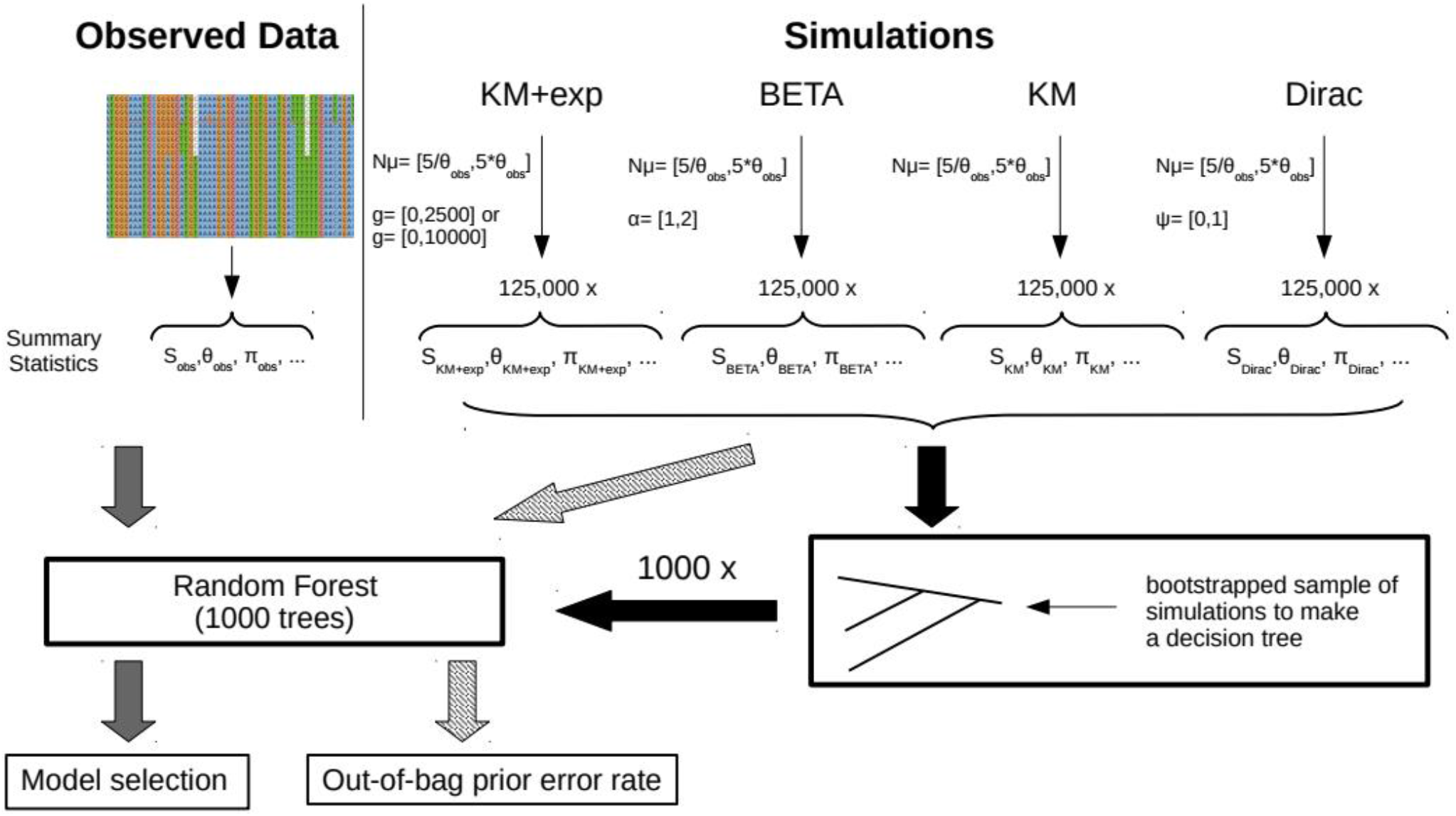
Workflow of the main ABC-RF analysis (model selection), see text and Methods for details.

We found that for most data sets, the ABC approach had overall good discriminatory power, with out-of-bag (OOB) error rates (the misclassification probabilities, see Methods and Table 2 for details) ranging from 4% to 16.4%. The only exception was the data set Eldholm 2016 (OOB error rate = 32.2%.), which was the data set with the lowest genetic diversity. Most importantly for our study, the probability that data generated under a model with standard offspring distribution (KM and KM+exp) was misclassified as multiple merger was low (1.1% – 7%), again the only exception was the data set Eldholm 2016 (18%).

**Table 2.** Results of model selection and parameter estimation

| Data set | Selected Model | OOB error rate (misclassification % as MMC)<sup>1</sup> | Posterior Probability | Median and 95% posterior credibility interval of coalescent parameters<sup>2</sup> |
| --- | --- | --- | --- | --- |
| Eldholm 2015 | BETA | 5.4% (2.1%) | 96.6% | $\alpha$: 1.19 (1.01 – 1.39) |
| Lee 2015 | BETA | 6.8% (2.6%) | 96.0% | $\alpha$: 1.27 (1.06 – 1.59) |
| Stucki 2016 | KM+exp | 4.1% (1.1%) | 100% | g: 2828 (676 - 4508) |
| Shitikov 2017 | KM+exp | 5.3% (1.8%) | 99.7% | g: 2833 (1158 – 4819) |
| Roetzer 2013 | BETA | 15.8% (7.0%) | 95.7% | $\alpha$: 1.23 (1.02 – 1.50) |
| Comas 2015 | BETA | 16.4% (5.4%) | 87.3% | $\alpha$: 1.31 (1.05 – 1.72) |
| Bainomugisa 2018 | BETA | 8.9% (3.3%) | 97.6% | $\alpha$: 1.22 (1.02 – 1.51) |
| Bjorn-Mortensen 2016 | BETA | 10.2% (4.2%) | 80.6% | $\alpha$: 1.03 (1.00 – 1.16) |
| Folkvardsen 2017 | BETA | 4.0% (1.4%) | 97.8% | $\alpha$: 1.13 (1.02-1.33) |
| Stucki 2015 | KM+exp | 13.2% (5.4%) | 87.4% | g: 13288 (3756 – 19932) |
| Eldholm 2016 | Dirac | 32.2% (18.0%) | 77.1% | $\psi$: 0.36 (0.19 – 0.61) |

<sup>1</sup> The out-of-bag error rate is the probability that a simulation is misclassified as coming from any other model class, between parentheses we report the probability that a simulation generated with KM or KM+exp is missclassified as a MMC (BETA or Dirac).
<sup>2</sup> The interval between the 0.025 quantile and the 0.975 quantile of the parameter of the selected model 255 (g for KM+exp, $\psi$ for Dirac and $\alpha$ for BETA). The growth rate g is reported as used in Hudson's ms (for diploid genealogies), thus all growth estimates have to be halved to be interpreted for MTB.

We found that BETA was the best fitting model for seven of the eleven data sets, KM+exp was the best model for three data sets, and Dirac was the best model for one data set. For all but one data set (Eldholm 2016), the posterior probability of the selected model was higher than 80% and therefore more than four times more likely than the second best fitting model (Table 2, Sup. Table 2).

One potential problem when performing model selection, is that none of the considered models is able to generate key features of the observed data (i.e. the considered models are not adequate; Gelman et al. 2013). To exclude this possibility, we performed posterior predictive checks, in which for each data set, we simulated data under the best fitting model using the median of the posterior distribution of the relative parameter. We then compared the observed data with the simulated data. If the selected model is adequate, we expect the simulated and observed data to be similar. Conversely, if the selected model is not adequate, we expect simulated and observed data to be different. We found that for all but two data sets, the observed values of 20 summary statistics were within the range of values obtained from the simulated data, indicating that the best model can reproduce the features of the observed data (Sup. Figs. 3-13). The two exceptions were Stucki 2016 and Shitikov 2017, for which respectively the 0.9 quantile of the Hamming distance, and the mean of the minimal observable clade size statistic were not overlapping with the simulated values (Sup. Figs. 12-13). This indicates that the best fitting model (KM+exp) cannot reproduce the observed data, and that none of the considered models is adequate for these two data sets.

### Hidden population structure and population decline in the data set Lee 2015

In our analysis, we focused on local data sets to control for the confounding effect of complex population dynamics and population structure. However, in one case (Lee 2015), it is possible that some degree of population structure was still present. Lee 2015 is a data set sampled from an epidemic in Inuit villages in Nunavik, Quebec, Canada (Lee et al. 2015b). Lee et al. (2015b) showed that transmission of MTB among patients was more frequent within a village than between villages, and that related strains tended to be present in the same village. This was supported by the reconstructed phylogenetic tree, which showed three clades separating at the root that could represent distinct subpopulations (Sup. Fig. 14; see also Fig. 2 in Lee et al. 2015b). This data suggests the existence of some degree of geographic population structure. Therefore, we tested whether this might influence the results of our model selection. To do this, we ran two analyses: 1) we repeated the ABC-RF analysis on three subsets of Lee 2015, which represent the three main clades described above (Sup. Fig. 14). Under the assumptions that the separate branches of the phylogeny reflect different sub-populations, and that migration does not alter the coalescent rates within the sub-populations, the genealogy of each subclade should then follow one of the coalescent models that we are fitting. We found that BETA was the best fitting model for two of the sub-clades, while Dirac was the best fitting model for the third (Table 3). The posterior predictive checks showed that the best model could reproduce the data of these three subsets (Sup. Figs. 15-17). However, the posterior probabilities were low compared to the complete data set, and the misclassification probabilities were larger. This was probably due to the smaller sample size of the individual subsets compared to the full data set (Table 3). 2) We performed an additional model selection analysis between three competing models, BETA, Dirac and a third scenario, in which we modeled a structured population with migration and with standard offspring distribution and exponential growth (KM+exp). Also in this case, BETA resulted to be the most likely model (Table 3, see Methods for details). Overall, our findings indicate that it is unlikely that the MMC signal in the Nunavik MTB population is an artifact caused by population structure.

**Figure 2.**
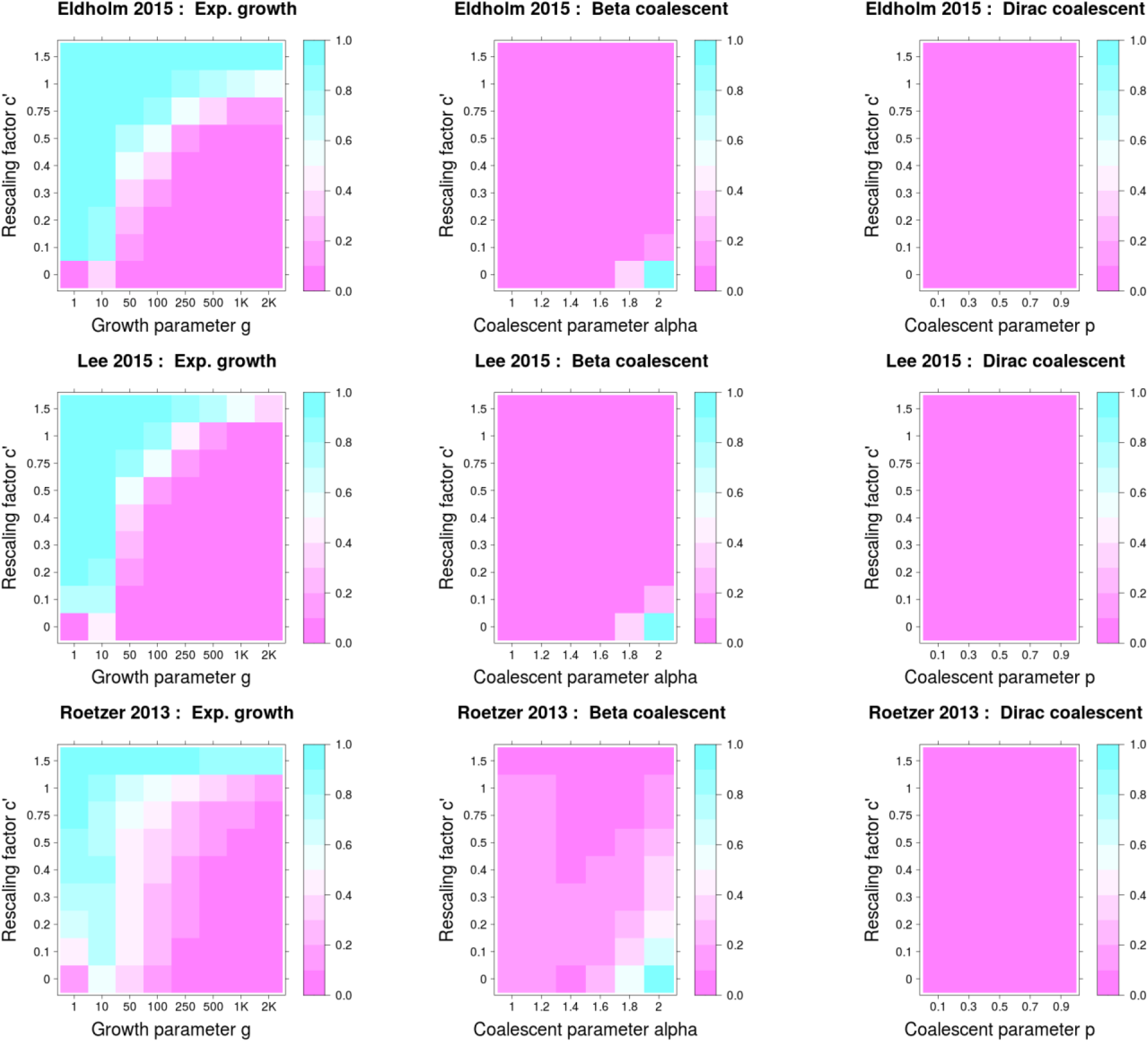
Proportion of model misidentification for serial simulations when model selection was performed via ABC using ultrametric tree models. Misclassification probabilities are shown as a function of c’ (the proportion of the genealogy corresponding to the time period in which samples are collected. i,e the period of sampling spans a time period c’oh, where h is the expected height of the genealogy without serial sampling), and of the parameter of the coalescent models. Misclassification was measured as follow: i) for simulations from serially sampled Kingman’s coalescent with exponential growth as being misidentified as either Beta or Dirac (first column) ii) for simulations from serially sampled Beta or Dirac coalescents as being misidentified as Kingman’s coalescent with or without exponential growth (second and third columns).

**Table 3.** Results of model selection for the complete Lee 2015 data set, and for the three major sub-clades separately. The shaded row represents the results of the standard analysis on the full data set.

| Data set | N. of strains | Selected Model | OOB error rate (misclassification % as MMC)<sup>1</sup> | Posterior Probability | Second best fitting model |
| --- | --- | --- | --- | --- | --- |
| Lee 2015 | 147 | BETA | 6.8% (2.6%) | 96.0% | Dirac |
| Lee 2015 Clade A | 61 | Dirac | 20.2% (10.0%) | 82.2% | BETA |
| Lee 2015 Clade B | 36 | BETA | 16.6% (6.8%) | 64.4% | KM |
| Lee 2015 Clade C | 49 | BETA | 12.8% (5.0%) | 78.6% | Dirac |
| Lee 2015 Pop. Structure<sup>2</sup> | 147 | BETA | 8.5% (6.9%) | 94.4% | Dirac |
| Lee 2015 Pop. Decline<sup>3</sup> | 147 | BETA | 3.4 % (3.0 %) | 99.9% | Pop. decline |

<sup>2</sup> Model selection among BETA, Dirac, and KM with structure
<sup>3</sup> Model selection among BETA and KM with population decline.

Structured populations have similar genealogies to populations that are shrinking in size (forward in time), with many lineages coalescing close to the root. In their original publication, Lee et al. (2015b) used the Bayesian Skyline Plot (Drummond et al. 2005) to reconstruct the fluctuations in population size of the Nunavik population, and found evidence for a slow population decline. Here, we are not interested in whether the inferred population decline is genuine or caused by unaccounted population structure, we only want to assess whether a decline in population size could bias our analysis. We repeated the ABC-RF model selection among two models: BETA and KM with population decline (see Methods for details). Again, we found that BETA was the best fitting model (Table 3), thus indicating that our results for this data set are unlikely to be an artifact caused by population decline.

### Serial sampling

One limitation of our analysis is that it assumes that all samples were collected at the same time (synchronous sampling). Generally, MTB strains are sampled from the sputum of patients, which is collected when they first present for diagnosis. All data sets that resulted in a MMC as best fitting model included samples obtained over extended periods of time (serial sampling), corresponding to between ~8% and ~100% of the estimated tree age (Sup. Table 3).

We investigated whether, at least in principle, the violation of the assumption of synchronous sampling could bias the results of the ABC analysis performed above, and whether the better fit of MMC could be an artifact due to such violation. To do this, we generated simulated data assuming serial sampling, and performed model selection on the simulated data assuming synchronous sampling (see Methods). Since this analysis depends on assumptions about the sample size, the genetic diversity, and the sampling times, we used the settings (sample size, observed generalized Watterson’s estimator as scaled mutation rate, and the real years of isolation) of three of the observed data sets, which differed in these characteristics (Eldholm 2015, Lee 2015 and Roetzer 2013).

We found that data simulated under KM+exp can be misclassified as BETA or Dirac if we do not account for serial sampling. Specifically, this was true for extended sampling periods compared to the expected height of the genealogy (on the coalescent time scale), and for low growth rates (Fig. 2). Similarly to model selection, not accounting for serial sampling affected the estimation of the growth rate parameter, and this effect was greater for large sampling periods and low growth rates (Sup. Fig. 18).

It is difficult to relate these results to the observed data sets because we do not know the scaling factor between coalescent time and real time (and therefore cannot estimate the value of c in Fig. 2, see Methods). However, for six of the eight data sets that resulted in a MMC as best fitting model, we estimated large growth rates (g ≥ 1,000) under the KM+exp model (Sup. Table 2), indicating that serial sampling is unlikely to affect the results of model selection in these cases (Fig. 2, Sup. Fig. 18). Nevertheless, we adopted a complementary approach, in which we drastically reduced serial sampling by sub-sampling only strains that were isolated in a single year. Since small data sets have lower discriminatory power, for this analysis, we selected the four data sets with the highest genetic diversity (data sets with more than 200 polymorphic positions: Eldholm 2015, Lee 2015, Folkvardsen 2017 and Bainomugisa 2018) among the ones for which the sampling times were available. For each data set, we repeated the ABC analysis on the largest possible subset of strains that were sampled in a single year (Sup. Table 1, Table 4). We found that all subsets had lower posterior probabilities and higher misclassification errors compared to the full data sets, most likely because of the smaller sample size. KM+exp was the best fitting model for two subsets, Dirac and BETA were the best fitting model for one subset each. For the data sets Eldholm 2015 and Folkvardsen 2017, the second and third most sampled years had a similar number of strains compared to the most sampled year. Therefore, we extended the analysis to these four additional subsets, which all resulted in BETA as the best fitting model (Table 4). We also noticed that in the subset of Lee 2015 (sampled 2012), all strains but one belonged to one of the three clades discussed above (clade A; Sup. Fig. 14). We suspected that this analysis was influenced by population structure and we repeated it excluding the single strain not belonging to clade A. Again, we found that Dirac was the best fitting model (Table 4, Sup. Table 2). We performed posterior predictive checks for all subsets and found that in all cases, the best fitting model could reproduce the observed data (Sup. Figs. 19-27).

**Table 4.** Results of model selection for the temporal subsets. Shaded rows contain the results for the full data sets.

| Data set<sup>1</sup> | N. of strains | Selected Model | OOB error rate (misclassification % as MMC)<sup>2</sup> | Posterior Probability | Second Best fitting model |
| --- | --- | --- | --- | --- | --- |
| Eldholm 2015 | 248 | BETA | 5.4% (2.1%.) | 96.6% | KM+exp |
| Eldholm 2015 (1998) | 34 | KM+exp | 20.9% (9.5%) | 75.9% | BETA |
| Eldholm 2015 (2001) | 31 | BETA | 18.3% (7.7%) | 79.9% | Dirac |
| Eldholm 2015 (2003) | 32 | BETA | 18.3% (7.8%) | 83.0% | KM+exp |
| Lee 2015 | 147 | BETA | 6.8% (2.6%) | 96.0% | Dirac |
| Lee 2015 (2012) | 45 | Dirac | 15.8% (6.5%) | 96.3% | BETA |
| Lee 2015 (2012) Clade A | 44 | Dirac | 22.7% (11.3%) | 84.5% | BETA |
| Bainomugisa 2018 | 81 | BETA | 8.9% (3.3%) | 97.6% | KM+exp |
| Bainomugisa 2018 (2014) | 56 | BETA | 11.3% (4.2%) | 94.9% | KM+exp |
| Folkvardsen 2017 | 702 | BETA | 4.0% (1.4%) | 97.8% | KM +exp |
| Folkvardsen 2017 (2009) | 53 | BETA | 14.0% (5.8%) | 85.1% | KM+exp |
| Folkvardsen 2017 (2010) | 64 | KM+exp | 13.5% (5.7%) | 83.8% | BETA |
| Folkvardsen 2017 (2012) | 52 | BETA | 15.1 % (6.4%) | 96.8% | KM |

<sup>1</sup> Between parentheses we report the year in which the strains were sampled (only for temporal subsets)
<sup>2</sup> The out-of-bag error rate is the probability that a simulation is misclassified as coming from any other model class, between parenthesis we report the probability that a simulation generated with KM or KM+exp was misclassified as MMC (BETA or Dirac)

Overall, these findings indicate that not accounting for serial sampling can indeed bias the results of model selection in favor of MMC models. However, this was unlikely to affect datasets with large growth rates (six out of eight). Additionally, seven of the nine subsets in which we minimized the serial sampling to one single year resulted in a MMC as best fitting model.

### Sensitivity to the choice of prior distributions

An important aspect of Bayesian analyses is to test whether the results are robust to different priors and model assumptions. Therefore, we performed a set of analyses investigating the sensitivity of the results of the ABC to changes to the prior distribution of the growth rate (g) for KM+exp, of the parameter α for BETA, and of the scaled mutation rate (θ) for all models. These analyses are reported in Appendix 1. Overall, we performed four additional ABC analyses testing different prior combinations, and for two of them we also tested the datasets obtained with the 75% threshold on the SNP call (Sup. Table 2, Sup. Figs. 28-35). We found that in 89% of cases, the results of model selection did not change compared to the main analysis presented above. The datasets that were sensitive to the prior choice were mostly the ones with low sample sizes, low number of polymorphisms, low posterior probabilities and large error rates, further highlighting that the results of small data sets should be taken with some caution.

### Modeling skewed offspring distribution and variable population size

So far, we considered only multiple merger models with constant effective population size. However, for many of the outbreak analyzed, it is reasonable to expect that the effective population size was growing (forward in time). We therefore performed a further model selection between the best fitting model resulted from the analysis described above, and two additional model classes: BETA with exponential population growth (BETA+exp), and Dirac with exponential population growth (Dirac+exp; see Methods for details).

Overall, we found that 22 of the 23 data sets (including subsets) resulted in a MMC (with or without growth) as best fitting model. For 11 data sets we found that the best fitting model was a MMC with exponential growth, suggesting that these populations were indeed growing in size (Sup. Table 2). However, for some data sets the posterior probabilities were low, indicating that different models fitted the data similarly well.

### Skewed offspring distribution can bias demographic inference with the Bayesian Skyline plot

To reconstruct the past demographic history of MTB and other organisms, many studies use nonparametric approaches such as the Bayesian Skyline Plot (BSP; Heled and Drummond 2008). In the last few years (since 2013), at least 16 studies applied the BSP to MTB data sets (cited in the Introduction). It was shown before that the BSP can be biased by unaccounted population structure (Heller et al. 2013), recombination, and non-random sampling (Lapierre et al. 2016). Hence, we next assessed the impact of skewed offspring distribution on demographic reconstruction with the BSP. To do this, we simulated 50 data sets under the BETA coalescent with constant population size. We used different values of α corresponding to the range of values estimated for the observed MTB data sets (α = 0.5, 0.75, 1, 1.25, 1.5; 10 replicates each, see Methods). We then performed an extended Bayesian skyline plot analysis on the simulated data using BEAST2 (Bouckaert et al. 2019). For 49 of the 50 simulated data sets we found that the 95% Highest Posterior Distribution interval of the number of population size changes did not include zero, thus rejecting the constant population size model (Heled and Drummond 2008). The inferred skyline plots showed different patterns of fluctuation of the effective population size (Fig. 3, Sup. Figs. 36-40). These results demonstrate that skewed offspring distribution alone can bias the outcome of the BSP, leading to the inference of complex population dynamics that are entirely due to the violation of the assumption on the offspring distribution.

**Figure 3.**
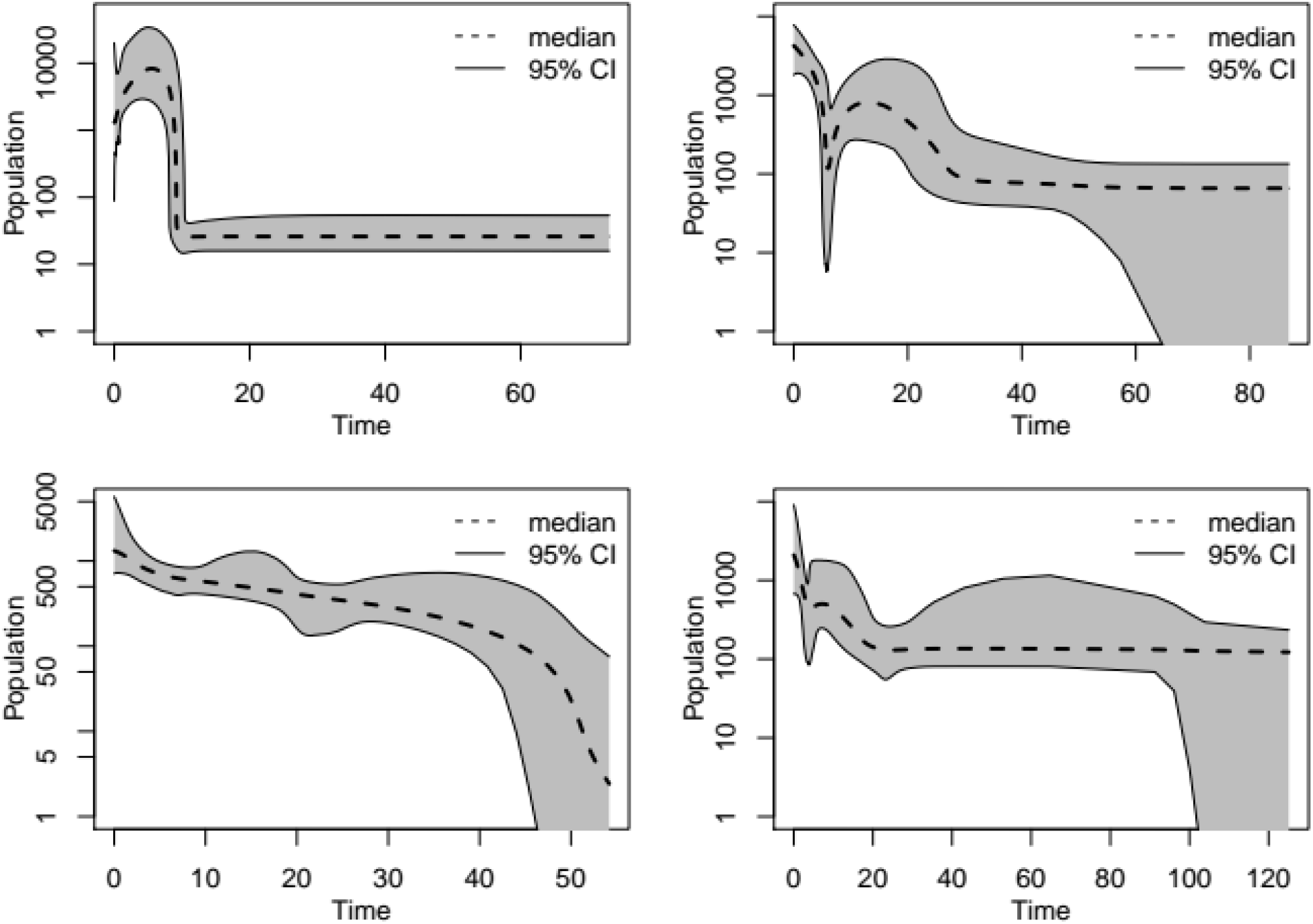
Four examples of Bayesian skyline plot obtained from data simulated under a Beta coalescent with constant effective population size, and α = 1 (top-left), α = 1.25 (top-right and bottomleft), and α = 1.5 (bottom-right). On the y axis the inferred effective population size, on the x axis the time in years before sampling. The plots for all 50 simulations are reported in Sup. Figs. 36-40.

## Discussion

As for many other organisms, demographic inference based on Kingman’s coalescent has become an important tool to study the evolution and epidemiology of MTB. As always when performing statistical inference, the results of these analyses depend on the assumptions of the model.

Our results showed that, when studying MTB local population and outbreaks, models that do not allow for skewed offspring distribution on the coalescent timescale (Kingman) had consistently worse fit compared to MMC, and can lead to the inference of false fluctuations of past effective population size.

### Better fit of MMC models is robust to possible confounders

We tested the robustness of our results by changing different aspects of the analysis, including one of the main SNP call parameters and the choice of prior distributions in the ABC (Appendix 1). We performed eight different model selection analyses on all 23 data sets (including subsets), and invariably found that a large majority of the data sets resulted in a MMC as best fitting model (Sup. Table 2). When we included MMC models with exponential growth in an additional ABC analysis, only one data set (Stucki 2016) resulted in Kingman as the best coalescent type. However, for this data set, the posterior predictive checks could not reproduce the observed genetic diversity, indicating that none of the models used were adequate (Sup. Fig. 13). While our results are robust to different settings, it is important to discuss some of the assumptions on which they are based, in particular regarding 1) sampling bias, 2) population structure and 3) serial sampling.

1. One factor that could bias the results of demographic inference is sampling bias (Lapierre et al. 2016). The majority of the data sets considered in this study are composed of strains sampled from (nearly) all known TB cases caused by a certain phylogenetic clade (Roetzer et al. 2013, Lee et al. 2015b, Stucki et al. 2015, Bjorn-Mortensen et al. 2016, Eldholm et al. 2016), or by a random subset of them (Folkvardsen et al. 2017). Therefore, these data sets should not be strongly affected by sampling bias.
2. In our analyses, we assumed that the data originated from single panmictic populations. For clonal organisms, or when only a single locus is available, recognizing population structure from genetic data is challenging, because clusters of genetically similar individuals can occur also in a panmictic population (at a single locus). Therefore, to avoid the effect of unaccounted population structure, we chose data sets from single outbreaks and local populations in restricted geographic regions. However, we cannot completely exclude that for some data sets, some level of geographic structure was present, and this might have affected the results of our analyses. For one data set where a prior analysis suggested some degree of population structure (Lee et al. 2015b), we found that including a model with population subdivision and migration, or sub-sampling the potential sub-populations, resulted again in a MMC as the best fitting model, indicating that population structure is unlikely to bias the results of this analysis.
3. Serial sampling could also affect the results of model selection. All the considered models assume a common sampling time for all strains, which is almost never the case for MTB data sets. Due to the relatively short generation time of MTB compared to the sampling period, this may likely correspond to serial sampling on the coalescent time scale. Under serial sampling on the coalescent time scale, our simulations of several scenarios mimicking three of our data sets revealed that Kingman’s genealogies under exponential growth can be misidentified as MMC, while inferring a true MMC is not affected. The misclassification probability was higher when the sampling window spanned a large part of the genealogical history of the sample, but dropped considerably under strong exponential growth (Fig. 2, Sup. Figs. 18, 28-29, Appendix 1).

For six of the eight full data sets that resulted in a MMC as best model, the fitted growth parameter under KM+exp was 1,000 or higher, and therefore, it is unlikely that the serial sampling influenced the results of model selection for these data sets. Additionally, when we sub-sampled strains from a single year from four data sets, thus minimizing the effect of serial sampling, seven of the nine subsets resulted in a MMC model (Table 4).

For most data sets the better fit of MMC is unlikely to be an artifact caused by serial sampling. Nevertheless, it is possible that serial sampling influenced some of these results. Obtaining truly synchronous samples is essentially impossible for MTB, and reducing the sampling time to one year or few months could still not be enough to avoid serial sampling on the coalescent time scale.

Thus, to overcome the limitation of assuming synchronous sampling, we encourage future studies to develop MMC models that explicitly consider the time of sampling. Such a model is proposed in Hoscheit and Pybus (2019) as an extension of the Beta coalescent, but without an explicit mechanism to convert real time units into coalescent time units.

Our results, that the Beta coalescent is generally fitting better to outbreaks of MTB, is in contrast with the use of Dirac coalescent as the model for within-host MTB genealogies in Morales-Arce et al (2020). While within-host and between-host dynamics are surely different, it would be interesting to test whether the Beta coalescent also fits better within-host data. A hint that this could be the case is that the strength of multiple-mergers (the coalescent parameter ψ) estimated in Morale-Arce et al. (2020; Figure 3) was very low.

### What are the biological processes generating multiple merger genealogies in MTB?

At least three different biological processes could lead to MMC genealogies in MTB: 1) repeated bottlenecks at transmission between hosts, 2) super-spreaders, and 3) rapid recurrent selection induced by the immune system, and/or by antibiotic treatment.

To formally test which of these three processes (if any) is generating the MMC genealogies in outbreaks of MTB, we would need to have an explicit population model for each of these scenarios and for their combinations. Additionally, for each population model, we would need to know the corresponding coalescent process. If these three mechanisms (or their combinations) lead to different coalescent processes, we could then use the observed data to test the different hypotheses.

However this information is only partly available:

1. Although the Beta coalescent has been proposed as genealogy model for populations with recurrent strong bottlenecks (Tellier and Lemaire 2014), rigorous mathematical modeling predicted different coalescent processes for extreme bottlenecks, which allow for simultaneous multiple mergers (Birkner et al. 2009, Casanova et al. 2019). In any case, these models are not appropriate for MTB outbreaks, because they model bottlenecks affecting the whole population. Transmission bottlenecks in a MTB population can be thought as multiple serial founder events. To model such processes, one would need to account for both intra-host dynamics within multiple hosts, and between-hosts transmission. Such a model is currently not available, and it is not clear to which coalescent process it would lead. It is possible that the Beta or the Dirac coalescent capture well the main features of the genealogies generated by such a mechanism. However, without an explicit model a formal test is impossible.
2. The Beta coalescent has also been proposed to model the genealogies of epidemics with superspreaders. A simulation study showed that the parameter α contains information about the degree of super-spreading, with lower values of α corresponding to higher level of super-spreading (and larger multiple merger events, Hoscheit and Pybus 2019). However, also in this case, an explicit population model is missing. Thus, we cannot formally test whether MMC genealogies in MTB outbreaks are due to super-spreaders.
3. The genealogies of populations evolving under recurrent rapid positive selection are described by the Bolthausen-Sznitman coalescent (BSZ; Bolthausen and Sznitman 1998, Brunet and Derrida 2013, Neher and Hallatschek 2013), which correspond to the Beta coalescent with α=1. Therefore, by estimating the value of α from the data, we can partially test whether rapid selection is the single factor causing the MMC signal in MTB outbreaks. More precisely, if the 95% credibility interval (CI) of the posterior distribution of α does not include one, we can exclude that rapid selection is the only factor involved.

For five of the data sets that resulted in a MMC as the best fitting model, the 95% CI of α did not include one (Sup. Figs. 33-35). These results suggest that for these data sets, we can exclude recurrent rapid positive selection as the only process leading to multiple merger genealogies, while for many others, the data fitted well with the BSZ. For instance, two data sets, Eldholm 2015 and Bainomugisa 2018, represent outbreaks of drug resistant clones which acquired several additional drug resistance mutations during the outbreak (Eldholm et al. 2015, Bainomugisa et al. 2018). It is possible that strains with novel drug resistance mutations were advantaged, resulting in recurrent positive selection. This scenario would lead to the BSZ (corresponding to α=1), and indeed the estimates of α encompassed 1 for both data sets.

In this analysis, we used the Bolthausen-Sznitman coalescent as a null model for ongoing constant selection pressure, as also proposed in Neher and Hallatschek (2013). This coalescent model arises in several models of ongoing selection in a rapidly adapting fixed-size population, where the constant size is maintained by viability selection against the least fit individuals (Berestycki et al. 2013, Brunet and Derrida 2013, Neher and Hallatschek 2013). In these models, multiple mergers occur when an individual becomes much fitter than the rest of the population. However, we want to stress that there are alternative MMC models that correspond to different selection scenarios, such as a model with a modified fitness distribution (Huillet 2014), models considering the effect of several beneficial mutations (Desai et al 2013, Huillet 2014, Schweinsberg 2019), or hitchhiking of a neutral locus with a single beneficial mutant (Gillespie 2000).

Because of the lack of explicit models, we cannot test the exact biological process or processes generating multiple merger genealogies in MTB. The better fit of MMC models only indicates that one or more processes leading to skewed offspring distribution (on the coalescent time scale) play an important role in shaping the diversity of MTB populations. Moreover, our estimates of α, the inference of the best fitting MMC models (Beta or Dirac), and whether the best fitting model included an exponential growth component, differed distinctively across data sets. This suggests that different processes, or different magnitudes of the same process, produced multiple merger genealogies in different populations. For example, two of the data sets that had an estimate of α lower than 1 (corresponding to larger multiple merger events, see Appendix 1) were sampled from different outbreaks with important super-spreading events (Stucki 2015, Lee 2015 Clade A). Stucki 2015 is a data set representing an outbreak characterized by one key patient that had two distinct disease episodes, separated by three years, both of which resulted in a large number of secondary infections (Stucki et al. 2015). The data set Lee 2015 Clade A is a sample of an outbreak occurred in an Inuit village in Quebec (Canada), in this case two patients were responsible for 75% of all secondary infections (Lee et al. 2015a, Lee et al. 2020). This data strongly suggests that for these data sets, the mechanism generating large multiple mergers was the super-spreading behavior of one or few patients. We found only one additional data sets that resulted in α lower than 1, and interestingly it was also an outbreak among Inuit villages in East Greenland (Bjorn-Mortensen et al. 2016). Lee et al. (2015a, 2020) found that all secondary cases of the Canadian outbreak visited the same local community gathering houses. It is possible that such social gatherings played a similar role also in the spread of the Greenlandic outbreak leading to large multiple merger events.

An additional observation is that that the magnitude of multiple mergers might also change through time. For example, the temporal subsets of Folkvardsen 2017 fitted values of α around 1, or larger than one, depending on the year of sampling (Sup. Fig. 34).

Finally, additional biological processes might also generate multiple merger genealogies. MTB infections can remain latent for several years, and reactivate under favorable circumstances (e.g. immuno-suppression due to age, or HIV co-infection), although these cases might be less common than previously thought (Behr et al. 2018). It is known that stochastic exit from dormancy can lead to heavy-tailed offspring distributions, with bacteria exiting dormancy earlier having an extremely high reproductive success (Wright and Vetsigian 2019). Also in this case, mathematical modeling will be necessary to investigate whether this mechanism can affect the genealogies of MTB, and to identify the coalescent type that would result from such process.

### Modeling MTB genealogies

A population model for MTB should include host-to-host transmission, intra-host evolution, superspreaders, serial sampling, latency, population size changes and the potential selective pressure caused by the host immune system and/or by the antibiotic treatment; although some of these factors might not influence strongly the genetic diversity when modeled in combination with other mechanisms. This might very well result in a different multiple merger model compared to the ones that we employed. Such a model could also close the following implicit modeling gap in applying MMC to MTB and to bacteria in general. Mathematically, MMC processes have been introduced as approximations (with changed time scale) of the genealogy in underlying discrete population reproduction models, so called Cannings models (e.g. Möhle and Sagitov 2001). The underlying population models feature many offspring of a single individual per generation (e.g. Schweinsberg 2003, Eldon and Wakeley 2006, Desai et al. 2013). However, bacteria replicate through binary fission. While such population models are not applicable directly to bacteria, the underlying mathematical theory only needs to guarantee that the mergers within a single time point on the coalescent time scale follow a certain probability distribution, such that similar models can be defined, in which the large offspring number of one individual per generation is spread over multiple generations (Möhle and Sagitov 2001).

## Conclusions

In this study, we investigated whether the Kingman’s assumption of low variance of reproductive success is violated in MTB populations, and whether demographic inference with Kingman as null model could lead to artifacts due to model misspecification. We found that MTB genealogies are indeed affected by skewed offspring distribution, and that this can significantly bias the results of demographic inference, resulting in spurious past population dynamics. Potentially, these results can be extended to other obligate pathogens with similar life histories.

Further research is needed to develop an explicit population model for MTB. This would help to identify the biological mechanisms leading to multiple merger genealogies, and the most appropriate genealogy model for MTB populations. In the meantime, we encourage researchers to be extremely cautious when interpreting the results of demographic inference of MTB data sets based on the Kingman coalescent.

## Methods

### Data set selection

We searched the literature for WGS studies of outbreaks or local populations of *Mycobacterium tuberculosis.* We selected local data sets to avoid as much as possible geographic population structure and sampling biases that could influence the analysis. We identified 11 data sets: eight outbreaks and three clades with a restricted geographical range (the inferred phylogenies for all data sets are reported in Sup. Figs. 41-51):

*- Roetzer et al. 2013:* lineage 4 outbreak in Hamburg, Germany (61 strains, 74 polymorphic positions).
*- Comas et al. 2015:* lineage 7 strains sampled Ethiopia. Lineage 7 is a rare human adapted lineage endemic to Ethiopia and perhaps also to neighboring countries, only few genomes are available and most of them are included in this data set (21 strains, 1334 polymorphic positions).
*- Eldholm et al. 2015:* lineage 4 multi-drug resistant outbreak in Buenos Aires, Argentina (248 strains, 497 polymorphic positions).
*- Lee et al. 2015b:* lineage 4 outbreak in 11 Inuit villages in Nunavik, Québeq, Canada. We considered only the major sub-lineage Mj, a second smaller outbreaks of an unrelated sub-lineage (Mn) was excluded (147 strains, 454 polymorphic positions).
*- Stucki et al. 2015:* lineage 4 outbreak in Bern, Switzerland (60 strains, 128 polymorphic positions).
*- Bjorn-Mortensen et al. 2016:* lineage 4 outbreak in Greenland. To minimize the potential effect of population structure we considered only the major cluster GC4, because the other clusters represent independent outbreaks belonging to other sub-lineages (121 strains 128 polymorphic positions).
*- Stucki et al. 2016: s*ub-lineage L4.6.1/Uganda, belonging to lineage 4. This sub-lineage is endemic to central African countries (175 strains, 6264 polymorphic positions). *Eldholm et al. 2016:* lineage 2 outbreak in Oslo, Norway. From the data set of the original publication we excluded all strains that did not belong to the Oslo outbreak (25 strains, 17 polymorphic positions).
*- Folkvardsen et al. 2017:* large lineage 4 outbreak in Copenhagen, Denmark (702 strains 514 polymorphic positions).
*- Shitikov et al. 2017:* W148 outbreak belonging to lineage 2, this clade has also been named B, B0, CC2, East European 2 and ECDC0002 (176 strains, 1164 polymorphic positions).
*- Bainomugisa et al. 2018:* lineage 2 multi-drug resistant outbreak on a small island (Daru) in Papua New Guinea. From the data set of the original publication we excluded all the strains that did not belong to the Daru outbreak (81 strains, 401 polymorphic positions).

### Bioinformatic pipeline

For all samples Illumina reads were trimmed with Trimmomatic v0.33 (SLIDINGWINDOW: 5:20,ILLUMINACLIP:{adapter}:2:30:10) (Bolger 2014). Reads shorter than 20 bp were excluded for the downstream analysis. Overlapping paired-end reads were then merged with SeqPrep (overlap size = 15; https://github.com/jstjohn/SeqPrep). The resulting reads were mapped to the reconstructed MTB complex ancestral sequence (Comas et al. 2013) with BWA v0.7.12 (mem algorithm; Li and Durbin 2009). Duplicates reads were marked by the MarkDuplicates module of Picard v 2.1.1 (https://github.com/broadinstitute/picard). The RealignerTargetCreator and IndelRealigner modules of GATK v.3.4.0 (McKenna et al. 2010) were used to perform local realignment of reads around Indels. Reads with alignment score lower than (0.93*read_length)-(read_length*4*0.07)) were excluded: this corresponds to more than seven miss-matches per 100 bp.

SNPs were called with Samtools v1.2 mpileup (Li 2011) and VarScan v2.4.1 (Koboldt et al. 2012) using the following thresholds: minimum mapping quality of 20, minimum base quality at a position of 20, minimum read depth at a position of 7X minimum percentage of reads supporting a call 90%. Genomes were excluded if they had 1) an average coverage < 20x, 2) more than 50% of their SNPs excluded due to the strand bias filter, 3) more than 50% of their SNPs having a percentage of reads supporting the call between 10% and 90%, or 4) contained single nucleotide polymorphisms that belonged to different MTB lineages, as this indicates that a mix of genomes was sequenced. Because missing data can significantly impact population genetic inference we further excluded all strains that had less SNP calls than (average - (2 * standard deviation)) of the respective data set (calculated after all previous filtering steps). The filters described above were applied to all data sets with one exception: in the Comas 2015 data set most strains failed the strand bias filter, therefore this filter was not applied.

To test the robustness of our results, we performed an additional SNP call, in which we used a 75% threshold on the minimum proportion of reads supporting a call.

The single vcf were merged with the CombineVariant module of GATK v.3.4.0 (McKenna et al. 2010), the genotype field was edited to make it haploid (0/0 => 0; 1/1 => 1; 0/1 and 1/0 =>.). Vcftools 0.1.14 (Danecek et al. 2011) was used to extract variable positions excluding predefined repetitive regions (Comas et al. 2013) and excluding position with missing data.

The variable positions were converted in a multi fasta file including the reconstructed ancestral sequence on which the mapping was performed.

A phylogenetic tree based on the resulting variable positions was built with RaxML 8.2.11 (Stamatakis 2014) using a GTRCAT model and the -V option.

To identify the MRCA of each data set, the tree was rooted using the reconstructed ancestral sequence of the MTB complex as published in Comas et al. (2013), which is also the genome reference sequence used for the mapping. PAML4 (baseml; Yang 2007) was used to reconstruct the ancestral sequence of each data set. For all data sets the sequence accuracy (the marginal probability of the reconstructed sequence) of the MRCA was larger than 0.999.

For each data set all polymorphic positions for all strains and their reconstructed ancestor were then collected in fasta files. The data (obtained with both the 90% and 75% threshold) is available together with the ABC pipeline at https://github.com/fabianHOH/mmc_R_gendiv/tree/master/MTB_MMC_repo.

### Model selection and parameter estimation

For model selection and parameter estimation, we used a random forest based Approximate Bayesian Computation approach (Pudlo et al. 2015, Raynal et al. 2018).

We selected between Kingman’s n-coalescent (KM), Kingman’s n-coalescent with exponential growth (KM+exp), Beta coalescent (BETA) and Dirac coalescent (Dirac). For each data set we collected the genetic polymorphisms identified with the bioinformatic analysis and calculated a set of 24 summary statistics following the recommendations from Freund and Siri-Jégousse (2019), Scenario 3: the (.1,.2,.3,.4,.5,.6,.7,.8,.9) quantiles of the mutant allele frequency spectrum, the (.1,.3,.5,.7,.9) quantiles of the pairwise Hamming distances, the (.1,.3,.5,.7,.9) quantiles of the minimal observable clade sizes of each sequence, the number of segregating sites, the nucleotide diversity and the mean, standard deviation and harmonic mean of the minimal observable clade sizes. For each model we performed 125,000 simulations of a sample of size n where n is the number of individuals in the data set, drawing the scaled population size from a binomial distribution on log-equally spaced discrete θ spanning one order of magnitude around the Watterson estimator (θ_obs_), i.e. 11 steps in [θ_obs_/5,5θ_obs_], as in Freund and Siri-Jégousse (2019). The Watterson estimator is calculated as 2s/E(L), where s is the number of mutations observed in the data set and E(L) is the expected length the genealogy. For KM+exp we drew the value of the exponential growth rate (g) from a uniform distribution on [0.5,5000] except for the data sets Bainomugisa 2018, Bjorn-Mortensen 2016, Eldholm 2015, Stucki 2015, Lee 2015 (sampled in 2012) and Folkvardsen 2017, where we used a uniform distribution on [0.5,20000]. Note that this is a growth rate for a coalescent within a diploid population, values should be halved for interpretation in a haploid setting. The choice of wider ranges were based on preliminary analyses of the data with narrower prior distributions that showed a posterior distribution of g skewed at the upper end. For comparison, we also used an alternative setting with log-uniform priors on these ranges (Appendix 1). For BETA and Dirac we drew the value of the free parameters α and ψ from a uniform distribution on [1,2] and [0,1] respectively. Additionally, we performed an alternative analysis in which we drew the value of α from a uniform prior distribution on [0,2] (Appendix 1). Note that while BSZ is theoretically included in BETA as α=1, it will not be chosen as a parameter since we use a continuously distributed prior. To further assess whether BSZ is a well fitting model, we alternatively employed a spike and slab type prior, i.e. we replaced 1% of all parameters drawn from the continuous uniform prior for BETA with α =1. We used this in an additional ABC analysis on BETA with α from [0,2], the log prior on the growth rate for KM+exp, and the standard setting for Dirac (Appendix 1).

Simulations were performed in R as described in Freund and Siri-Jégousse (2019), the code is available at https://github.com/fabianHOH/mmc_R_gendiv.

As described in Pudlo et al. (2015), we performed model selection via Approximate Bayesian Computation using a random forest of decision trees, using the R package abcrf (Pudlo et al. 2015). We drew 1,000 bootstrap samples of size 100,000 from the simulations and then constructed decision trees based on decision nodes of the form S>t, where S is one of the summary statistics used. For each node, S and t are chosen so that the bootstrap sample is divided as well as possible in sets coming from the same of the four model classes (minimal Gini impurity). Nodes are added to the tree until all simulations of the bootstrap samples are sorted into sets from the same model class. Misclassification is measured by the out-of-bag (OOB) error, i.e. the proportion of decision trees for each simulation that sorts it into a wrong model class, averaged over simulations and, for the overall OOB error, model classes.

For parameter estimation within a model class, we followed Raynal et al. (2018). Here, the decision (regression) trees are constructed analogously, only S and t are chosen so that the parameters of the simulations have similar values in both sets divided by the node. This is achieved by minimizing the L^2^ loss, i.e. minimizing, for the two sets divided by the node, the L^2^ distances of the simulation parameter to the mean parameter in the set. Nodes are added until all simulations sorted into one leaf have the same parameter or there are less than 5 simulations allocated to the leaf.

The observed data is then assigned to the model class where the majority of decision trees for model selection assign it, and its posterior parameter distribution is given by the distribution of the weighted average parameter of the allocated leaf across all trees in the (regression) random forest (see Raynal et al. 2018, sections 2.3.2 and 2.3.3). The posterior probability for model selection is computed as a machine learning estimate of classifying the model class correctly, which includes another regression tree. See Pudlo et al. (2015) for details, a summary can be found in Appendix A.2 in Freund and Siri-Jégousse (2019).

### Misclassification probabilities

The misclassification probabilities were calculated as follow. After building the random forest, all simulated data sets were assigned to one of the models based on a random forest composed only of trees built from bootstrap samples not including this simulation (so that the data was not used to produce the decision trees). Since the true model is known, we can easily calculate the proportion of trees that classify this simulation in a wrong model class. The out-of-bag error rate (OOB) for a model class is the proportion of misclassified simulations over all simulations from the model class. The mean OOB error is the average of OOB errors across model classes. More informative error rates can also be calculated, for example the proportion of simulations from a bifurcating model that are classified as a multiple merger.

### Posterior predictive checks

To assess whether the best fitting model could reproduce the observed data, we performed posterior predictive checks. We simulated 10,000 sets of summary statistics under the best fitting model (using the median of the posterior growth rate or of the multiple merger coalescent parameter, obtained from the main analysis, Analysis 1 in Sup. Table 2) and compared them graphically with the value of the statistics observed in each data set. As scaled mutation rate, we used the generalized Watterson estimate 2s/E(L), where s is the number of mutations observed in the data set and E(L) the expected length of all branches for the best fitting coalescent model.

### Population structure and declining population size for the data set Lee 2015

To assess the effect of population structure in the data set Lee 2015, we simulated samples under Kingman’s n-coalescent with population structure. From the phylogenetic tree (Sup. Fig. 14), we identified four different clades with sizes 61, 36, 49 and 1. We then assumed these to be sampled from different sub-populations of equal size in an island model with scaled symmetric migration. We performed coalescent simulations under a structured (Kingman) coalescent with exponential growth. We used a discrete uniform prior on {0,2,4,.,.,5000} for growth rates and additionally drew the scaled migration rate m (in units of 4Nm*, where m* is the migration rate in the discrete island model) from the uniform discrete distribution {.25,.5,1,2,3}. We approximated Watterson’s estimator for a specific choice of parameters by replacing the expected total length of the coalescent by the mean total length from 10,000 coalescent simulations with these parameters. This approximation comes with an increased computational load compared to our standard approach, which in turn led us to the discretization of the prior described above.

For generating samples under Kingman’s n-coalescent with exponential decline, we had to slightly change the simulation procedure using ms. Since population decline may lead to coalescent times to large too simulate, we fixed the maximal population size in the past to 1,000 times the present population size. Then, given an exponential growth rate g<0, the decline starts at time log(1000)/(-g) (in coalescent time units backwards in time from time of sampling) and continues until the sampling time.

To compute Watterson’s estimator in this scenario for any g, we need the expected total length of the coalescent tree. Instead of computing it analytically, we recorded the total coalescent tree length of 10,000 simulations under the model and used their mean as an approximation of the expected total branch length.

As parameters for exponential decline, we use exponential growth rates drawn uniformly from {-250, 200,-150,-100,-50,-25,-10}, again we used a discrete prior distribution because Watterson’s estimator was too costly to approximate for a continuous range.

For both exponential decline and population structure, we ran the ABC-RF analysis as for all other data sets. Simulations were produced with Hudson’s ms as implemented in the R package phyclust.

### Accounting for serial sampling

Following Hoscheit and Pybus (2019), we add serial sampling to the MMC and to Kingman’s coalescent with exponential growth simply by stopping the coalescent at times (on the coalescent time scale) where further individuals are sampled. Then, we start a new (independent) coalescent tree that has rates and waiting times as the non-serial coalescent (multiple merger or with growth) started in the last state of the stopped coalescent plus adding one block with a single individual for each individual sampled at this time. A R implementation is available at https://github.com/fabianHOH/mmc_R_gendiv/tree/master/MTB_MMC_repo.

A problem with this approach is that one needs the scaling factor between coalescent time and real time. While estimation procedures coming from phylogenetics are available in the case of Kingman’s coalescent (e.g. Drummond and Rodrigo 2000), they cannot be applied directly to the case of multiple merger coalescents. Additionally, a brute force search for appropriate scaling on top of our models is computationally unfeasible with the ABC approach that we adopted in this study.

Hence, we assessed, for different fixed scaling factors, how strong the effect of ignoring serial sampling in the models is. We considered the setting of Eldholm 2015 (n=248, s=497, where n is the sample size and s is the number of mutations), Lee 2015 (n=147, s=454) and Roetzer 2013 (n=61, s=74). We used the real dates of the serial sampling for these data sets and we performed serial coalescent simulations as described above. We used different time (re)scaling factors c, such as c determines the time ct at which an individual sampled at real time -t (0 corresponds to the latest sampling time) is added as a new lineage to the coalescent tree (so ct is in coalescent time units). Here, we assessed c by setting the earliest sampling time (highest t) to a fraction c≥0 of the expected height of the coalescent tree if there was no serial sampling (so keeping all other parameters, but assuming c=0). For each c’ in {0,0.1,0.2,0.3,0.4,0.5,0.75,1,1.5}, we simulated 1,000 simulations under each parameter set (g in {1,10,50,100,250,500,1000,2000}, α in {1,1.2,1.4,1.6,1.8,2}, ψ in {0.1,0.3,0.5,0.7,0.9]) and then performed ABC model selection for each simulation (as described above), recording how often the serial coalescent simulations were sorted to which non-serial model class. We also reported the quality of parameter estimation for the growth rate or coalescent parameter by measuring the (absolute) distances of the estimated parameter to the parameter used for the simulation (Sup. Figs. 18, 29).

### Multiple-merger coalescents with exponential growth

We define Beta coalescents with exponential growth as limit processes of modified Moran models with variable population sizes, as described in Corollary 1 and Eq. 10 in Freund (2020). This means that we take a Beta(2-α,α) coalescent, but change the time scale with the function t →(gα)^-1^(exp(gαt)-1), where g is the exponential growth rate; i.e. the time changed coalescent at time t corresponds to the original coalscent at time (gα)^-1^(exp(gαt)-1). For Dirac coalescents, we consider the approach as described in Matuszewski et al. (2018), which adds exponential growth to the modified Moran models with skewed offspring distributions from Eldon and Wakeley (2006). This results in a Dirac coalescent whose time scale is changed by the function t → (1.5g)^-1^(exp(1.5gt)-1), regardless of the Dirac coalescent parameter (we choose γ=1.5 in the pre-limit modified Moran models).

We again approximate the Watterson estimator by approximating the total expected length of the coalescent tree with the mean over 1,000 simulations of the chosen Dirac coalescent with exponential growth. Due to the computational load, this led us to use a discrete prior, a uniform distribution on both the coalescent parameter and the growth parameter.

For both models, the growth rates were uniformly chosen as exp(g) from 10 equidistant steps g between 0 and log(5000) (including both values), in other words we used a discretized log uniform prior on growth rates. The coalescent parameter was similarly chosen from 10 equidistant steps between 1 and 1.9 for BETA (so 1,1.1,…,1.9) and between 0.05 and 0.95 for Dirac.

We performed a model section between these two models and the best fitting model class resulted from the main analysis (analysis 1 in Sup. Table 2). We performed 125,000 simulations (reduced to 80,000 for Stucki 2016 and Folkvardsen 2017 due to large computation times). As for all other scenarios, we drew the scaled population size from a binomial distribution on log-equally spaced discrete θ spanning one order of magnitude around the Watterson estimator (θobs), and then performed the ABC analysis as described above.

### Bayesian Skyline plots

We simulated data under the BETA coalescent with different values of alpha spanning the range of estimates obtained from the observed data (α=0.5, α=0.75, α=1, α=1.25, α=1.5; 10 simulations for each value of α). For this analysis we used the settings corresponding to a medium sized data set such as Eldholm 2015 (n = 250, s = 500, θ set to the generalized Watterson’s estimate 2s/E(L)). The simulated data sets are lists of mutations and their states (derived or ancestral) for all individuals. We transformed the simulated data in sequence data by randomly assigning nucleotides (A, T, C or G) to ancestral and derived mutations, drawing them from the empirical nucleotide frequency distribution of the data set Eldholm 2015. For all 50 simulated data sets we ran an extended Bayesian skyline analysis (Heled and Drummond 2008) with BEAST 2.5.0 (Bouckaert et al. 2019).

We assumed a strict clock, with clock rate equal to 5×10^-8^ nucleotide changes per site per year, a value that falls in the range of possible evolutionary rates for MTB (Menardo et al. 2019). Importantly assuming a different evolutionary rate would result in a different time scale, but it would not affect the demographic reconstruction. We assumed the GTR substitution model, and a 1/X [0-100,000] prior on the mean of the distribution of population sizes. For each data set we ran two runs of 400 million generations, discarded the first 40 million generation as burn-in and combined the two runs. For all data sets the effective sample sizes of the posterior distribution and of the number of changes in population size (sum(indicators.alltrees)) were larger than 180 (and in most cases larger than 200). The plots were produced with the plotEBSP R script available here (https://www.beast2.org/tutorials/). The simulated data and an example of the xml file are available at https://github.com/fabianHOH/mmc_R_gendiv/tree/master/MTB_MMC_repo.

## Acknowledgments

FM and SG were supported by the Swiss National Science Foundation (grants 310030_188888, IZRJZ3_164171, IZLSZ3_170834 and CRSII5_177163), the European Research Council (grant 309540-EVODRTB), and SystemsX.ch. FF was funded by the DFG grant FR 3633/2-1 through Priority Program 1590: Probabilistic Structures in Evolution. The authors acknowledge the support by the state of Baden-Württemberg through bwHPC. Part of the calculations were performed at sciCORE (http://scicore.unibas.ch/) scientific computing core facility at the University of Basel. We thank two anonymous reviewers for their comments and suggestions.

## Notes

https://github.com/fabianHOH/mmc_R_gendiv/tree/master/MTB_MMC_repo

### Competing Interest Statement

The authors have declared no competing interest.

## References

Ámason, E., & Halldórsdóttir, K. (2015). Nucleotide variation and balancing selection at the Ckma gene in Atlantic cod: analysis with multiple merger coalescent models. PeerJ, 3, e786.

Bainomugisa, A., Lavu, E., Hiashiri, S., Majumdar, S., Honjepari, A., Moke, R., … & Coulter, C. (2018). Multi-clonal evolution of multi-drug-resistant/extensively drug-resistant Mycobacterium tuberculosis in a high-prevalence setting of Papua New Guinea for over three decades. Microbial genomics, 4(2).

Behr, M. A., Edelstein, P. H., & Ramakrishnan, L. (2018). Revisiting the timetable of tuberculosis. Bmj, 362, k2738.

Berestycki, J., Berestycki, N., & Schweinsberg, J. (2013). The genealogy of branching Brownian motion with absorption. The Annals of Probability, 41(2), 527–618.

Birkner, M., Blath, J., Möhle, M., Steinrücken, M., & Tams, J. (2009). A modified lookdown construction for the Xi-Fleming-Viot process with mutation and populations with recurrent bottlenecks. Alea, 6, 25–61.

Biswas, S., & Akey, J. M. (2006). Genomic insights into positive selection. TRENDS in Genetics, 22(8), 437–446.

Bjorn-Mortensen, K., Soborg, B., Koch, A., Ladefoged, K., Merker, M., Lillebaek, T., … & Kohl, T. A. (2016). Tracing Mycobacterium tuberculosis transmission by whole genome sequencing in a high incidence setting: a retrospective population-based study in East Greenland. Scientific reports, 6, 33180.

Bolger, A. M., Lohse, M., & Usadel, B. (2014). Trimmomatic: A flexible trimmer for Illumina Sequence Data. Bioinformatics, btu170.

Bolthausen E, Sznitman AS (1998) On Ruelle’s probability cascades and an abstract cavity method. Communications in Mathematical Physics, 197, 247–276.

Bos, K. I., Harkins, K. M., Herbig, A., Coscolla, M., Weber, N., Comas, I., … & Campbell, T. J. (2014). PreColumbian mycobacterial genomes reveal seals as a source of New World human tuberculosis. Nature, 514(7523), 494.

Bouckaert, R., Vaughan, T. G., Barido-Sottani, J., Duchêne, S., Fourment, M., Gavryushkina, A., … & Matschiner, M. (2019). BEAST 2.5: An advanced software platform for Bayesian evolutionary analysis. PLoS computational biology, 15(4), e1006650.

Brown, T. S., Challagundla, L., Baugh, E. H., Omar, S. V., Mustaev, A., Auld, S. C., … & Narechania, A. (2019). Pre-detection history of extensively drug-resistant tuberculosis in KwaZulu-Natal, South Africa. Proceedings of the National Academy of Sciences, 116(46), 23284–23291.

Brunet, É., & Derrida, B. (2013). Genealogies in simple models of evolution. Journal of Statistical Mechanics: Theory and Experiment, 2013(01), P01006.

Casanova, A. G., Pina, V. M. and Siri-Jégousse, A. (2019). The symmetric coalescent and Wright-Fisher models with bottlenecks.” arXiv preprint arXiv:1903.05642.

Chiner-Oms, Á., Sánchez-Busó, L., Corander, J., Gagneux, S., Harris, S. R., Young, D., … & Comas, I. (2019). Genomic determinants of speciation and spread of the Mycobacterium tuberculosis complex. Science Advances, 5(6), eaaw3307.

Comas, I., Coscolla, M., Luo, T., Borrell, S., Holt, K. E., Kato-Maeda, M., … & Yeboah-Manu, D. (2013). Out-of-Africa migration and Neolithic coexpansion of Mycobacterium tuberculosis with modern humans. Nature genetics, 45(10), 1176.

Comas, I., Hailu, E., Kiros, T., Bekele, S., Mekonnen, W., Gumi, B., … & Goig, G. A. (2015). Population genomics of Mycobacterium tuberculosis in Ethiopia contradicts the virgin soil hypothesis for human tuberculosis in Sub-Saharan Africa. Current Biology, 25(24), 3260–3266.

Danecek, P., Auton, A., Abecasis, G., Albers, C. A., Banks, E., DePristo, M. A., … & McVean, G. (2011). The variant call format and VCFtools. Bioinformatics, 27(15), 2156–2158.

Desai, M. M., Walczak, A. M., & Fisher, D. S. (2013). Genetic diversity and the structure of genealogies in rapidly adapting populations. Genetics, 193(2), 565–585.

Drummond, A., & Rodrigo, A. G. (2000). Reconstructing genealogies of serial samples under the assumption of a molecular clock using serial-sample UPGMA. Molecular Biology and Evolution, 17(12), 1807–1815.

Drummond, A. J., Rambaut, A., Shapiro, B., & Pybus, O. G. (2005). Bayesian coalescent inference of past population dynamics from molecular sequences. Molecular biology and evolution, 22(5), 1185–1192.

Durrett, R., & Schweinsberg, J. (2005). A coalescent model for the effect of advantageous mutations on the genealogy of a population. Stochastic processes and their applications, 115(10), 1628–1657.

Eldholm, V., Monteserin, J., Rieux, A., Lopez, B., Sobkowiak, B., Ritacco, V., & Balloux, F. (2015). Four decades of transmission of a multidrug-resistant Mycobacterium tuberculosis outbreak strain. Nature communications, 6, 7119.

Eldholm, V., Pettersson, J. H. O., Brynildsrud, O. B., Kitchen, A., Rasmussen, E. M., Lillebaek, T., … & Balloux, F.. (2016). Armed conflict and population displacement as drivers of the evolution and dispersal of Mycobacterium tuberculosis. Proceedings of the National Academy of Sciences, 113(48), 13881–13886.

Eldon, B., Birkner, M., Blath, J., & Freund, F. (2015). Can the site-frequency spectrum distinguish exponential population growth from multiple-merger coalescents?. Genetics, 199(3), 841–856.

Eldon, B., & Wakeley, J. (2006). Coalescent processes when the distribution of offspring number among individuals is highly skewed. Genetics, 172(4), 2621–2633.

Excoffier, L., Dupanloup, I., Huerta-Sánchez, E., Sousa, V. C., & Foll, M. (2013). Robust demographic inference from genomic and SNP data. PLoS genetics, 9(10), e1003905.

Folkvardsen, D. B., Norman, A., Andersen, Å. B., Michael Rasmussen, E., Jelsbak, L., & Lillebaek, T. (2017). Genomic epidemiology of a major mycobacterium tuberculosis outbreak: retrospective cohort study in a low-incidence setting using sparse time-series sampling. The Journal of infectious diseases, 216(3), 366–374.

Freund, F., & Siri-Jégousse, A. (2019). Distinguishing coalescent models-which statistics matter most? BioRxiv, 679498.

Freund, F. (2020). Cannings models, population size changes and multiple-merger coalescents. Journal of Mathematical Biology, 80(5), 1497–1521.

Gagneux, S. (2018). Ecology and evolution of Mycobacterium tuberculosis. Nature Reviews Microbiology, 16(4), 202.

Gardy, J. L., Johnston, J. C., Sui, S. J. H., Cook, V. J., Shah, L., Brodkin, E., … & Varhol, R. (2011). Whole-genome sequencing and social-network analysis of a tuberculosis outbreak. New England Journal of Medicine, 364(8), 730–739.

Gelman, A., Carlin, J. B., Stern, H. S., Dunson, D. B., Vehtari, A., & Rubin, D. B. (2013). Model checking, in Bayesian data analysis. CRC press, 141–163.

Gillespie, J. H. (2000). Genetic drift in an infinite population: the pseudohitchhiking model. Genetics, 155(2), 909–919.

Griffiths, R. C., & Tavare, S. (1994). Sampling theory for neutral alleles in a varying environment. Phil. Trans. R. Soc. Lond. B, 344(1310), 403–410.

Heled, J., & Drummond, A. J. (2008). Bayesian inference of population size history from multiple loci. BMC Evolutionary Biology, 5(1), 289.

Heller, R., Chikhi, L., & Siegismund, H. R. (2013). The confounding effect of population structure on Bayesian skyline plot inferences of demographic history. PLoS One, 8(5), e62992.

Hernandez, R. D., Kelley, J. L., Elyashiv, E., Melton, S. C., Auton, A., McVean, G., … & Przeworski, M. (2011). Classic selective sweeps were rare in recent human evolution. Science, 331(6019), 920–924.

Hershberg, R., Lipatov, M., Small, P. M., Sheffer, H., Niemann, S., Homolka, S., … & Gagneux, S. (2008). High functional diversity in Mycobacterium tuberculosis driven by genetic drift and human demography. PLoS biology, 6(12), e311.

Horns, F., Vollmers, C., Dekker, C. L., & Quake, S. R. (2019). Signatures of selection in the human antibody repertoire: Selective sweeps, competing subclones, and neutral drift. Proceedings of the National Academy of Sciences, 116(4), 1261–1266.

Hoscheit, P., & Pybus, O. G. (2019). The multifurcating skyline plot. Virus Evolution, 5(2), vez031.

Huang, H., Ding, N., Yang, T., Li, C., Jia, X., Wang, G., … & Zong, Z. (2019). Cross-sectional whole-genome sequencing and epidemiological study of multidrug-resistant Mycobacterium tuberculosis in China. Clinical Infectious Diseases, 69(3), 405–413.

Hudson, R. R. (1983). Properties of a neutral allele model with intragenic recombination. Theoretical population biology, 23(2), 183–201.

Hudson, R. R. (2002). Generating samples under a Wright–Fisher neutral model of genetic variation. Bioinformatics, 18(2), 337–338.

Huillet, T. E. (2014). Pareto genealogies arising from a Poisson branching evolution model with selection. Journal of mathematical biology, 68(3), 727–761.

Irwin, K. K., Laurent, S., Matuszewski, S., Vuilleumier, S., Ormond, L., Shim, H., … & Jensen, J. D. (2016). On the importance of skewed offspring distributions and background selection in virus population genetics. Heredity, 117(6), 393.

Joy, D. A., Feng, X., Mu, J., Furuya, T., Chotivanich, K., Krettli, A. U., … & Beerli, P. (2003). Early origin and recent expansion of Plasmodium falciparum. Science, 300(5617), 318–321.

Kaplan, N. L., Darden, T., & Hudson, R. R. (1988). The coalescent process in models with selection. Genetics, 120(3), 819–829.

Kato, M., Vasco, D. A., Sugino, R., Narushima, D., & Krasnitz, A. (2017). Sweepstake evolution revealed by population-genetic analysis of copy-number alterations in single genomes of breast cancer. Royal Society open science, 4(9), 171060.

Kay, G. L., Sergeant, M. J., Zhou, Z., Chan, J. Z. M., Millard, A., Quick, J., … & Achtman, M. (2015). Eighteenth-century genomes show that mixed infections were common at time of peak tuberculosis in Europe. Nature communications, 6, 6717.

Kingman, J. F. C. (1982a). The coalescent. Stochastic processes and their applications, 13(3), 235–248.

Kingman, J. F. (1982b). On the genealogy of large populations. Journal of applied probability, 19(A), 27–43.

Koboldt, D., Zhang, Q., Larson, D., Shen, D., McLellan, M., Lin, L., Miller, C., Mardis, E., Ding, L., & Wilson, R. (2012). VarScan 2: Somatic mutation and copy number alteration discovery in cancer by exome sequencing Genome Research DOI: 10.1101/gr.129684.111

Kuhner, M. K. (2009). Coalescent genealogy samplers: windows into population history. Trends in Ecology & Evolution, 24(2), 86–93.

Lapierre, M., Blin, C., Lambert, A., Achaz, G., & Rocha, E. P. (2016). The impact of selection, gene conversion, and biased sampling on the assessment of microbial demography. Molecular biology and evolution, 33(7), 1711–1725.

Lee, R. S., Radomski, N., Proulx, J. F., Manry, J., McIntosh, F., Desjardins, F., … & Behr, M. A. (2015a). Reemergence and amplification of tuberculosis in the Canadian arctic. The Journal of infectious diseases, 211(12), 1905–1914.

Lee, R. S., Radomski, N., Proulx, J. F., Levade, I., Shapiro, B. J., McIntosh, F., … & Behr, M. A. (2015b). Population genomics of Mycobacterium tuberculosis in the Inuit. Proceedings of the National Academy of Sciences, 112(44), 13609–13614.

Lee, R. S., Proulx, J. F., McIntosh, F., Behr, M. A., & Hanage, W. P. (2020). Previously undetected superspreading of Mycobacterium tuberculosis revealed by deep sequencing. Elife, 9, e53245.

Li H. and Durbin R. (2009) Fast and accurate short read alignment with Burrows-Wheeler Transform. Bioinformatics, 25:1754–60.

Li H A statistical framework for SNP calling, mutation discovery, association mapping and population genetical parameter estimation from sequencing data. Bioinformatics. 2011 Nov 1;27(21):2987–93. Epub 2011 Sep 8. [PMID: 21903627]

Lin, P. L., Ford, C. B., Coleman, M. T., Myers, A. J., Gawande, R., Ioerger, T., … & Flynn, J. L. (2014). Sterilization of granulomas is common in active and latent tuberculosis despite within-host variability in bacterial killing. Nature medicine, 20(1), 75.

Liu, Q., Ma, A., Wei, L., Pang, Y., Wu, B., Luo, T., … & Zuo, T. (2018). China’s tuberculosis epidemic stems from historical expansion of four strains of Mycobacterium tuberculosis. Nature ecology & evolution, 2(12), 1982.

Luo, T., Comas, I., Luo, D., Lu, B., Wu, J., Wei, L., … & Shen, X. (2015). Southern East Asian origin and coexpansion of Mycobacterium tuberculosis Beijing family with Han Chinese. Proceedings of the National Academy of Sciences, 201424063.

Matuszewski, S., Hildebrandt, M. E., Achaz, G., & Jensen, J. D. (2018). Coalescent processes with skewed offspring distributions and nonequilibrium demography. Genetics, 208(1), 323–338.

McKenna, A., Hanna, M., Banks, E., Sivachenko, A., Cibulskis, K., Kernytsky, A., Garimella, K., Altshuler, D., Gabriel, S., Daly, M., DePristo, M. A., (2010). The Genome Analysis Toolkit: a MapReduce framework for analyzing next-generation DNA sequencing data. Genome Research 20:1297–303

Menardo, F., Duchêne, S., Brites, D. & Gagneux, S. (2019). The molecular clock of *Mycobacterium tuberculosis*. PLoS Pathogens 15(9): e1008067.

Merker, M., Barbier, M., Cox, H., Rasigade, J. P., Feuerriegel, S., Kohl, T. A., … & Andres, S. (2018). Compensatory evolution drives multidrug-resistant tuberculosis in Central Asia. eLife, 7, e38200.

Merker, M., Blin, C., Mona, S., Duforet-Frebourg, N., Lecher, S., Willery, E., … & Allix-Béguec, C. (2015). Evolutionary history and global spread of the Mycobacterium tuberculosis Beijing lineage. Nature genetics, 47(3), 242.

Möhle, Martin, and Serik Sagitov. A classification of coalescent processes for haploid exchangeable population models. The Annals of Probability 29.4 (2001): 1547–1562.

Morales-Arce, A. Y., Harris, R. B., Stone, A. C., & Jensen, J. D. (2020). Evaluating the contributions of purifying selection and progeny-skew in dictating within-host Mycobacterium tuberculosis evolution. Evolution.

Mulholland, C., Shockey, A., Aung, H., Cursons, R., O’Toole, R., Gautam, S., … & Cook, G. (2019). Dispersal of Mycobacterium tuberculosis driven by historical European trade in the South Pacific. Frontiers in microbiology, 10, 2778.

Neher, R. A., & Hallatschek, O. (2013). Genealogies of rapidly adapting populations. Proceedings of the National Academy of Sciences, 110(2), 437–442.

Neher, R. A., & Walczak, A. M. (2018). Progress and open problems in evolutionary dynamics. arXiv preprint arXiv:1804.07720.

Neuhauser, C., & Krone, S. M. (1997). The genealogy of samples in models with selection. Genetics, 145(2), 519–534.

Niwa, H. S., Nashida, K., Yanagimoto, T., & Handling editor: W. Stewart Grant. (2016). Reproductive skew in Japanese sardine inferred from DNA sequences. ICES Journal of Marine Science, 73(9), 2181–2189.

Nourmohammad, A., Otwinowski, J., Łuksza, M., Mora, T., & Walczak, A. M. (2019). Fierce selection and interference in B-cell repertoire response to chronic HIV-1. Molecular biology and evolution.

O’Neill, M. B., Shockey, A., Zarley, A., Aylward, W., Eldholm, V., Kitchen, A., & Pepperell, C. S. (2019). Lineage specific histories of Mycobacterium tuberculosis dispersal in Africa and Eurasia. Molecular ecology.

Pepperell, C. S., Casto, A. M., Kitchen, A., Granka, J. M., Cornejo, O. E., Holmes, E. C., … & Feldman, M. W. (2013). The role of selection in shaping diversity of natural M. tuberculosis populations. PLoS pathogens, 9(8), e1003543.

Pudlo, P., Marin, J. M., Estoup, A., Cornuet, J. M., Gautier, M., & Robert, C. P. (2015). Reliable ABC model choice via random forests. Bioinformatics, 32(6), 859–866.

Pybus, O. G., Charleston, M. A., Gupta, S., Rambaut, A., Holmes, E. C., & Harvey, P. H. (2001). The epidemic behavior of the hepatitis C virus. Science, 292(5525), 2323–2325.

Raynal, L., Marin, J. M., Pudlo, P., Ribatet, M., Robert, C. P., & Estoup, A. (2018). ABC random forests for Bayesian parameter inference. Bioinformatics, 35(10), 1720–1728.

Rocha, E. P. (2018). Neutral theory, microbial practice: challenges in bacterial population genetics. Molecular biology and evolution, 35(6), 1338–1347.

Roetzer, A., Diel, R., Kohl, T. A., Rückert, C., Nübel, U., Blom, J., … & Supply, P. (2013). Whole genome sequencing versus traditional genotyping for investigation of a Mycobacterium tuberculosis outbreak: a longitudinal molecular epidemiological study. PLoS medicine, 10(2), e1001387.

Rosenberg, N. A., & Nordborg, M. (2002). Genealogical trees, coalescent theory and the analysis of genetic polymorphisms. Nature Reviews Genetics, 3(5), 380.

Sargsyan, O., & Wakeley, J. (2008). A coalescent process with simultaneous multiple mergers for approximating the gene genealogies of many marine organisms. Theoretical population biology, 74(1), 104–114.

Schweinsberg, J. (2003). Coalescent processes obtained from supercritical Galton–Watson processes. Stochastic Processes and their Applications, 106(1), 107–139.

Schweinsberg, J. (2017). Rigorous results for a population model with selection II: genealogy of the population. Electronic Journal of Probability, 22.

Shitikov, E., Kolchenko, S., Mokrousov, I., Bespyatykh, J., Ischenko, D., Ilina, E., & Govorun, V. (2017). Evolutionary pathway analysis and unified classification of East Asian lineage of Mycobacterium tuberculosis. Scientific reports, 7(1), 9227.

Spence, J. P., Kamm, J. A., & Song, Y. S. (2016). The site frequency spectrum for general coalescents. Genetics, 202(4), 1549–1561.

Stamatakis A. RAxML version 8: a tool for phylogenetic analysis and post-analysis of large phylogenies. Bioinformatics. 2014;30(9): 1312–1313. doi: 10.1093/bioinformatics/btu033.

Stucki, D., Ballif, M., Bodmer, T., Coscolla, M., Maurer, A. M., Droz, S., … & Fenner, L. (2015). Tracking a tuberculosis outbreak over 21 years: strain-specific single-nucleotide polymorphism typing combined with targeted whole-genome sequencing. The Journal of infectious diseases, 211(8), 1306–1316.

Stucki, D., Brites, D., Jeljeli, L., Coscolla, M., Liu, Q., Trauner, A., … & Gagneux, S.. (2016). Mycobacterium tuberculosis lineage 4 comprises globally distributed and geographically restricted sublineages. Nature genetics, 48(12), 1535.

Tellier, A., & Lemaire, C. (2014). Coalescence 2.0: a multiple branching of recent theoretical developments and their applications. Molecular ecology, 23(11), 2637–2652.

Walker, T. M., Ip, C. L., Harrell, R. H., Evans, J. T., Kapatai, G., Dedicoat, M. J., … & Parkhill, J. (2013). Whole-genome sequencing to delineate Mycobacterium tuberculosis outbreaks: a retrospective observational study. The Lancet infectious diseases, 13(2), 137–146.

Wilkinson-Herbots, H. M. (1998). Genealogy and subpopulation differentiation under various models of population structure. Journal of Mathematical Biology, 37(6), 535–585.

World Health Organization. (2019) Global tuberculosis report 2019. (https://www.who.int/tb/publications/global_report/en/).

Wright, E. S., & Vetsigian, K. H. (2019). Stochastic exits from dormancy give rise to heavy-tailed distributions of descendants in bacterial populations. Molecular ecology, 28(17), 3915–3928.

Yang, Z. (2007). PAML 4: phylogenetic analysis by maximum likelihood. Molecular biology and evolution, 24(8), 1586–1591.

Ypma, R. J., Altes, H. K., van Soolingen, D., Wallinga, J., & van Ballegooijen, W. M. (2013). A sign of superspreading in tuberculosis: highly skewed distribution of genotypic cluster sizes. Epidemiology, 24(3), 395–400.

