## Supplementary Material for "Multiple merger genealogies in outbreaks of *Mycobacterium tuberculosis*"

#### Appendix 1. Sensitivity to the choice of prior distributions

An important aspect of Bayesian analyses is to test whether the results are robust to different priors and model assumptions. We performed a set of analyses investigating the sensitivity of the results of the ABC to changes of the prior of 1) the growth rate g for the KM+exp model, 2) the parameter α for the BETA model, and 3) the scaled population size θ for all models.

1) In our analysis the uniform prior on the growth rate in the KM+exp model placed a large weight on large growth rates (g), and almost none on small ones. Because of this, data simulated under a KM+exp model with low g was likely to be misclassified as MMC also in absence of serial sampling (Fig. 2, c' =0). When we replaced the uniform prior on g with a logarithmic one, the probability of misclassifying data generated under KM+exp as MMC was low for all values of g (Sup. Figs. 28-29, c' =0 ). We tested whether the prior choice could have biased the results of model selection, and repeated the ABC-RF analysis using the logarithmic prior. We found the same best fitting model for 21 of the 23 data sets analyzed (including subsets; Sup. Table 2: Analysis 2), two data sets switched from KM+exp to BETA (Stucki 2015 and Folkvardsen 2017 sampled in 2010).

2) We noticed that for some of the data sets that resulted in BETA as best fitting model, the weight of the posterior distribution of α was heavily shifted toward one (e.g. Bjorn-Mortensen 2016, Sup. Figs. 30-32). This could be both an indication that the data fitted well values of α close to one, but could also be due to prior misspecification. We therefore repeated the ABC analysis extending the prior of α to the range [0,2] (and with a log prior on g). With α < 1 the Beta coalescent does not correspond to an explicit population model, but describes multiple merger genealogies with larger multiple mergers compared to 1< α < 2. We found that for most data sets the posterior distribution of α was not strongly influenced by the different priors. However, for five data sets the median of the posterior estimate of α was below one, confirming that these were cases of prior misspecification (Sup. Table 2: Analysis 3; Sup. Figs. 33-35). The results of the model selection did not change significantly, one data set switched from BETA to KM (Lee 2015 Clade B), one from BETA to Dirac (Lee 2015 Clade C), and one from BETA to KM+exp (Comas 2015; Sup. Table 2). However the data set Comas 2015 fitted well to BETA with prior on α [1,2], therefore the better fit to the KM+exp is likely an artifact due to adding prior mass on values of α < 1 that do not fit the data.

Finally, the Bolthausen-Sznitman coalescent corresponds to the BETA coalescent with $\alpha = 1$. The value $\alpha=1$ is included in the prior of all analyses reported so far. However, because we used continuous priors, simulations were performed with values very close to one, but not exactly one. We repeated the analysis with the prior [0,2], but we additionally performed 1% of the simulations with $\alpha=1$ (Methods). The results of the model selection did not change further (Sup. Table 2: Analysis 4).

3) In our main analysis we drew the value of the scaled population size $\theta$ from a prior spanning one order of magnitude around the Watterson estimator ($\theta_{obs}$). We wanted to check the robustness of the results of the ABC to a less informative prior on $\theta$, and repeated the main analysis using a prior spanning two orders of magnitude around $\theta_{obs}$, i.e. 11 steps in [$\theta_{obs}$/10,10$\theta_{obs}$], see Methods. We found that all data sets resulted in the same best fitting model compared to main analysis (Sup. Table 2: Analysis 5), indicating that the analysis is robust to moderate variations to the prior on $\theta$.

#### Supplementary Tables

**Supplementary Table 1:**

List of accession numbers used in this study

File: Supplementary_table1.xslx

**Supplementary Table 2:**

Results of all ABC analyses

File: Supplementary_table2.xslx

**Supplementary Table 3.**

Sampling period and estimated age of the most recent common ancestor for the data sets that resulted in BETA or Dirac as best fitting model. For Comas 2015 this data was not available.

| **Data set** | **Sampling window in years** | **Age of the tree, in years before most recent sample**[1] |
|---|---|---|
| Eldholm 2015 | 14 | ~ 40 |
| Lee 2015 | 22 | ~ 100 |
| Roetzer 2013 | 14 | ~ 15 |
| Bainomugisa 2018 | 4 | ~ 50 |
| Bjorn-Mortensen 2016 | 21 | ~ 25 |
| Folkvardsen 2017 | 23 | ~ 55 |
| Stucki 2015 | 21 | NA |
| Eldholm 2016 | 6 | ~ 10 |

[1] The estimated age of the most recent common ancestors were obtained from the original publications

### Supplementary Figures

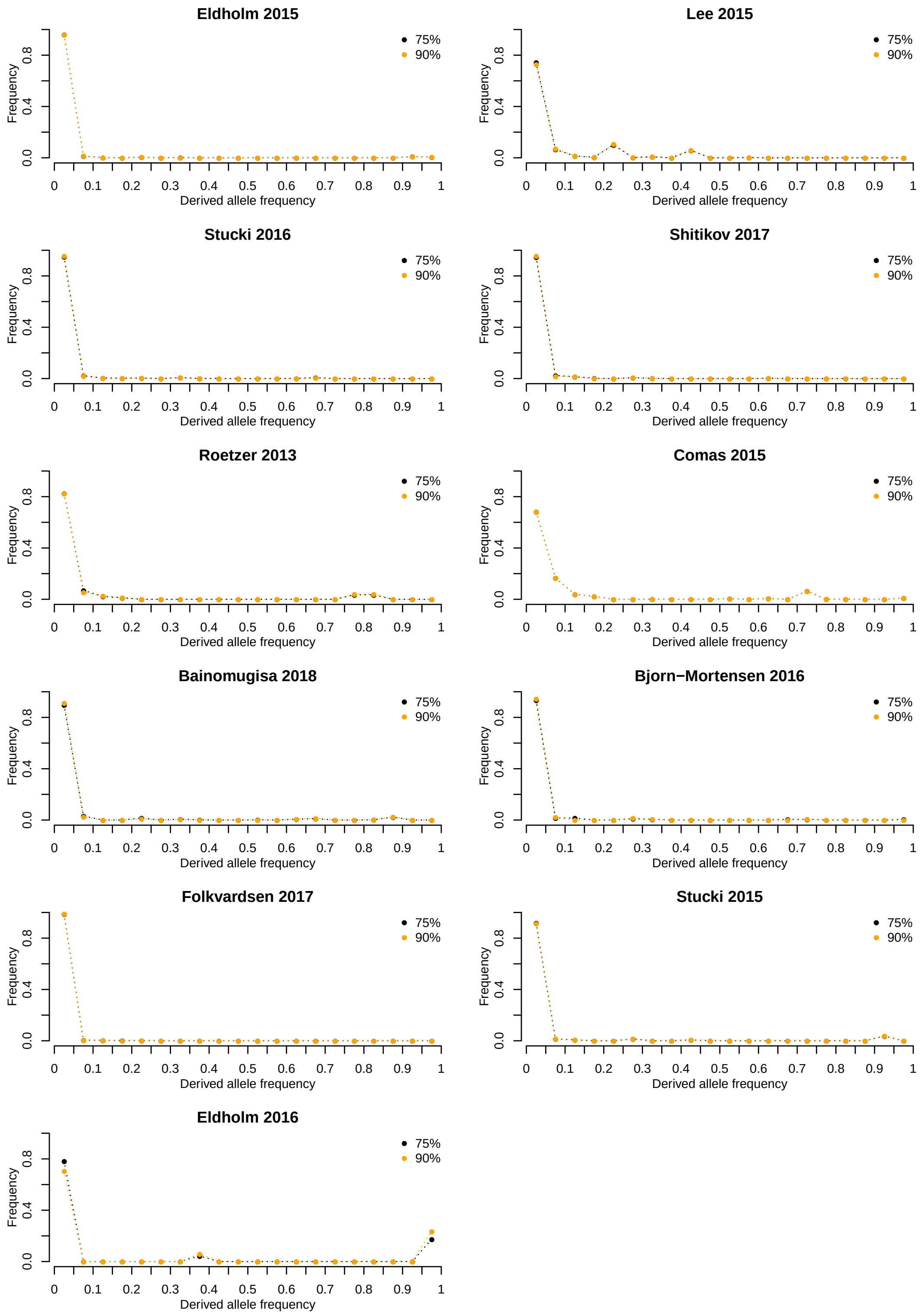


**Supplementary Figure 1.** Allele frequency spectrum of the full data sets with two different SNP calls. In black the data obtained with a minimum proportion of read supporting a genotype call of 75%, in orange 90% (See methods for details).

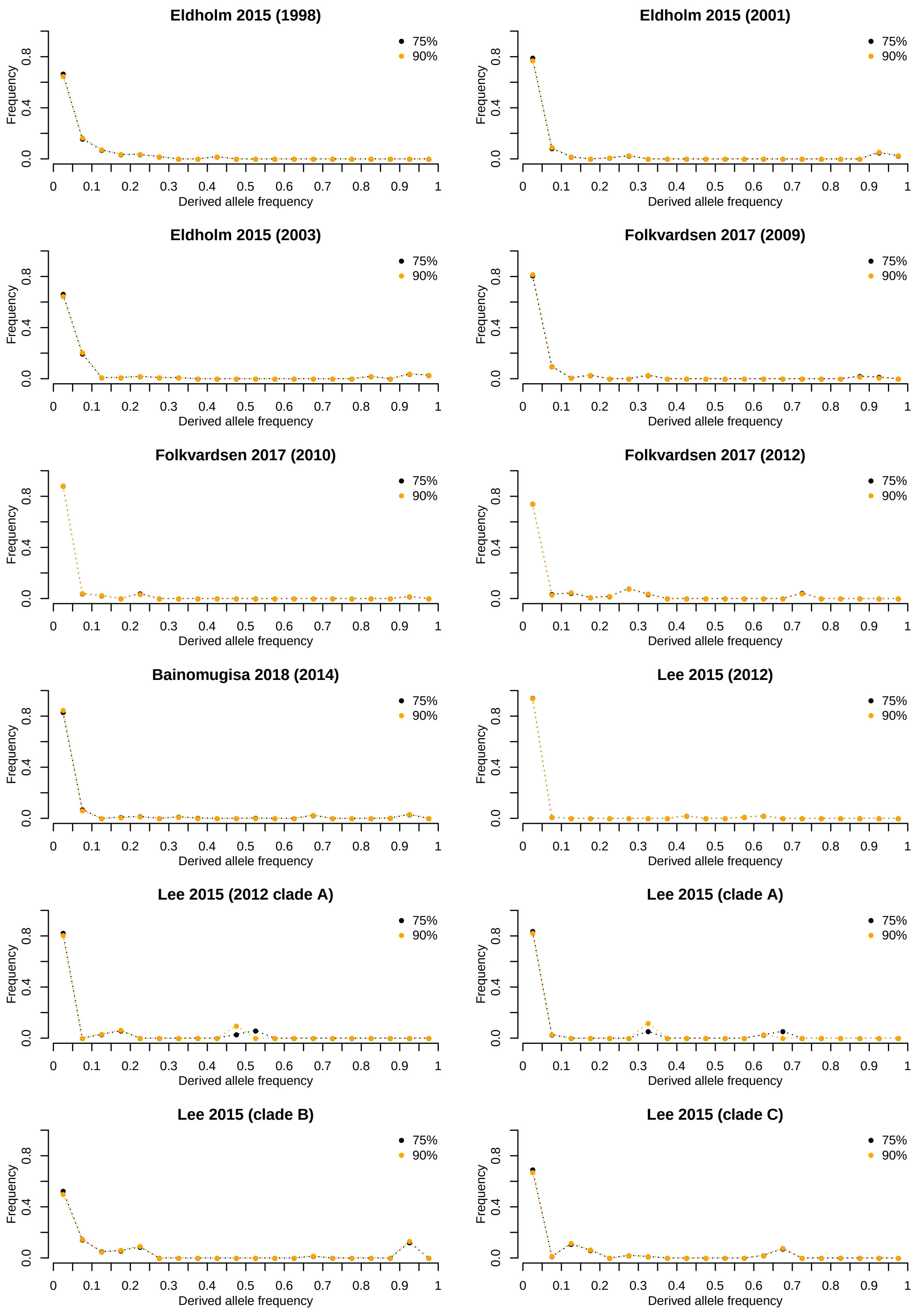


**Supplementary Figure 2.** Allele frequency spectrum of all subsets with two different SNP calls. In black the data obtained with a minimum proportion of read supporting a genotype call of 75%, in orange 90% (See methods for details).

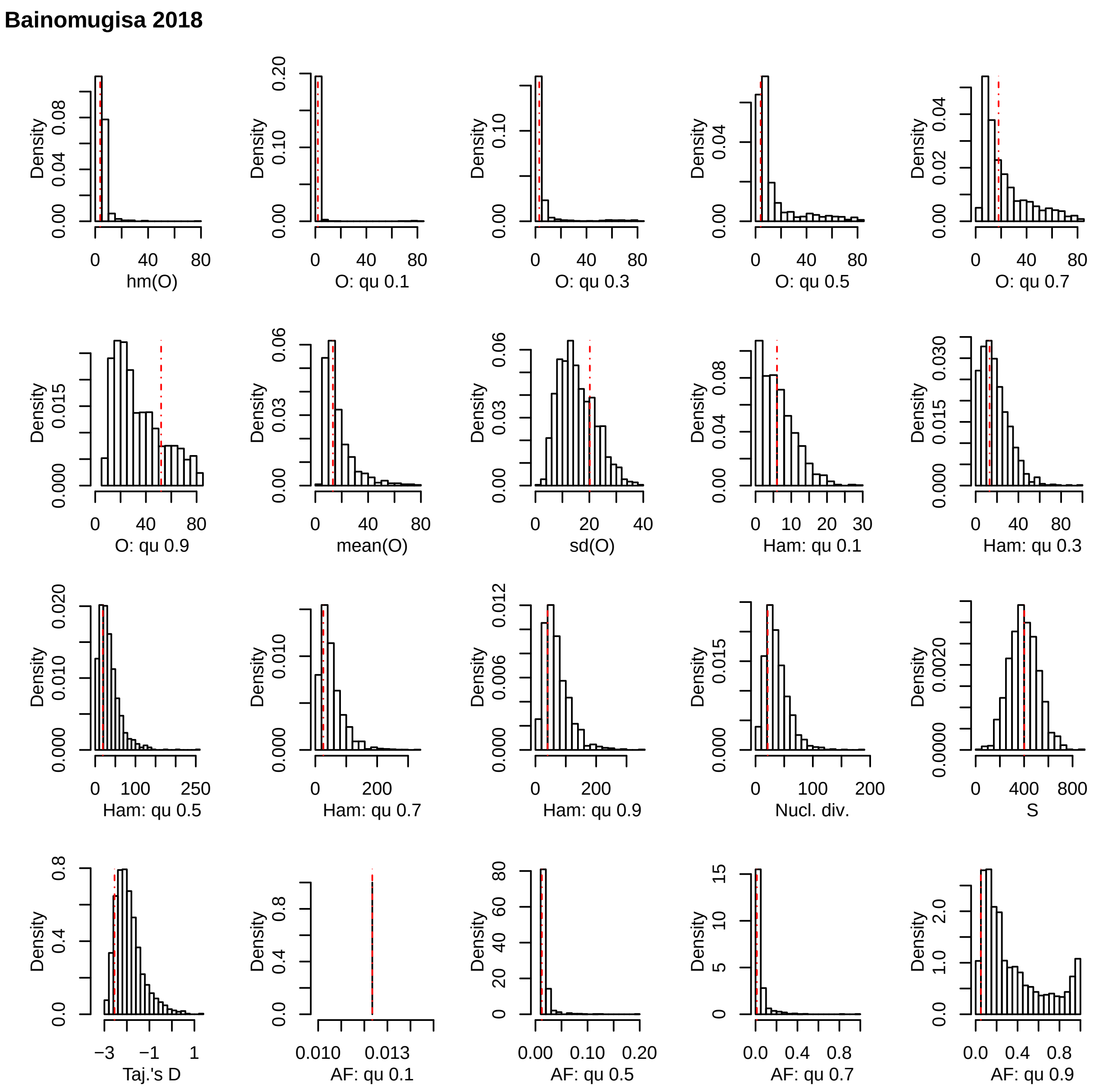


**Supplementary Figure 3.** Posterior predictive check for the data set Bainomugisa 2018. The red lines represent the values for the observed data, the histograms represent the results of 10,000 simulations under the best fitting model (BETA) using the median of the posterior distribution of the parameter α. hm: harmonic mean; qu: quantile; sd: standard deviation; O: minimal observable clade size; Ham: Hamming distance; Nuc. Div.: nucleotide diversity (π); S: number of polymorphic positions; Taj's D: Tajima's D; AF: mutant allele frequency.

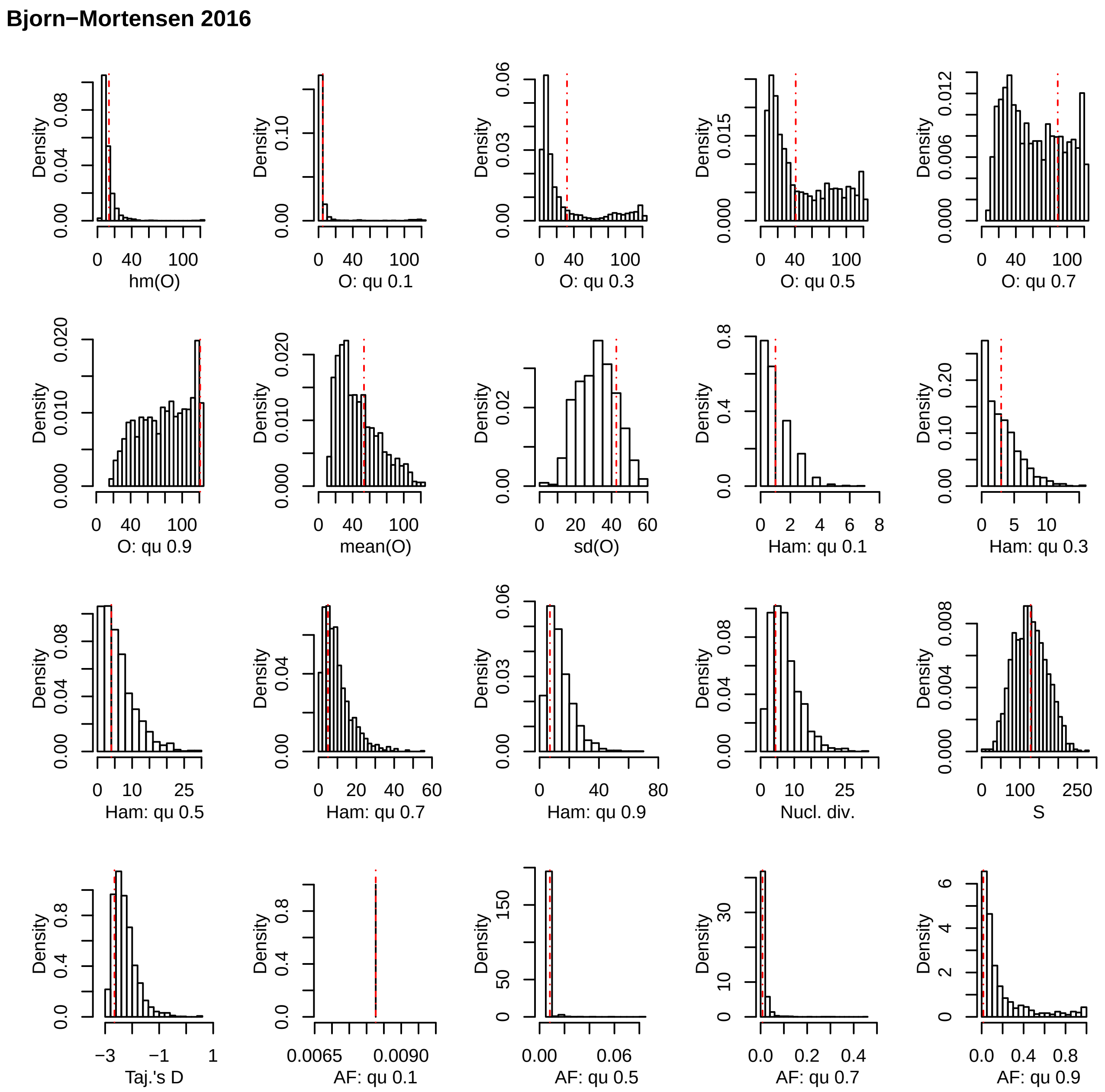


**Supplementary Figure 4.** Posterior predictive check for the data set Bjorn-Mortensen 2016. The red lines represent the values for the observed data, the histograms represent the results of 10,000 simulations under the best fitting model (BETA) using the median of the posterior distribution of the parameter α. hm: harmonic mean; qu: quantile; sd: standard deviation; O: minimal observable clade size; Ham: Hamming distance; Nuc. Div.: nucleotide diversity (π); S: number of polymorphic positions; Taj's D: Tajima's D; AF: mutant allele frequency.

**Comas 2015**

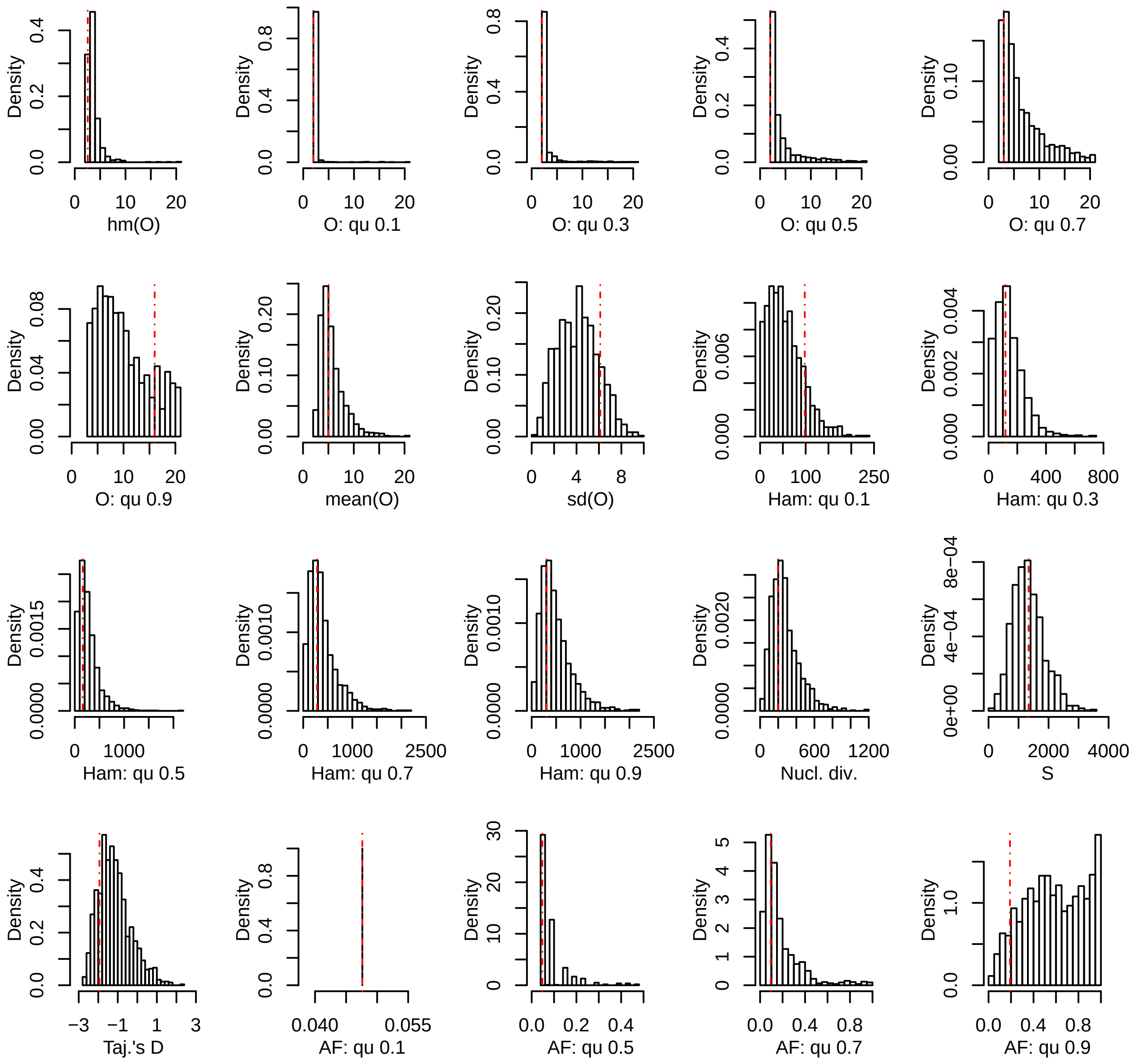


**Supplementary Figure 5.** Posterior predictive check for the data set Comas 2015. The red lines represent the values for the observed data, the histograms represent the results of 10,000 simulations under the best fitting model (BETA) using the median of the posterior distribution of the parameter α. hm: harmonic mean; qu: quantile; sd: standard deviation; O: minimal observable clade size; Ham: Hamming distance; Nuc. Div.: nucleotide diversity (π); S: number of polymorphic positions; Taj's D: Tajima's D; AF: mutant allele frequency.

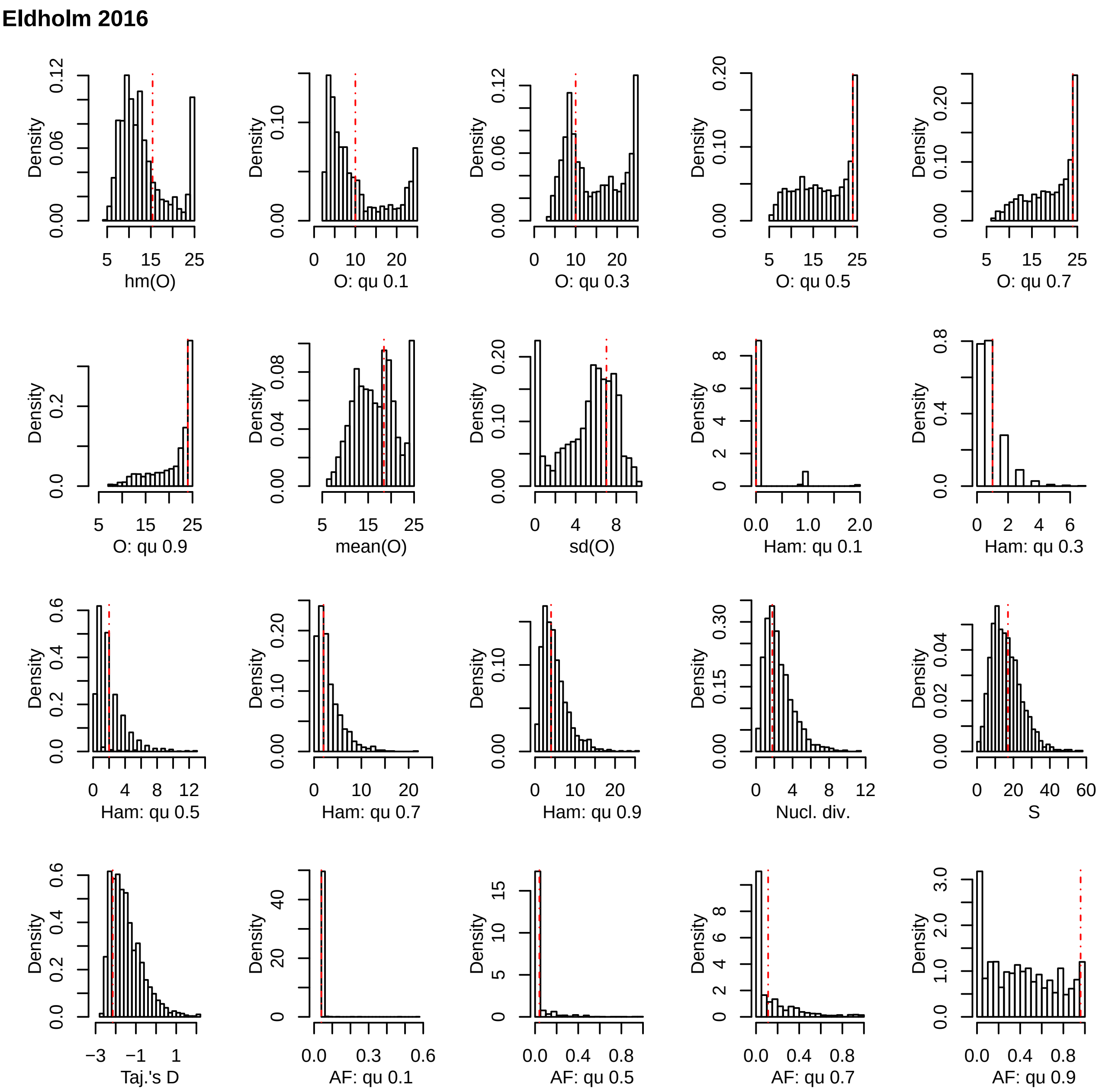


**Supplementary Figure 6**. Posterior predictive check for the data set Eldholm 2016. The red lines represent the values for the observed data, the histograms represent the results of 10,000 simulations under the best fitting model (Dirac) using the median of the posterior distribution of the parameter α. hm: harmonic mean; qu: quantile; sd: standard deviation; O: minimal observable clade size; Ham: Hamming distance; Nuc. Div.: nucleotide diversity (π); S: number of polymorphic positions; Taj's D: Tajima's D; AF: mutant allele frequency.

**Eldholm 2015**

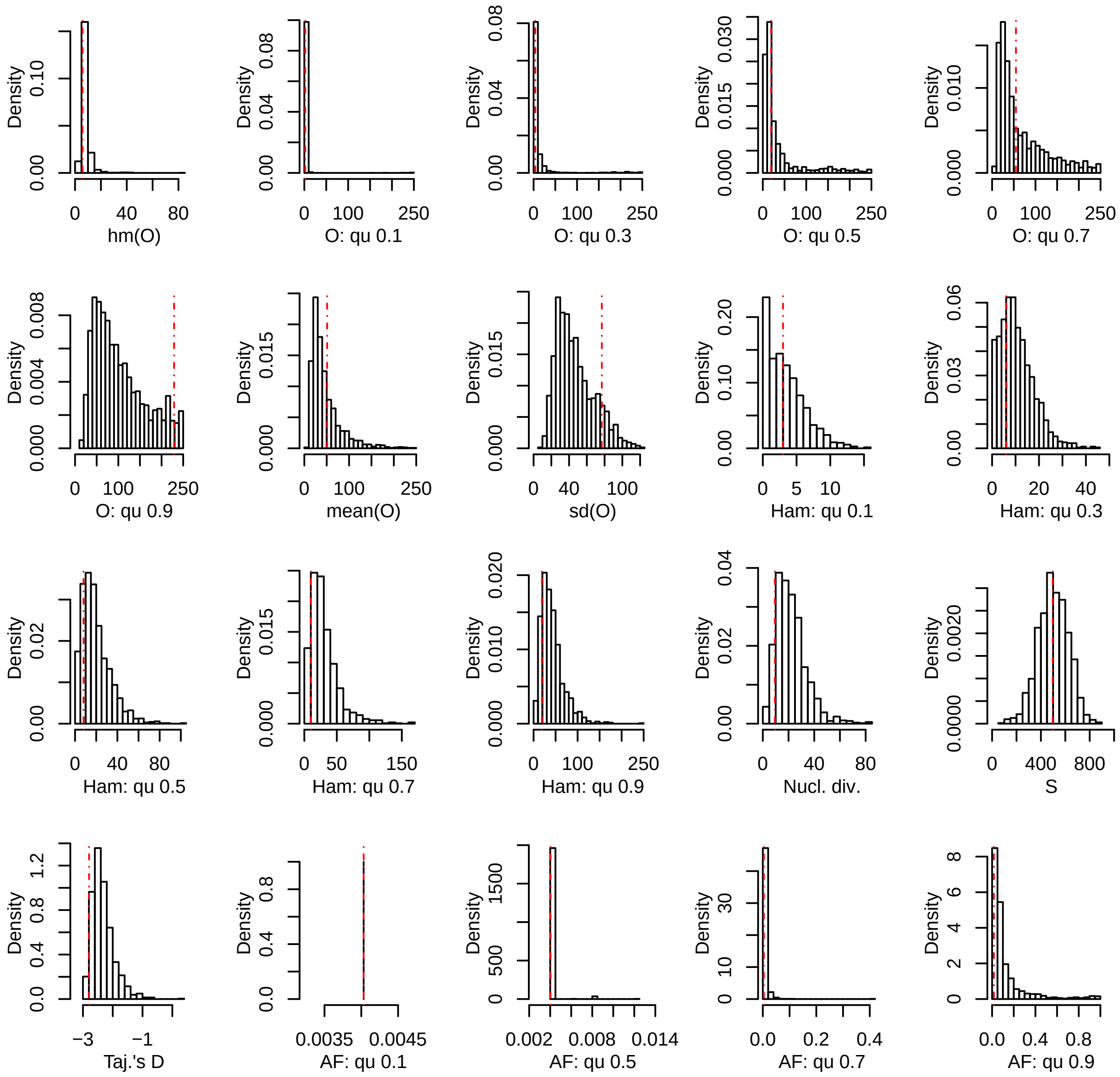


**Supplementary Figure 7**. Posterior predictive check for the data set Eldholm 2015. The red lines represent the values for the observed data, the histograms represent the results of 10,000 simulations under the best fitting model (BETA) using the median of the posterior distribution of the parameter α. hm: harmonic mean; qu: quantile; sd: standard deviation; O: minimal observable clade size; Ham: Hamming distance; Nuc. Div.: nucleotide diversity (π); S: number of polymorphic positions; Taj’s D: Tajima’s D; AF: mutant allele frequency.

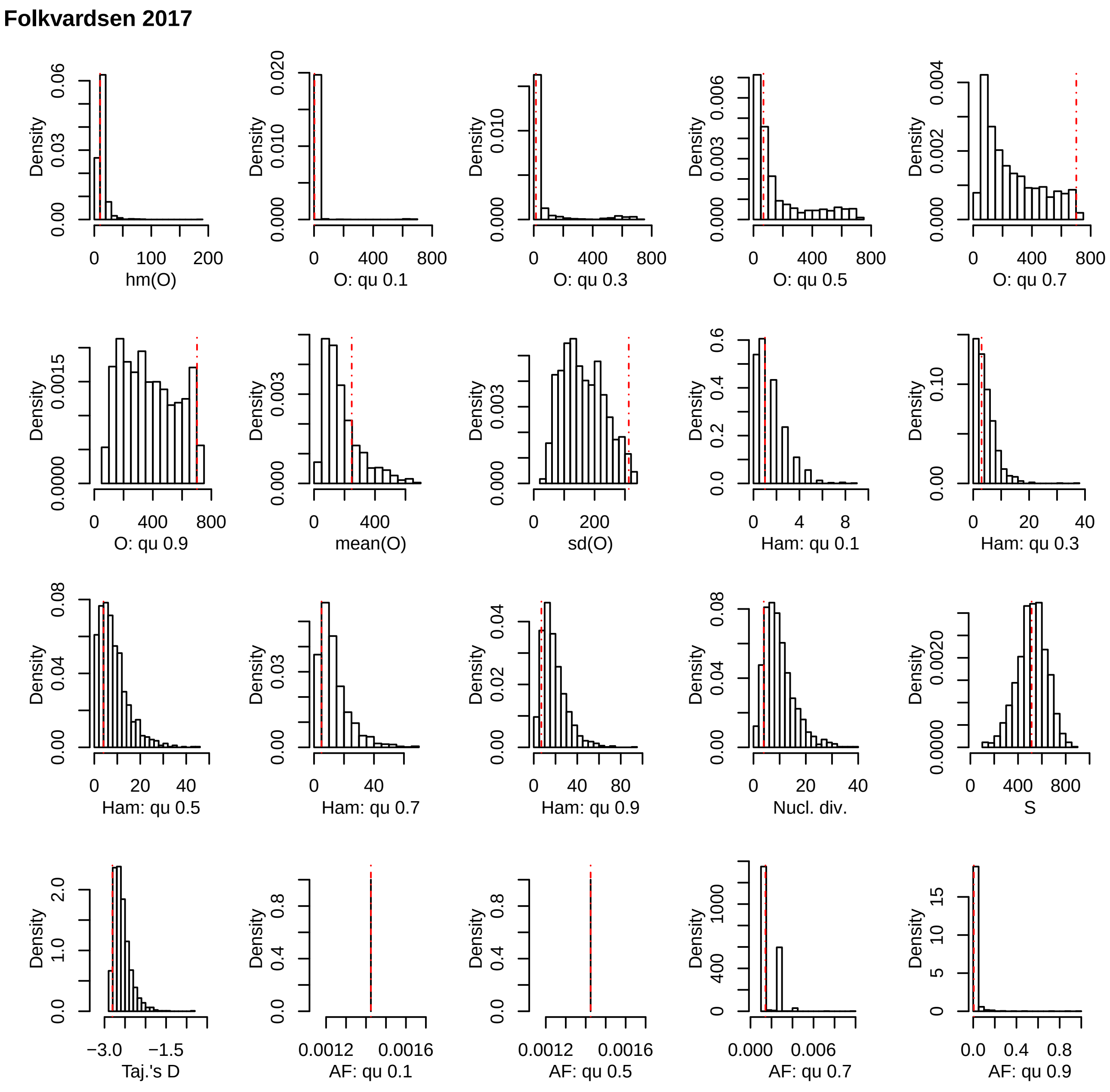


**Supplementary Figure 8**. Posterior predictive check for the data set Folkvardsen 2017. The red lines represent the values for the observed data, the histograms represent the results of 10,000 simulations under the best fitting model (BETA) using the median of the posterior distribution of the parameter α. hm: harmonic mean; qu: quantile; sd: standard deviation; O: minimal observable clade size; Ham: Hamming distance; Nuc. Div.: nucleotide diversity (π); S: number of polymorphic positions; Taj's D: Tajima's D; AF: mutant allele frequency.

**Lee 2015**

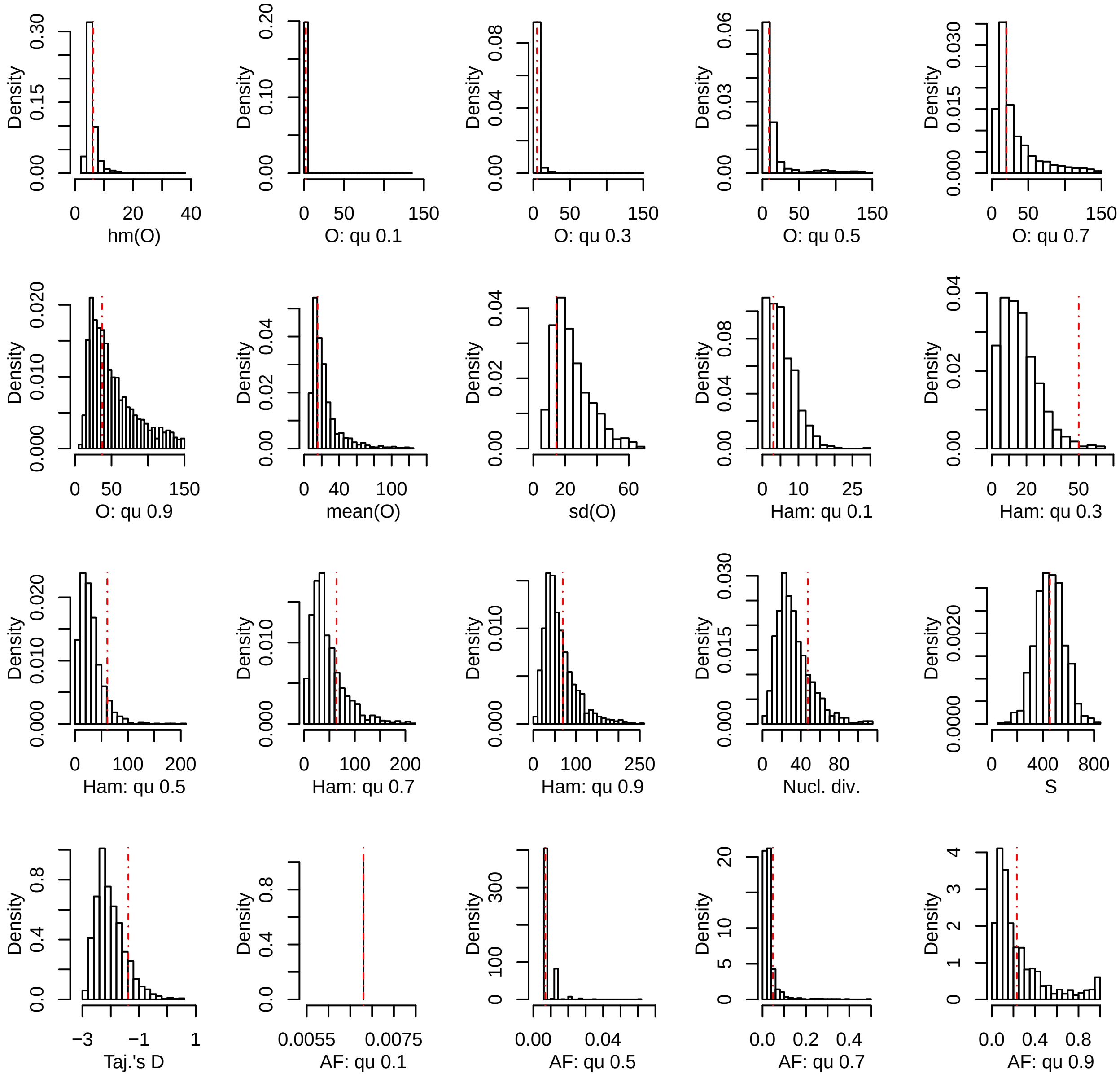


**Supplementary Figure 9**. Posterior predictive check for the data set Lee 2015. The red lines represent the values for the observed data, the histograms represent the results of 10,000 simulations under the best fitting model (BETA) using the median of the posterior distribution of the parameter α. hm: harmonic mean; qu: quantile; sd: standard deviation; O: minimal observable clade size; Ham: Hamming distance; Nuc. Div.: nucleotide diversity (π); S: number of polymorphic positions; Taj’s D: Tajima’s D; AF: mutant allele frequency.

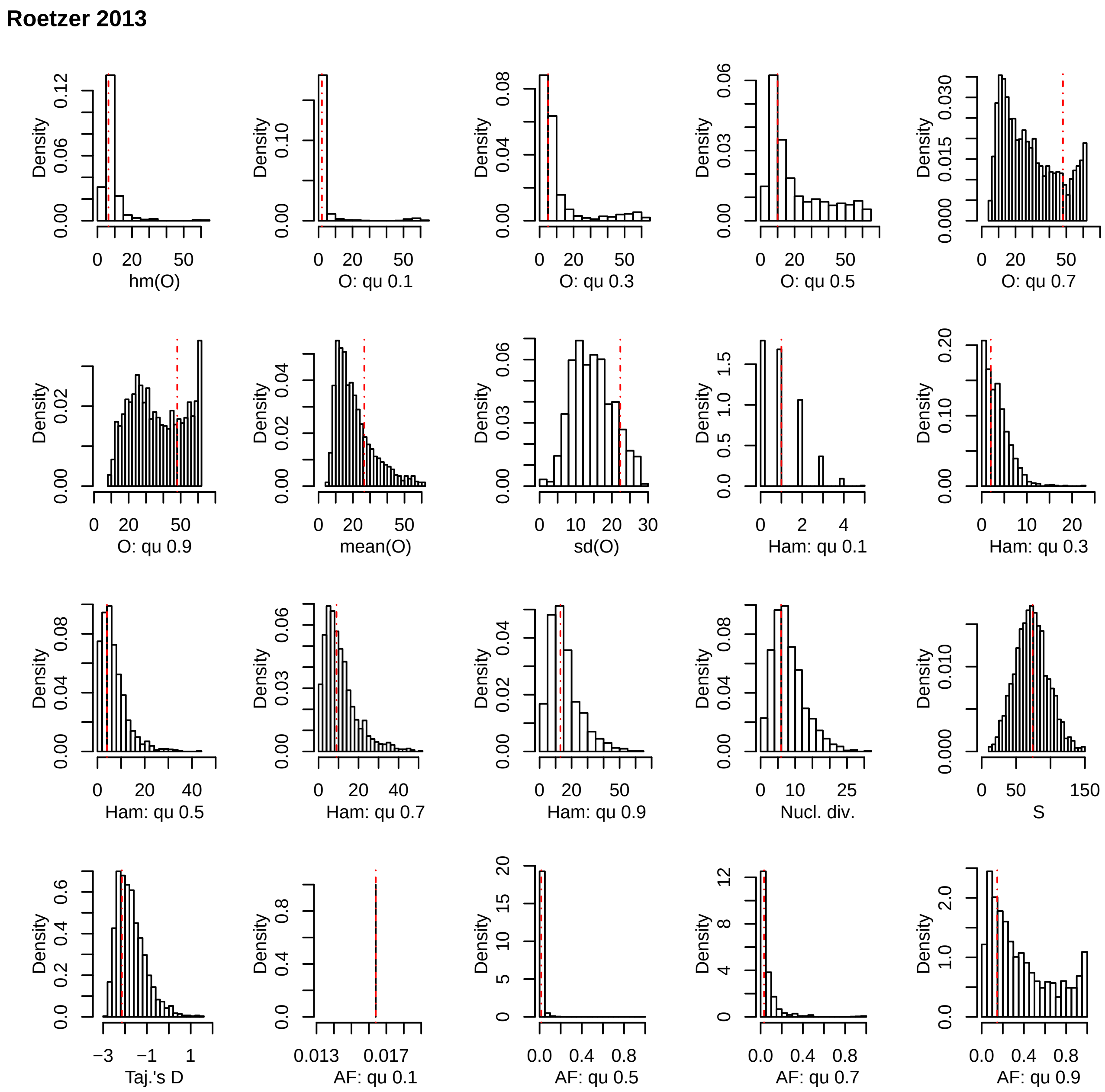


**Supplementary Figure 10.** Posterior predictive check for the data set Roetzer 2013. The red lines represent the values for the observed data, the histograms represent the results of 10,000 simulations under the best fitting model (BETA) using the median of the posterior distribution of the parameter α. hm: harmonic mean; qu: quantile; sd: standard deviation; O: minimal observable clade size; Ham: Hamming distance; Nuc. Div.: nucleotide diversity (π); S: number of polymorphic positions; Taj’s D: Tajima’s D; AF: mutant allele frequency.

**Stucki 2015**

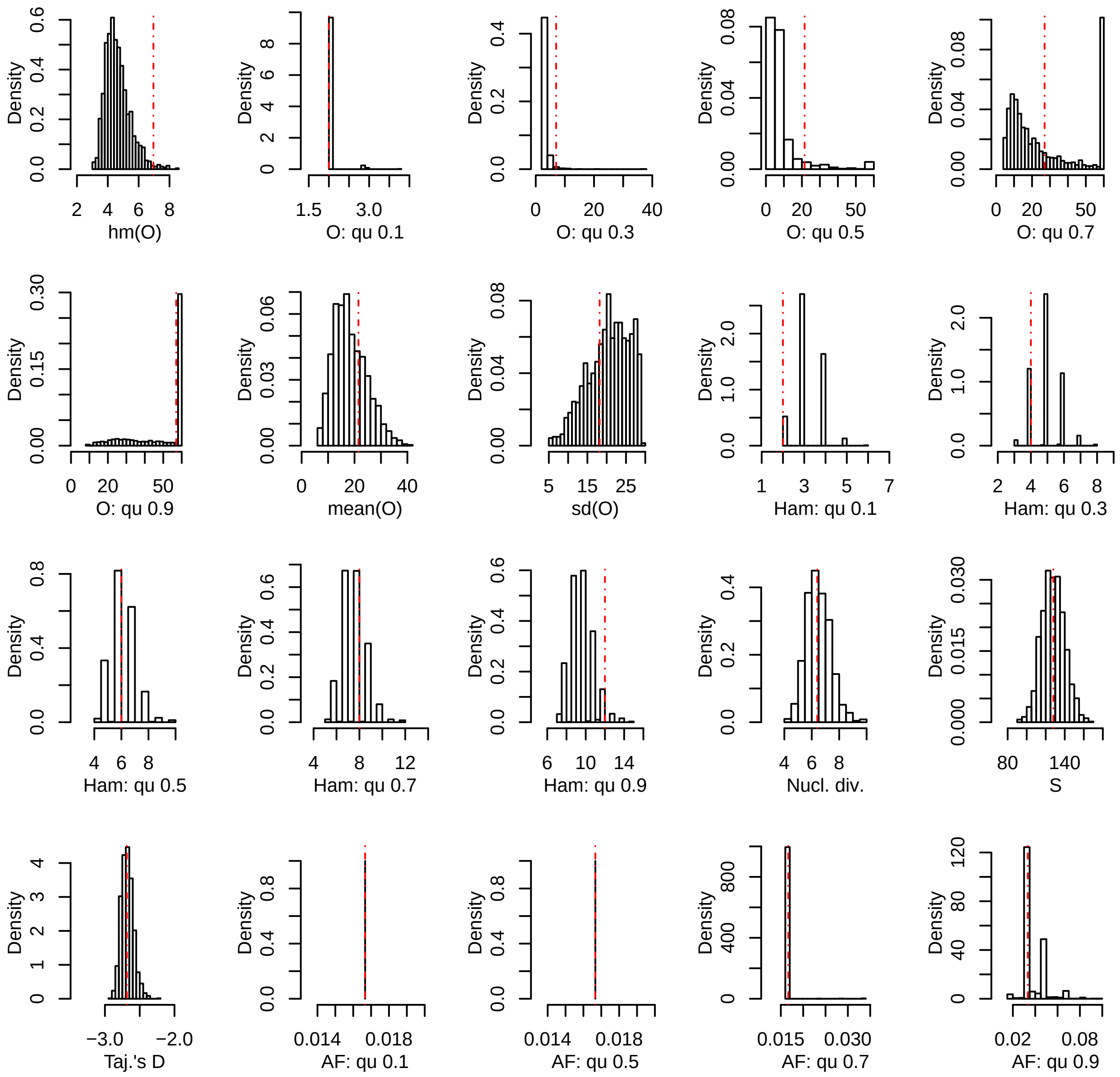


**Supplementary Figure 11.** Posterior predictive check for the data set Stucki 2015. The red lines represent the values for the observed data, the histograms represent the results of 10,000 simulations under the best fitting model (BETA) using the median of the posterior distribution of the parameter α. hm: harmonic mean; qu: quantile; sd: standard deviation; O: minimal observable clade size; Ham: Hamming distance; Nuc. Div.: nucleotide diversity (π); S: number of polymorphic positions; Taj's D: Tajima's D; AF: mutant allele frequency.

**Stucki 2016**

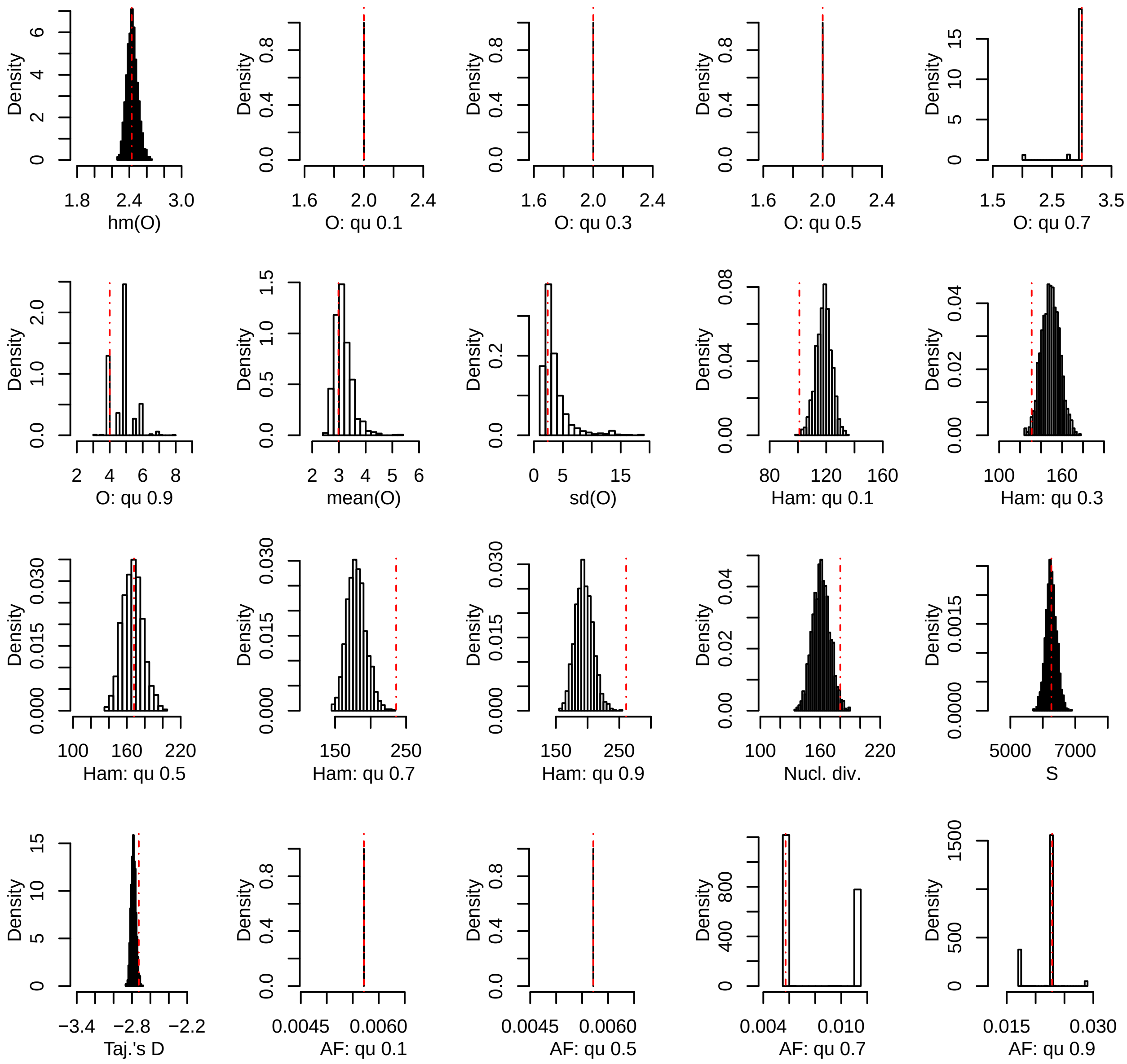


**Supplementary Figure 12.** Posterior predictive check for the data set Stucki 2016. The red lines represent the values for the observed data, the histograms represent the results of 10,000 simulations under the best fitting model (KM+exp) using the median of the posterior distribution of the parameter α. hm: harmonic mean; qu: quantile; sd: standard deviation; O: minimal observable clade size; Ham: Hamming distance; Nuc. Div.: nucleotide diversity (π); S: number of polymorphic positions; Taj’s D: Tajima’s D; AF: mutant allele frequency.

**Shitikov 2017**

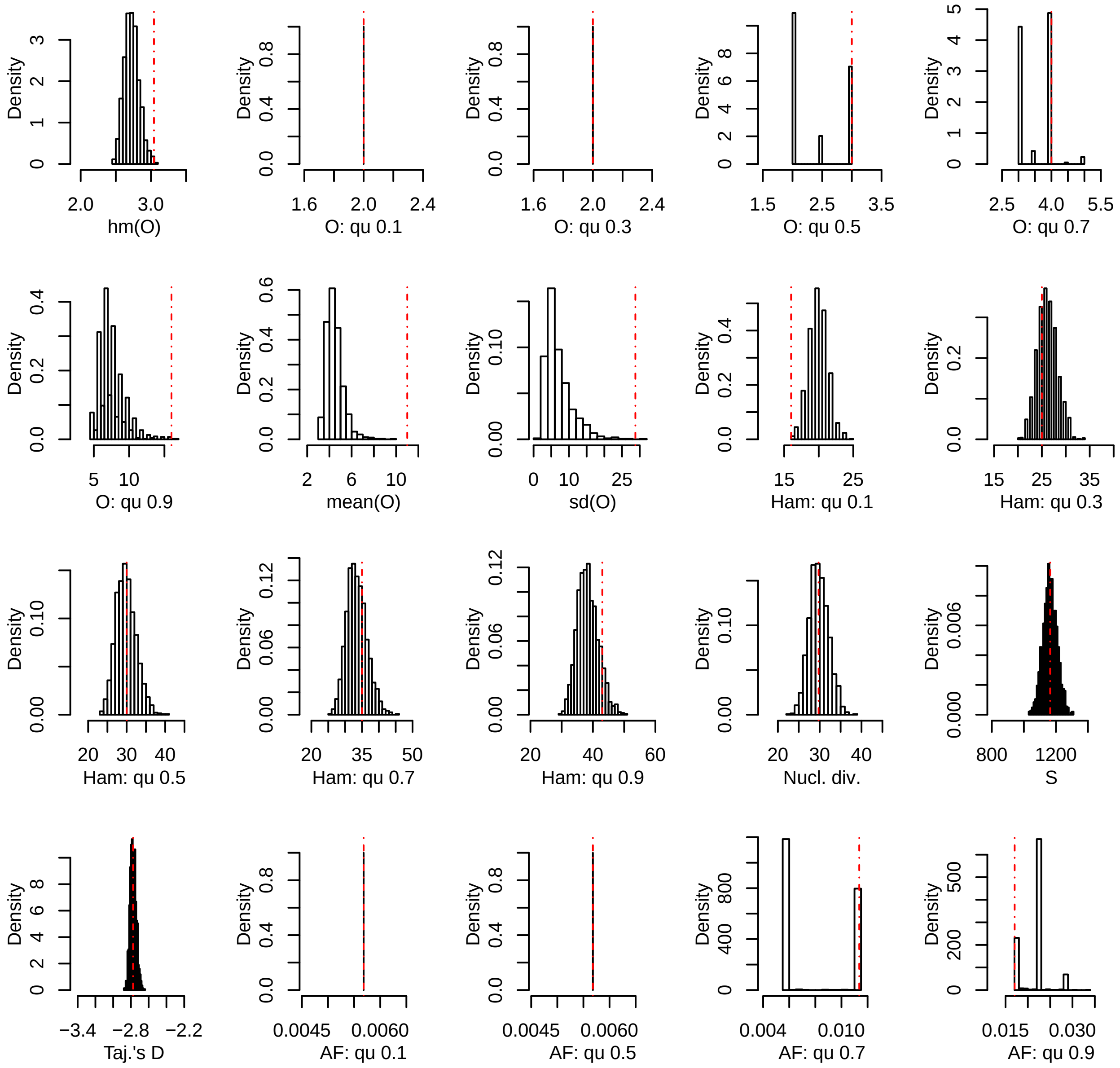


**Supplementary Figure 13.** Posterior predictive check for the data set Shitikov 2017. The red lines represent the values for the observed data, the histograms represent the results of 10,000 simulations under the best fitting model (KM+exp) using the median of the posterior distribution of the parameter α. hm: harmonic mean; qu: quantile; sd: standard deviation; O: minimal observable clade size; Ham: Hamming distance; Nuc. Div.: nucleotide diversity (π); S: number of polymorphic positions; Taj’s D: Tajima’s D; AF: mutant allele frequency.

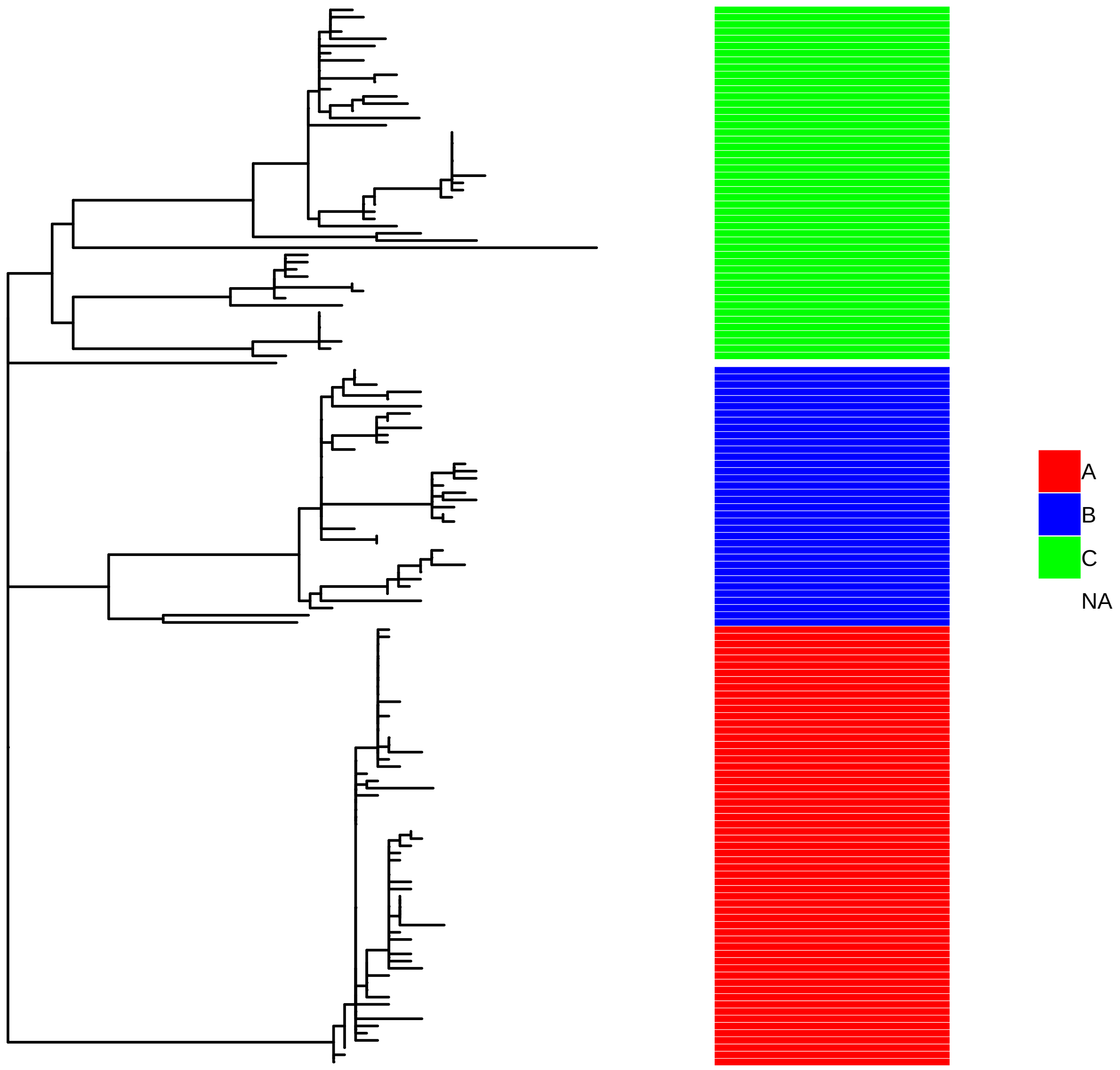


**Supplementary Figure 14.** Phylogenetic tree of the data set Lee 2015 with the three sub-clades highlighted.

**Lee 2015 Clade A**

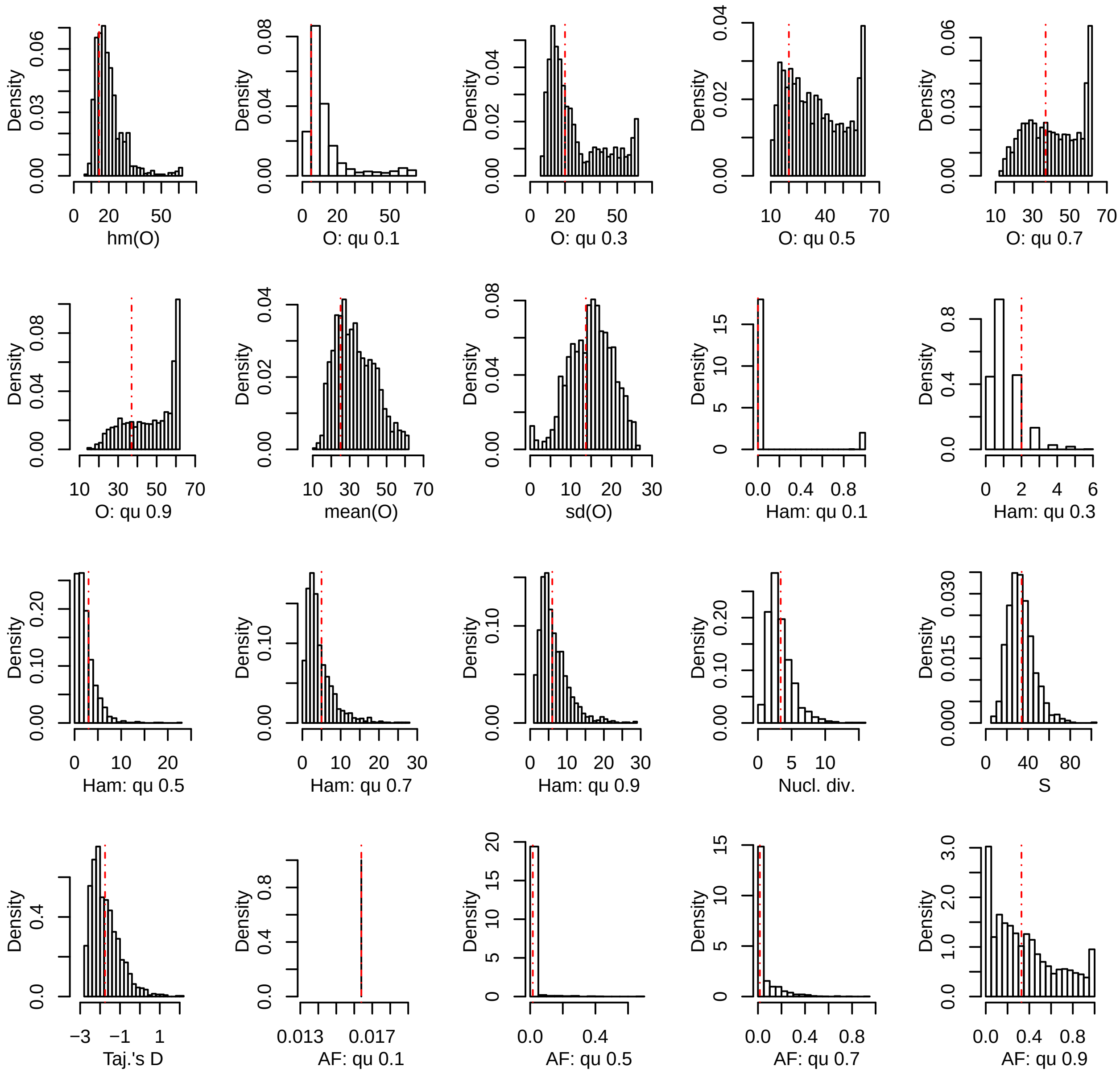


**Supplementary Figure 15.** Posterior predictive check for the data set Lee 2015 clade A. The red lines represent the values for the observed data, the histograms represent the results of 10,000 simulations under the best fitting model (Dirac) using the median of the posterior distribution of the parameter α. hm: harmonic mean; qu: quantile; sd: standard deviation; O: minimal observable clade size; Ham: Hamming distance; Nuc. Div.: nucleotide diversity (π); S: number of polymorphic positions; Taj’s D: Tajima’s D; AF: mutant allele frequency.

#### Lee 2015 Clade B

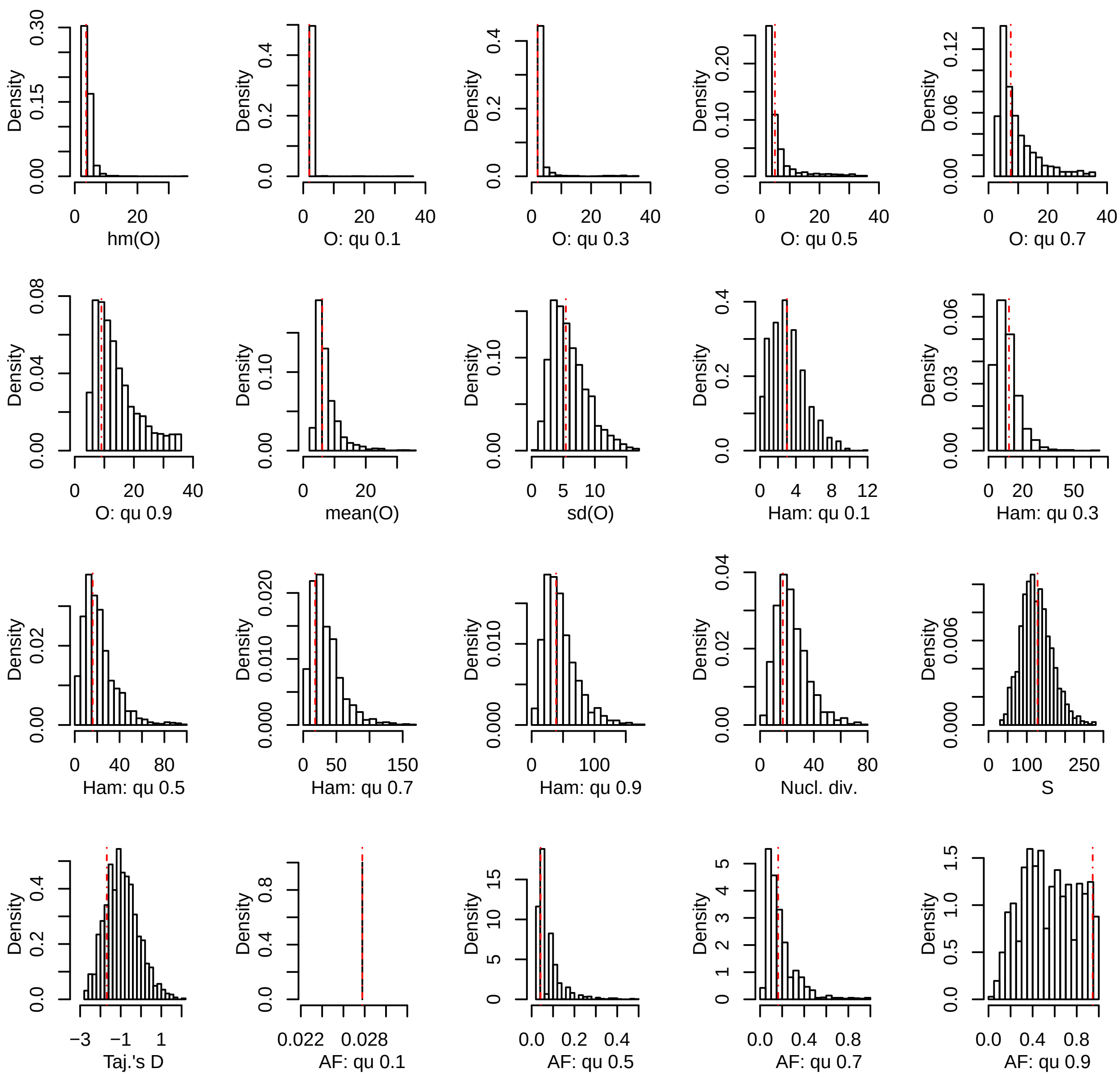


**Supplementary Figure 16.** Posterior predictive check for the data set Lee 2015 clade B. The red lines represent the values for the observed data, the histograms represent the results of 10,000 simulations under the best fitting model (BETA) using the median of the posterior distribution of the parameter α. hm: harmonic mean; qu: quantile; sd: standard deviation; O: minimal observable clade size; Ham: Hamming distance; Nuc. Div.: nucleotide diversity (π); S: number of polymorphic positions; Taj's D: Tajima's D; AF: mutant allele frequency.

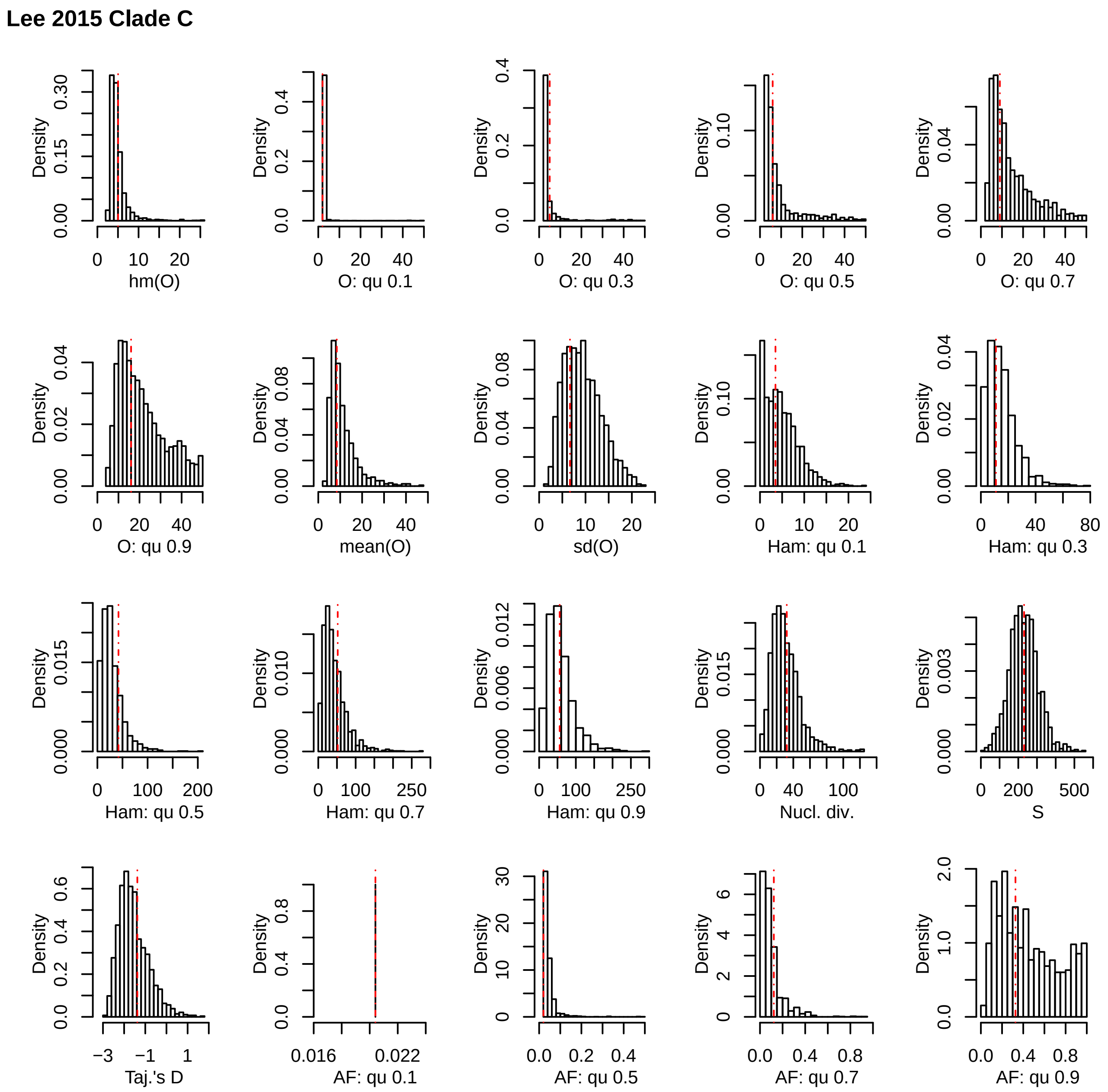


**Supplementary Figure 17.** Posterior predictive check for the data set Lee 2015 clade C. The red lines represent the values for the observed data, the histograms represent the results of 10,000 simulations under the best fitting model (BETA) using the median of the posterior distribution of the parameter α. hm: harmonic mean; qu: quantile; sd: standard deviation; O: minimal observable clade size; Ham: Hamming distance; Nuc. Div.: nucleotide diversity (π); S: number of polymorphic positions; Taj’s D: Tajima’s D; AF: mutant allele frequency.

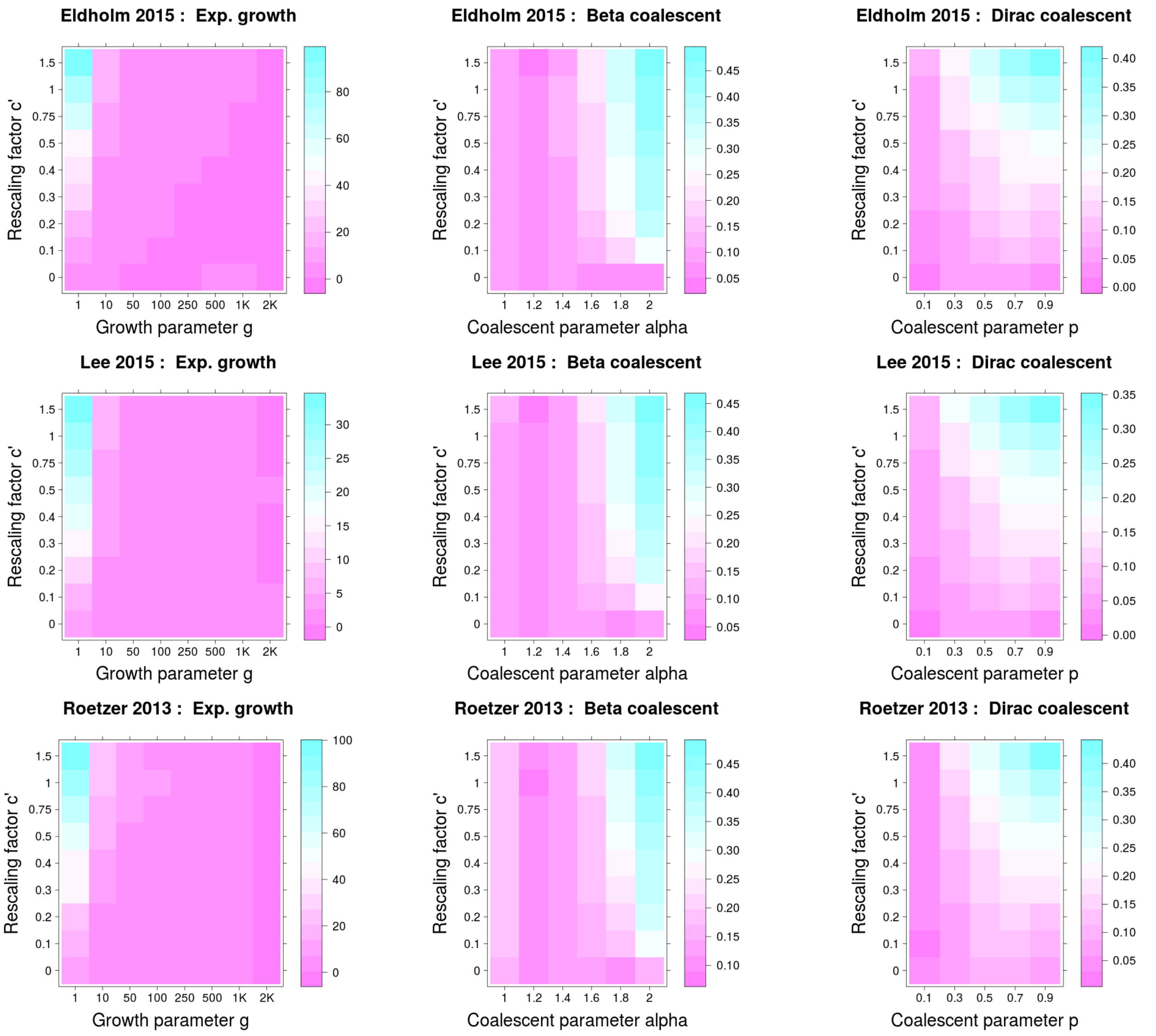


**Supplementary Figure 18.** Mean error of parameter estimation for simulations with serial sampling times, when model selection was performed via ABC using ultrametric tree models. First column (simulations under serially sampled Kingman's coalescent with exponential growth): colors show the absolute error in units of the true parameter, i.e. an error value of 10 corresponds to an average error of 10x the true parameter. Second and third column (serially sampled MMCs): colors show absolute error.

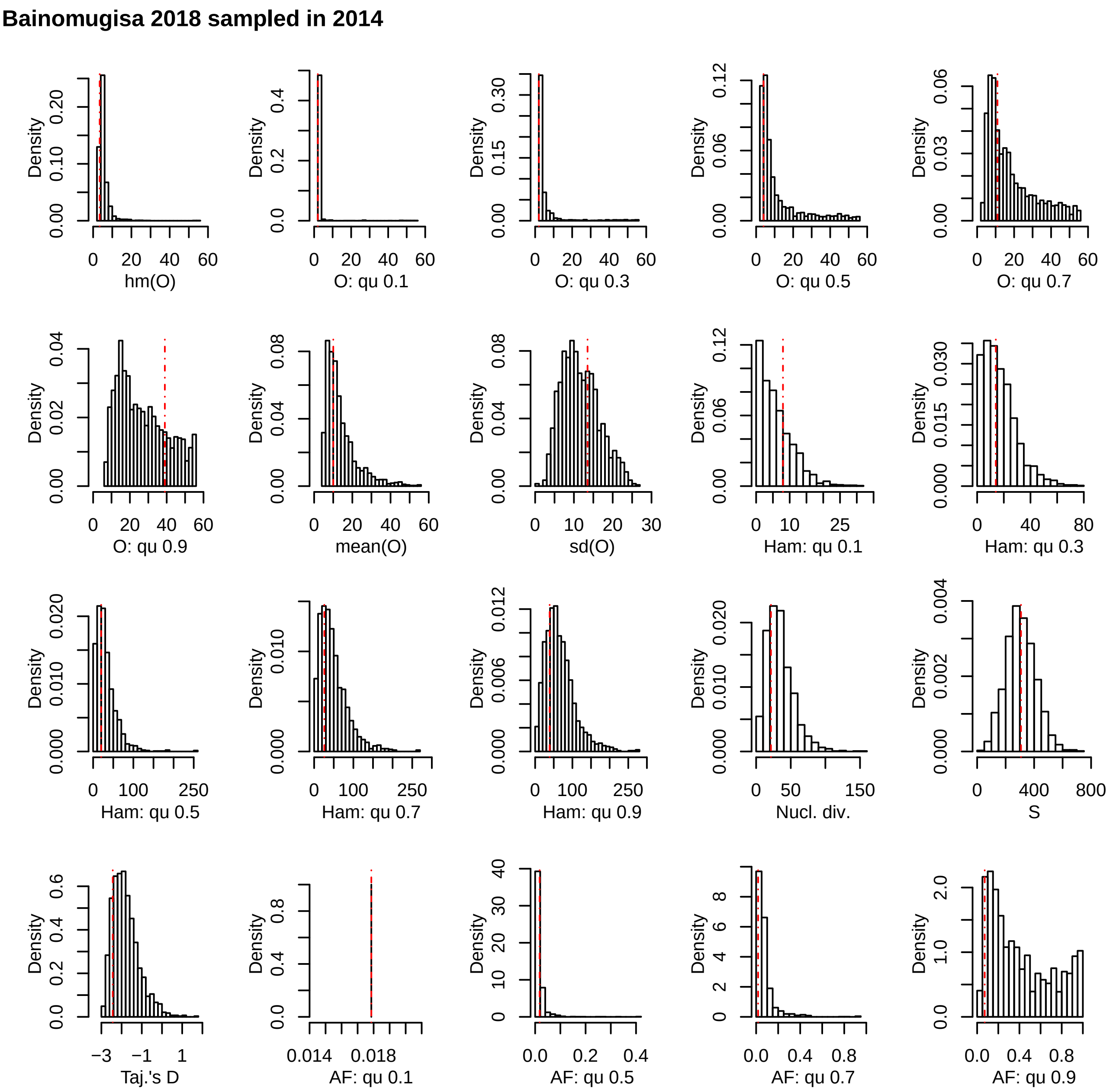


**Supplementary Figure 19.** Posterior predictive check for the data set Bainomugisa 2018 sampled in 2014. The red lines represent the values for the observed data, the histograms represent the results of 10,000 simulations under the best fitting model (BETA) using the median of the posterior distribution of the parameter α. hm: harmonic mean; qu: quantile; sd: standard deviation; O: minimal observable clade size; Ham: Hamming distance; Nuc. Div.: nucleotide diversity (π); S: number of polymorphic positions; Taj's D: Tajima's D; AF: mutant allele frequency.

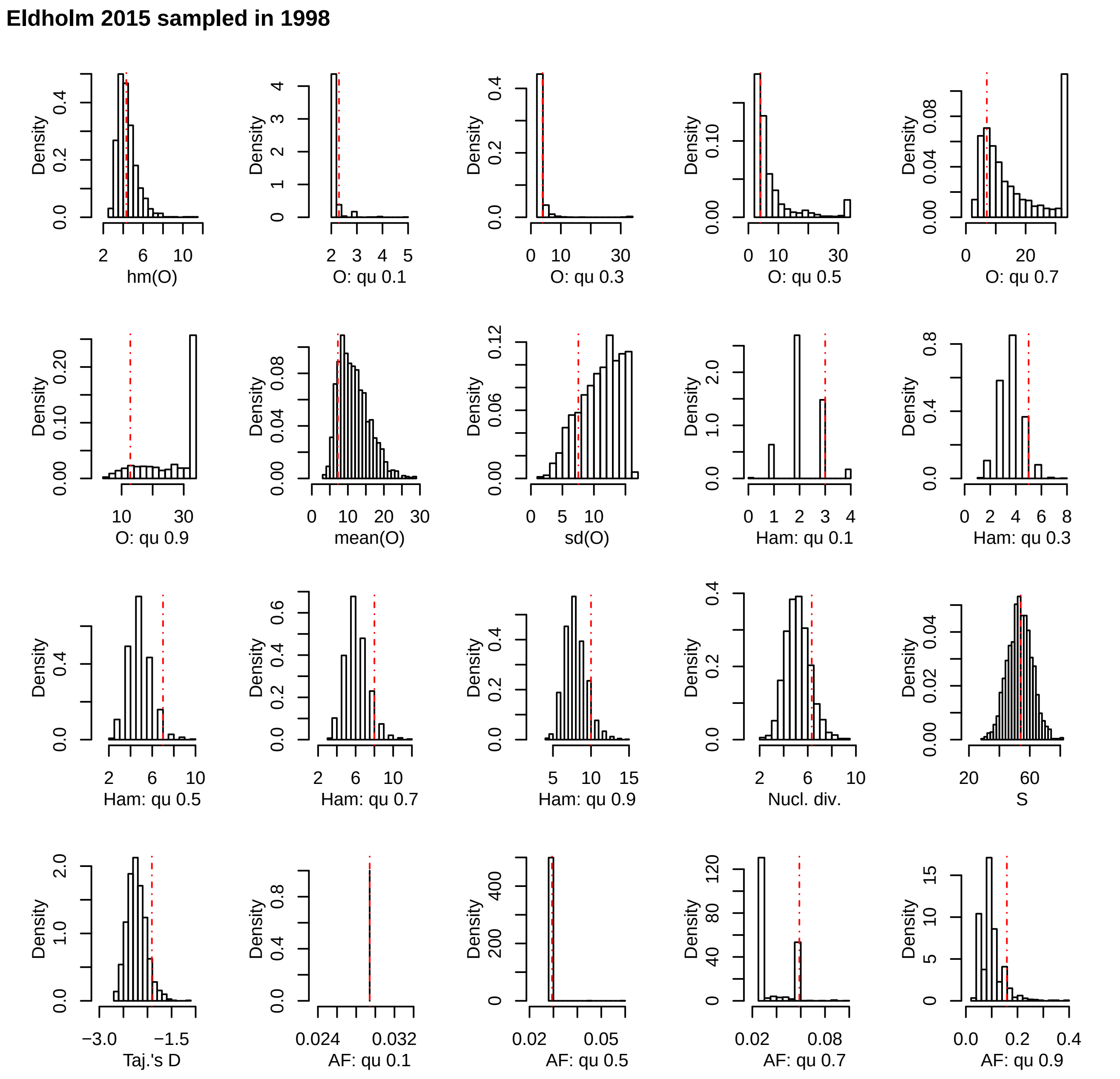


**Supplementary Figure 20.** Posterior predictive check for the data set Eldholm 2015 sampled in 1998. The red lines represent the values for the observed data, the histograms represent the results of 10,000 simulations under the best fitting model (KM+exp) using the median of the posterior distribution of the parameter α. hm: harmonic mean; qu: quantile; sd: standard deviation; O: minimal observable clade size; Ham: Hamming distance; Nuc. Div.: nucleotide diversity (π); S: number of polymorphic positions; Taj's D: Tajima's D; AF: mutant allele frequency.

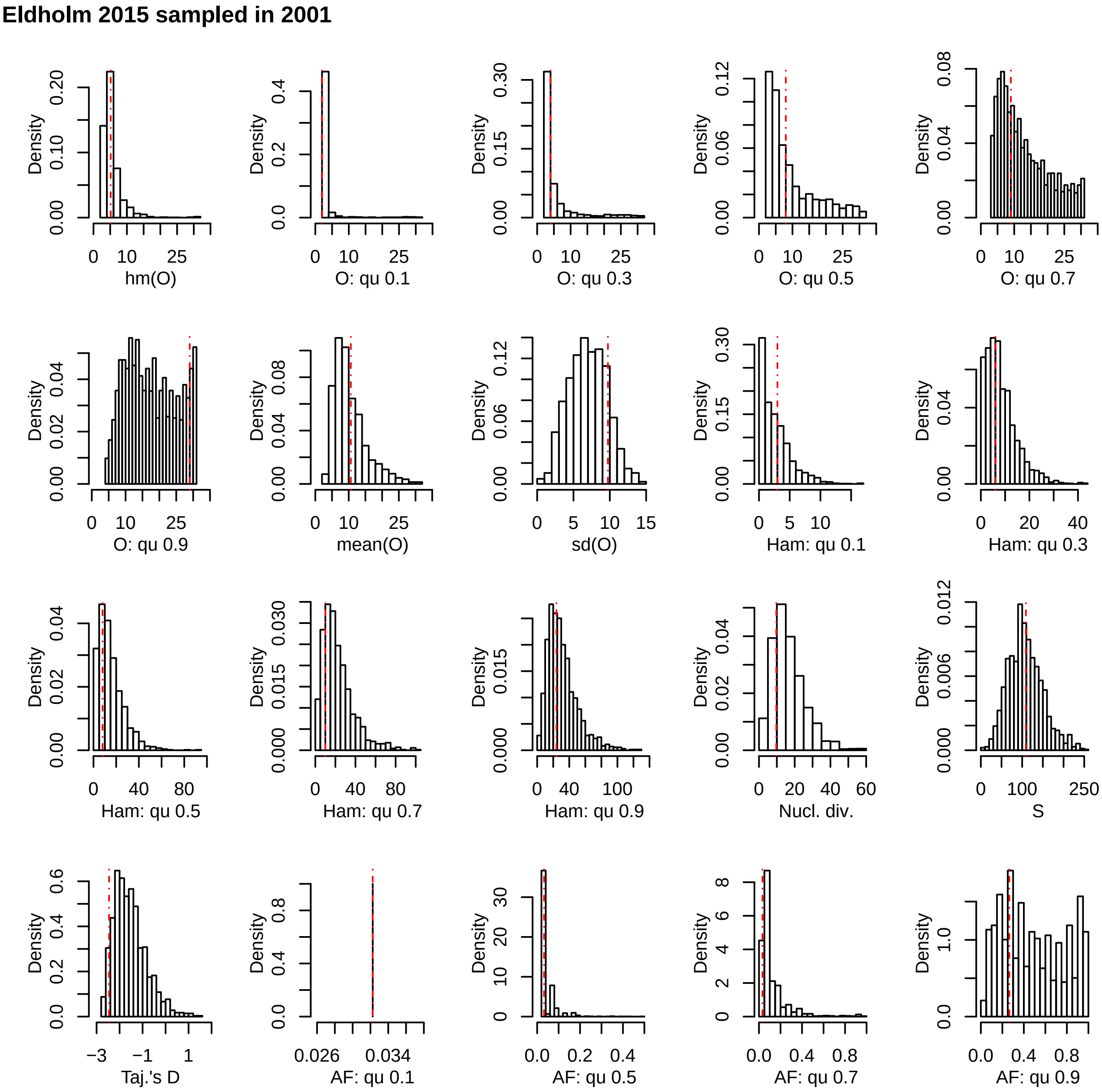


**Supplementary Figure 21.** Posterior predictive check for the data set Eldholm 2015 sampled in 2001. The red lines represent the values for the observed data, the histograms represent the results of 10,000 simulations under the best fitting model (BETA) using the median of the posterior distribution of the parameter α. hm: harmonic mean; qu: quantile; sd: standard deviation; O: minimal observable clade size; Ham: Hamming distance; Nuc. Div.: nucleotide diversity (π); S: number of polymorphic positions; Taj’s D: Tajima’s D; AF: mutant allele frequency.

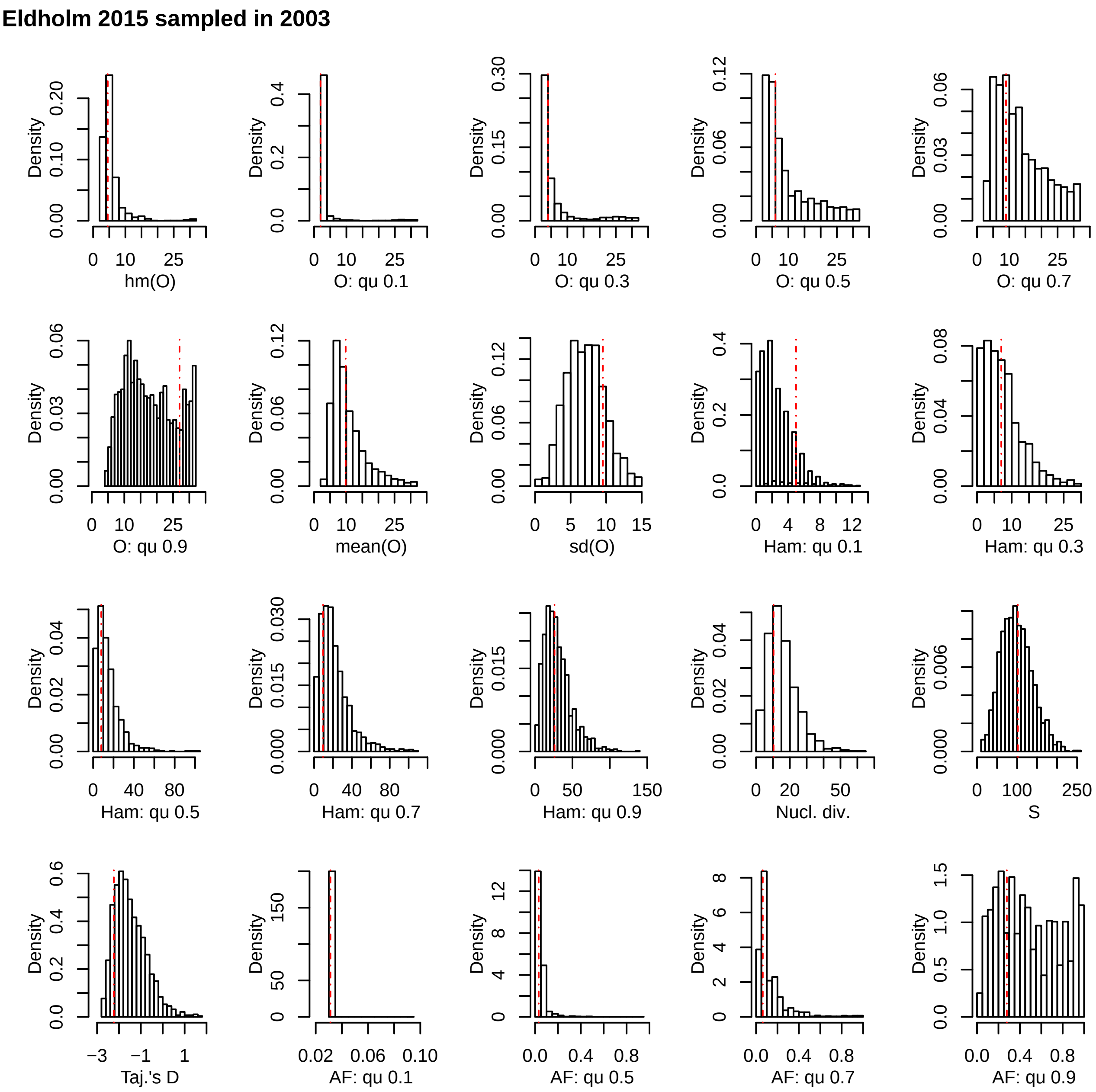


**Supplementary Figure 22.** Posterior predictive check for the data set Eldholm 2015 sampled in 2003. The red lines represent the values for the observed data, the histograms represent the results of 10,000 simulations under the best fitting model (BETA) using the median of the posterior distribution of the parameter α. hm: harmonic mean; qu: quantile; sd: standard deviation; O: minimal observable clade size; Ham: Hamming distance; Nuc. Div.: nucleotide diversity (π); S: number of polymorphic positions; Taj's D: Tajima's D; AF: mutant allele frequency.

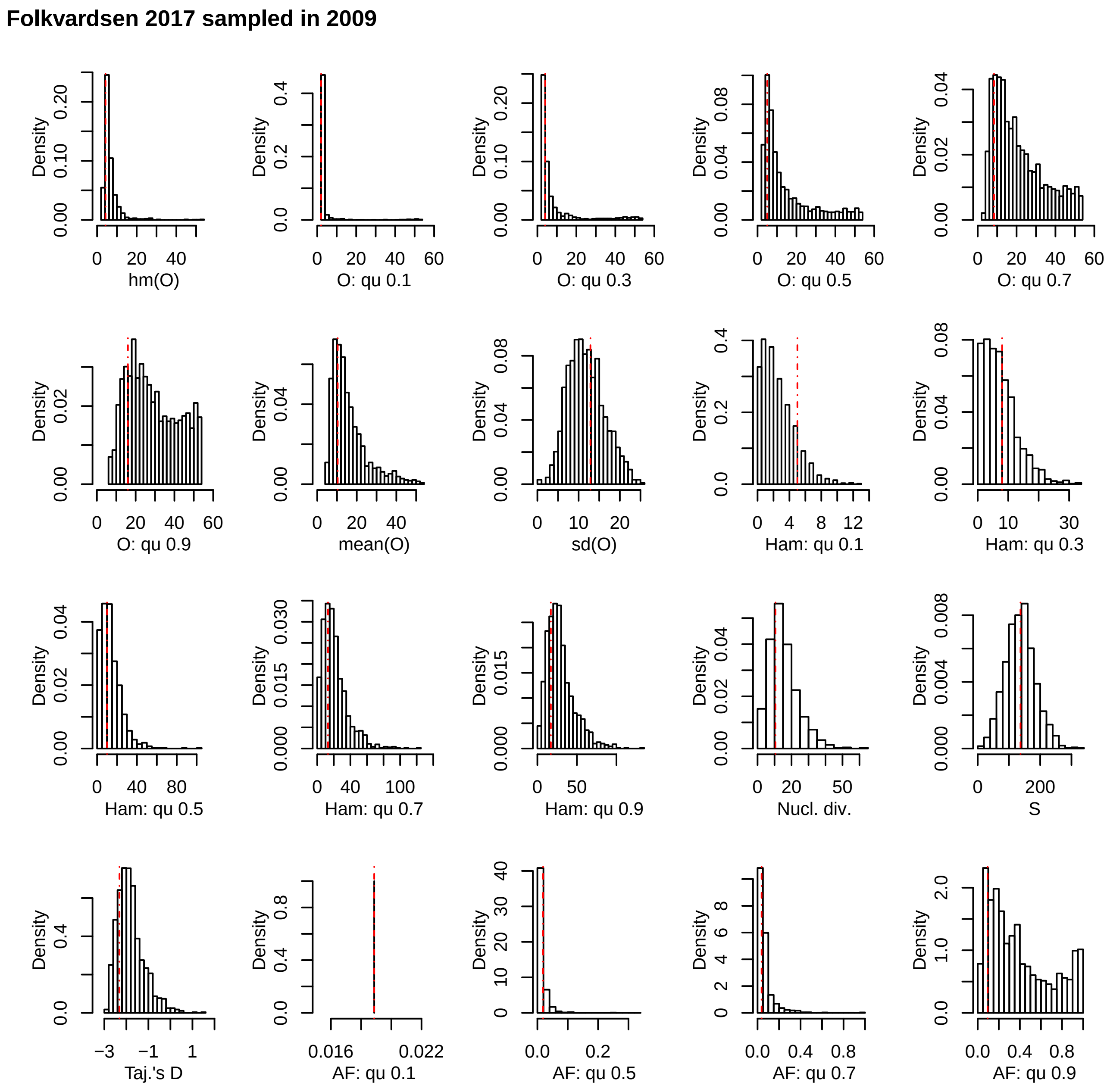


**Supplementary Figure 23.** Posterior predictive check for the data set Folkvardsen 2017 sampled in 2009. The red lines represent the values for the observed data, the histograms represent the results of 10,000 simulations under the best fitting model (BETA) using the median of the posterior distribution of the parameter α. hm: harmonic mean; qu: quantile; sd: standard deviation; O: minimal observable clade size; Ham: Hamming distance; Nuc. Div.: nucleotide diversity (π); S: number of polymorphic positions; Taj's D: Tajima's D; AF: mutant allele frequency.

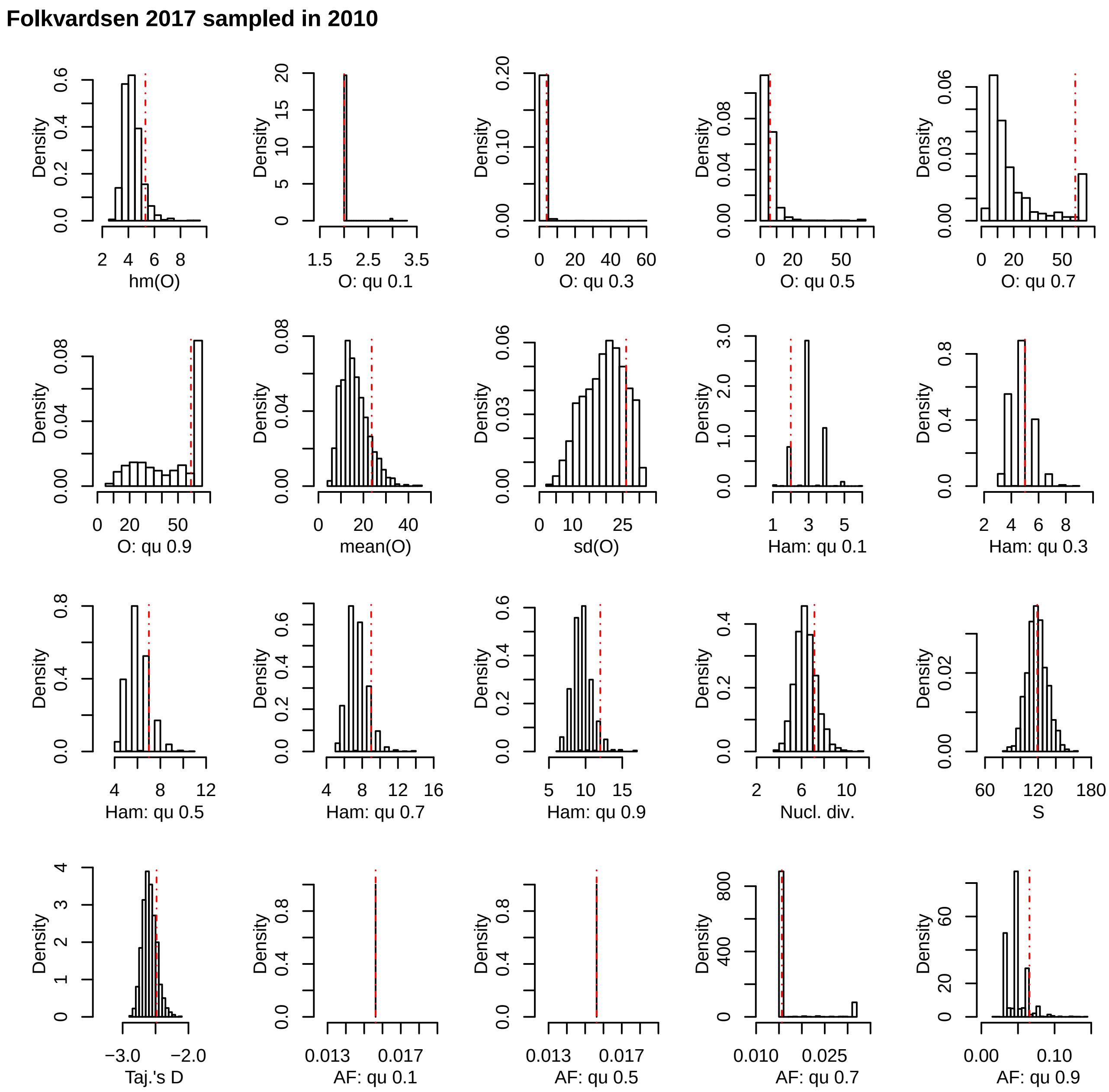


**Supplementary Figure 24.** Posterior predictive check for the data set Folkvardsen 2017 sampled in 2010. The red lines represent the values for the observed data, the histograms represent the results of 10,000 simulations under the best fitting model (BETA) using the median of the posterior distribution of the parameter α. hm: harmonic mean; qu: quantile; sd: standard deviation; O: minimal observable clade size; Ham: Hamming distance; Nuc. Div.: nucleotide diversity (π); S: number of polymorphic positions; Taj's D: Tajima's D; AF: mutant allele frequency.

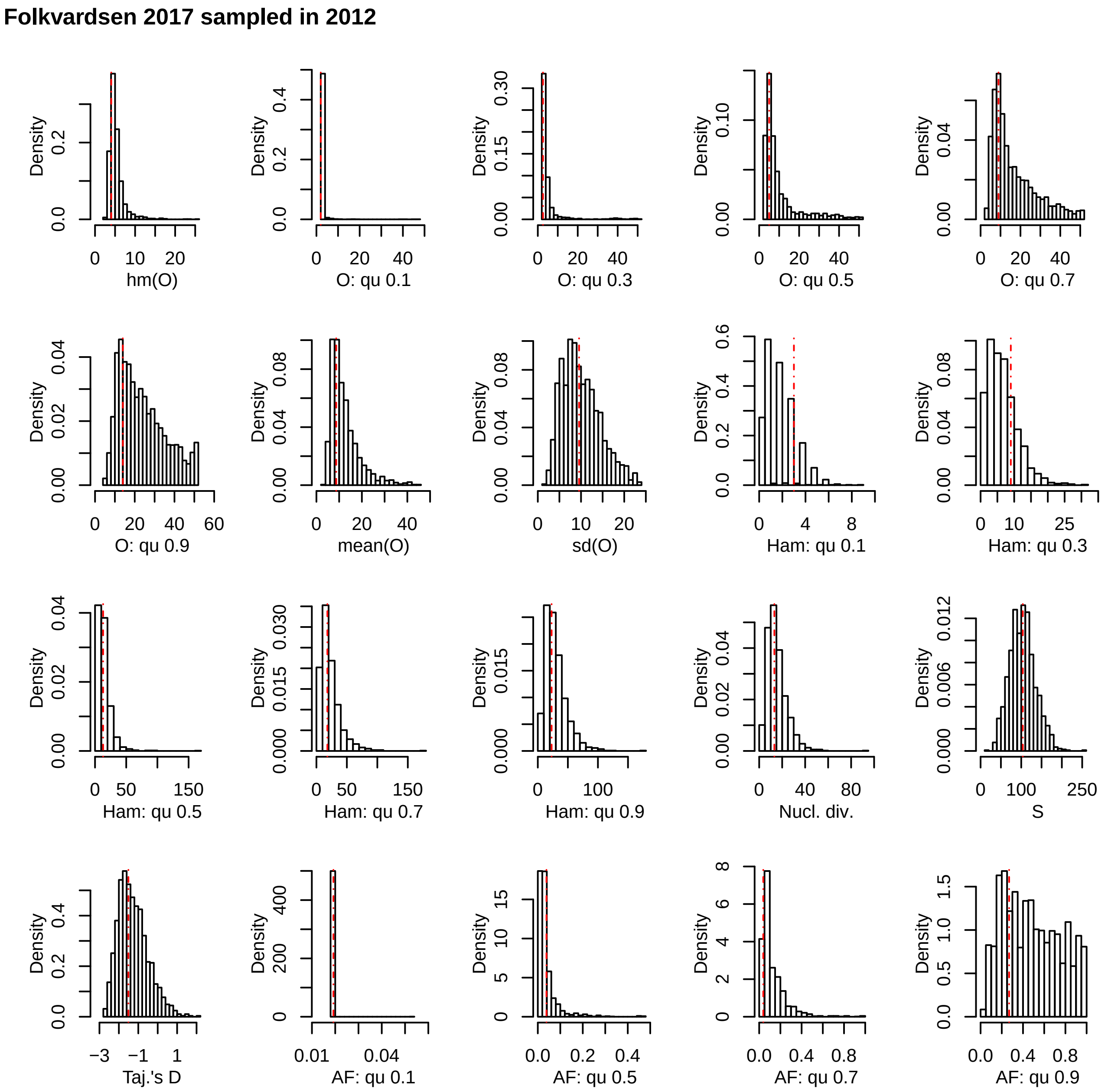


**Supplementary Figure 25.** Posterior predictive check for the data set Folkvardsen 2017 sampled in 2012. The red lines represent the values for the observed data, the histograms represent the results of 10,000 simulations under the best fitting model (BETA) using the median of the posterior distribution of the parameter α. hm: harmonic mean; qu: quantile; sd: standard deviation; O: minimal observable clade size; Ham: Hamming distance; Nuc. Div.: nucleotide diversity (π); S: number of polymorphic positions; Taj’s D: Tajima’s D; AF: mutant allele frequency.

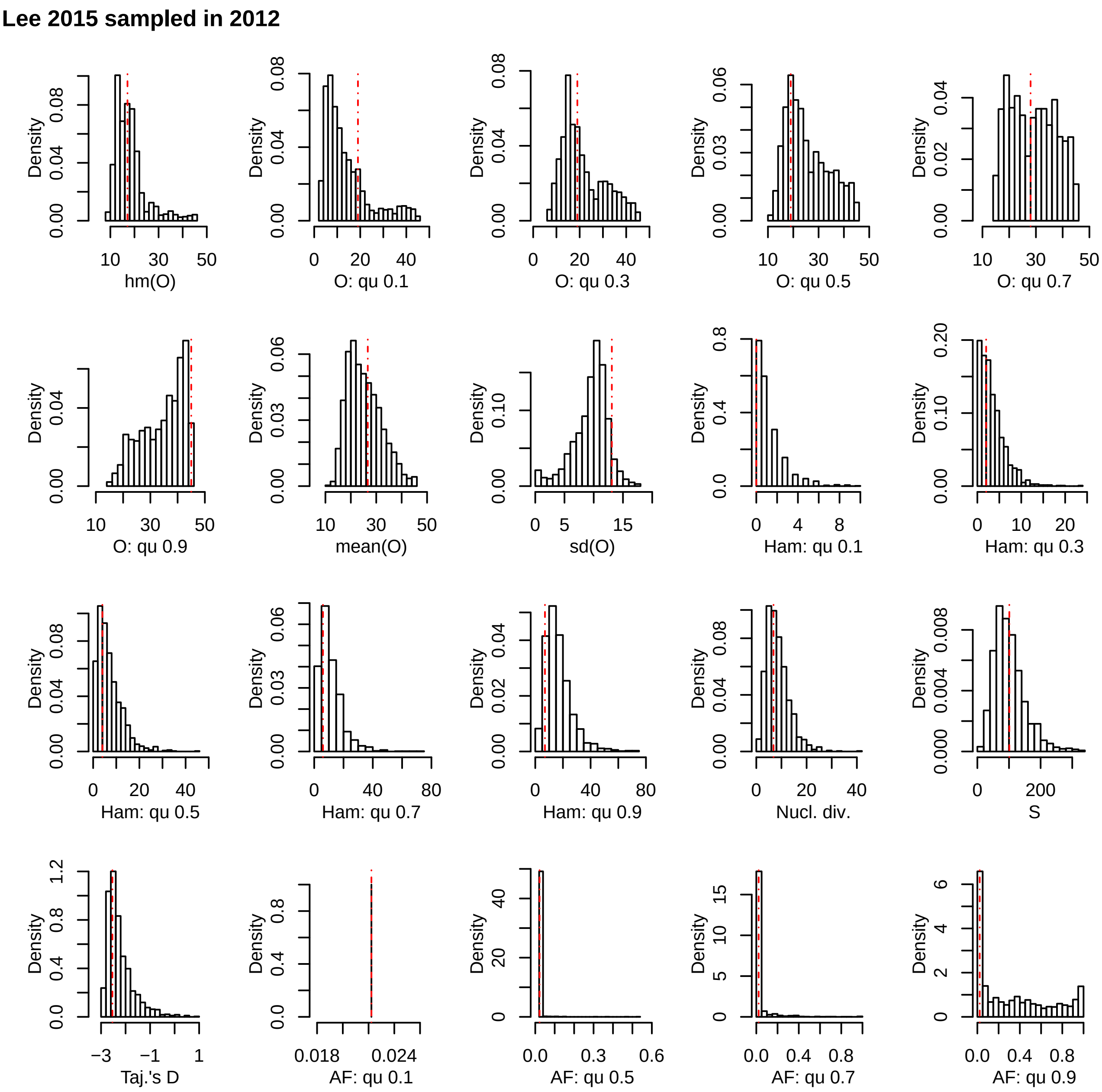


**Supplementary Figure 26**. Posterior predictive check for the data set Lee 2017 sampled in 2012. The red lines represent the values for the observed data, the histograms represent the results of 10,000 simulations under the best fitting model (Dirac) using the median of the posterior distribution of the parameter α. hm: harmonic mean; qu: quantile; sd: standard deviation; O: minimal observable clade size; Ham: Hamming distance; Nuc. Div.: nucleotide diversity (π); S: number of polymorphic positions; Taj's D: Tajima's D; AF: mutant allele frequency.

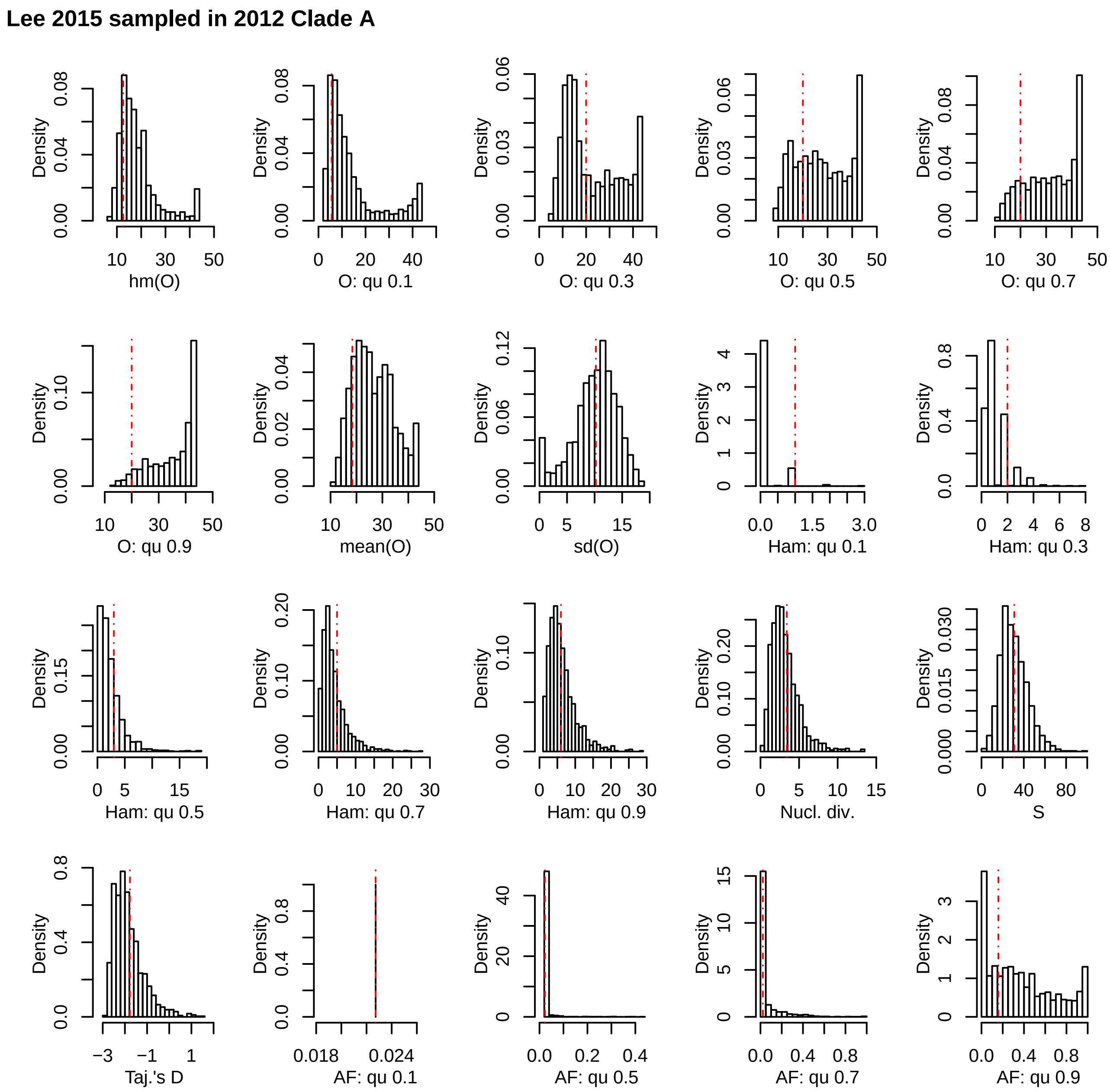


**Supplementary Figure 27**. Posterior predictive check for the data set Lee 2017 sampled in 2012, clade A. The red lines represent the values for the observed data, the histograms represent the results of 10,000 simulations under the best fitting model (Dirac) using the median of the posterior distribution of the parameter α. hm: harmonic mean; qu: quantile; sd: standard deviation; O: minimal observable clade size; Ham: Hamming distance; Nuc. Div.: nucleotide diversity (π); S: number of polymorphic positions; Taj's D: Tajima's D; AF: mutant allele frequency.

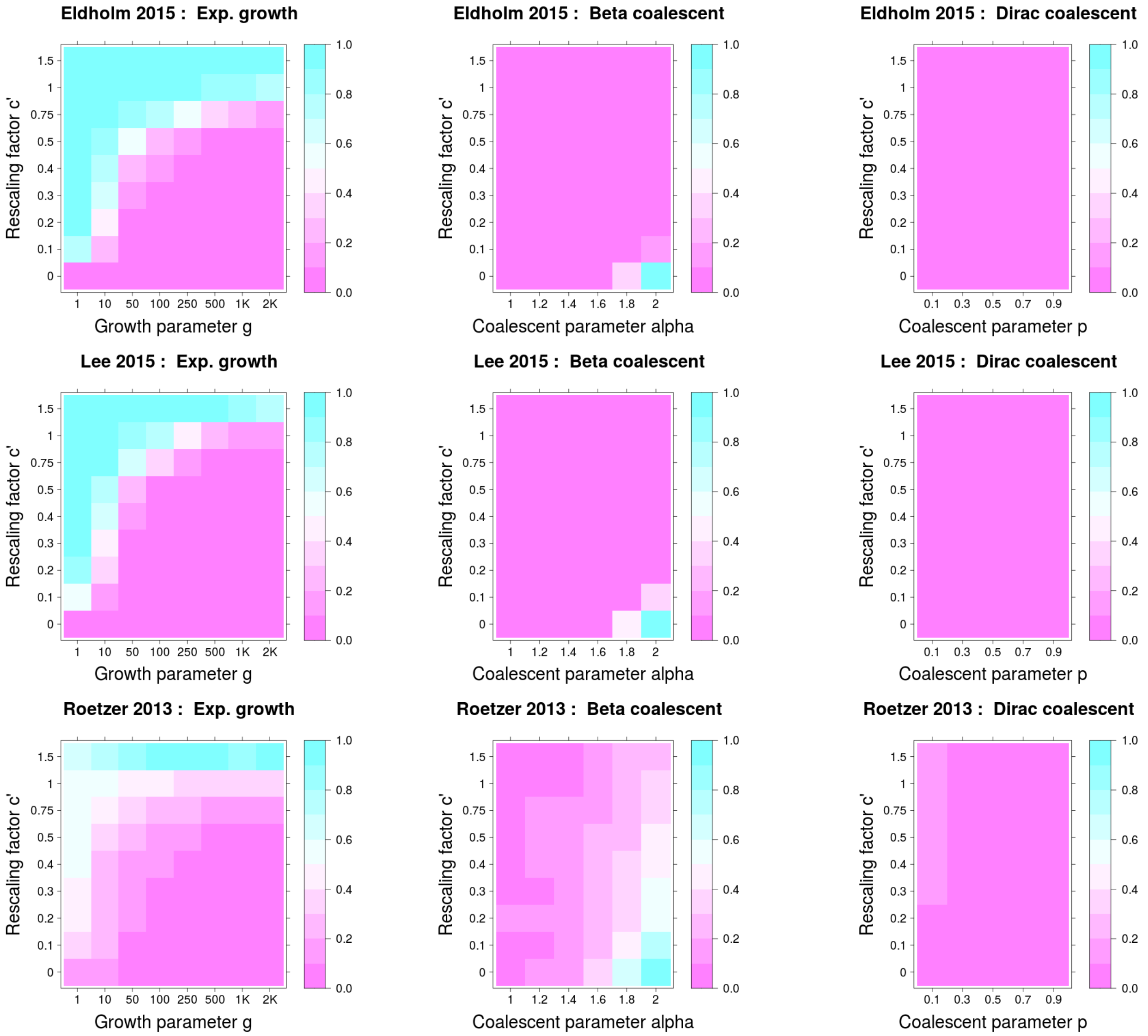


**Supplementary Figure 28**. Proportion of model misidentification for simulations with serial sampling times, when model selection was performed via ABC using ultrametric tree models. The difference from Figure 2 is that we used a logarithmic prior on g for the model KM+exp. Misclassification probabilities are shown as a function of c' (the proportion of the genealogy corresponding to the time period in which samples are collected. i,e the period of sampling spans a time period c'·h, where h is the expected height of the genealogy without serial sampling), and of the parameter of the coalescent models. Misclassification was measured as follow: i) for simulations from serially sampled Kingman's coalescent with exponential growth as being misidentified as either Beta or Dirac (first column) ii) for simulations from serially sampled Beta or Dirac coalescents as being misidentified as Kingman's coalescent with or without exponential growth (second and third columns).

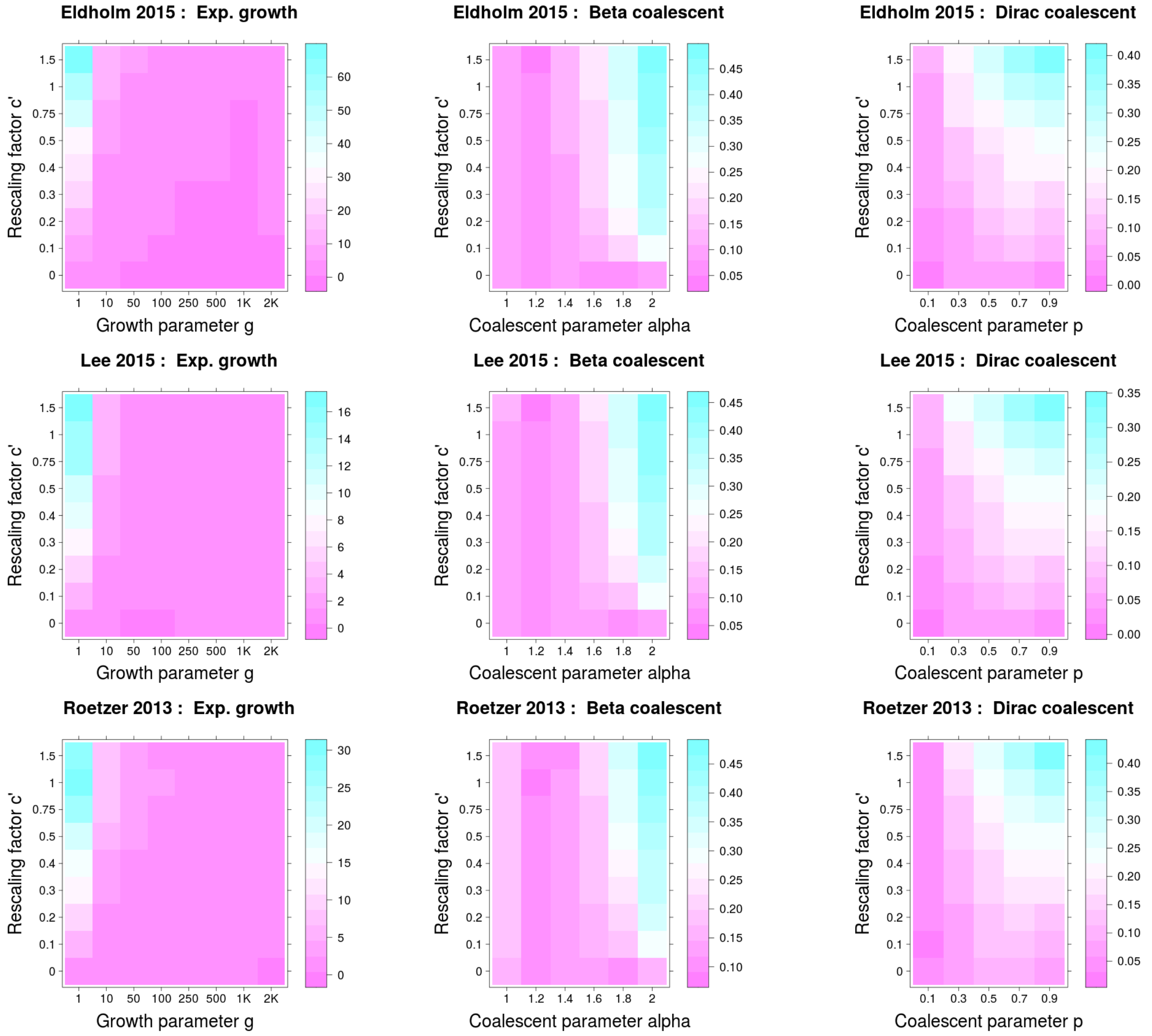


**Supplementary Figure 29.** Mean error of parameter estimation for simulations with serial sampling times, when model selection was performed via ABC using ultrametric tree models. The difference from Sup. Fig. 18 is that we used a logarithmic prior on g for the model KM+exp. First column (simulations under serially sampled Kingman's coalescent with exponential growth): colors show the absolute error in units of the true parameter, i.e. an error value of 10 corresponds to an average error of 10x the true parameter. Second and third column (serially sampled MMCs): colors show absolute error.

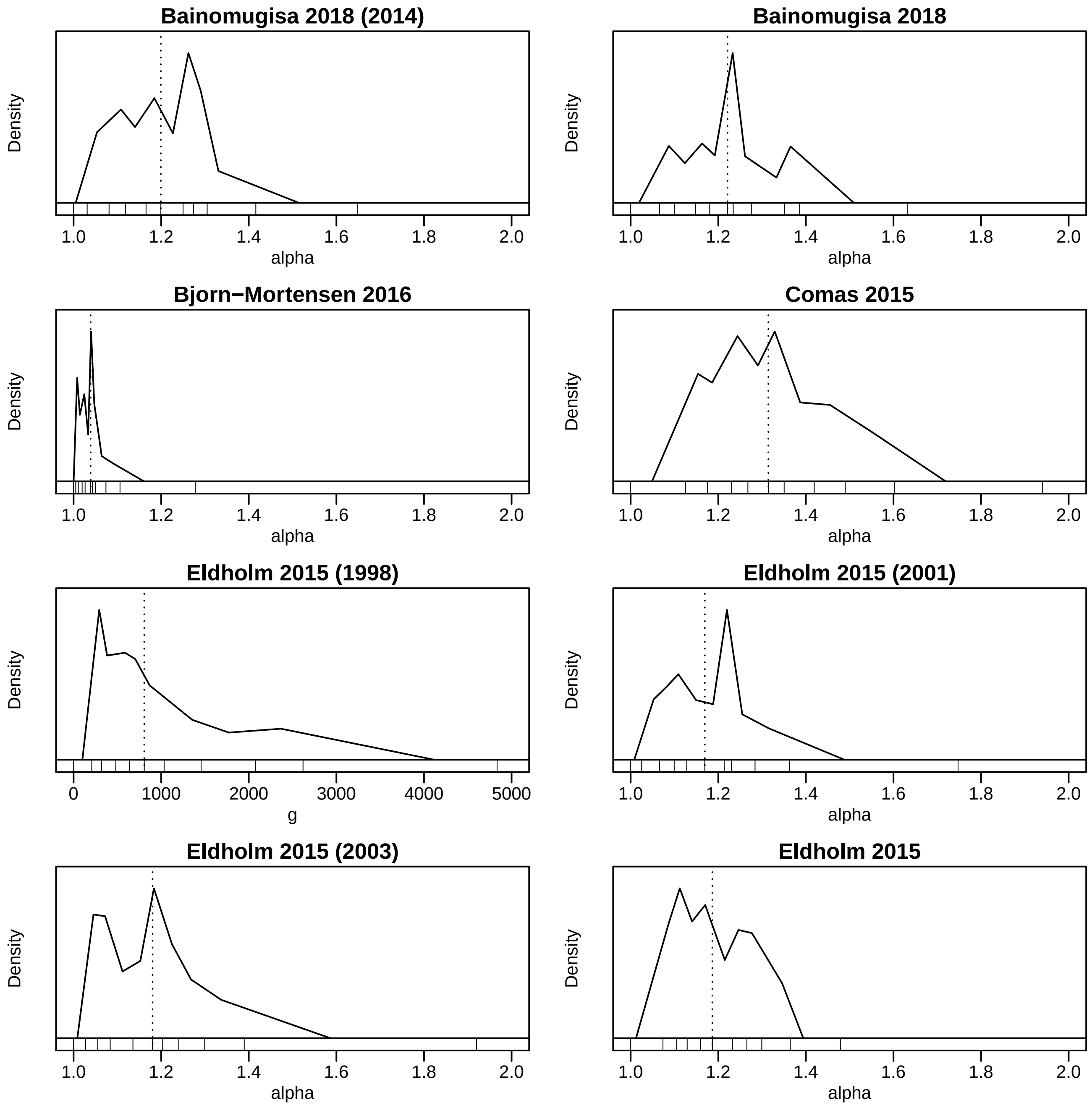


**Supplementary Figures 30-32.** Posterior distribution of the parameter of the best fitting model for all data sets. The plots are approximations of the 95% central posterior density of the parameter obtained from the density of the quantiles resulted from the ABC-RF. The values of the deciles (q 0, q 0.1, q 0.2 … q 0.9, q 1) are plotted at the base of the graph. The script for the plot is available at https://github.com/fabianHOH/mmc_R_gendiv/tree/master/MTB_MMC_repo/plot_posterior_best.R.

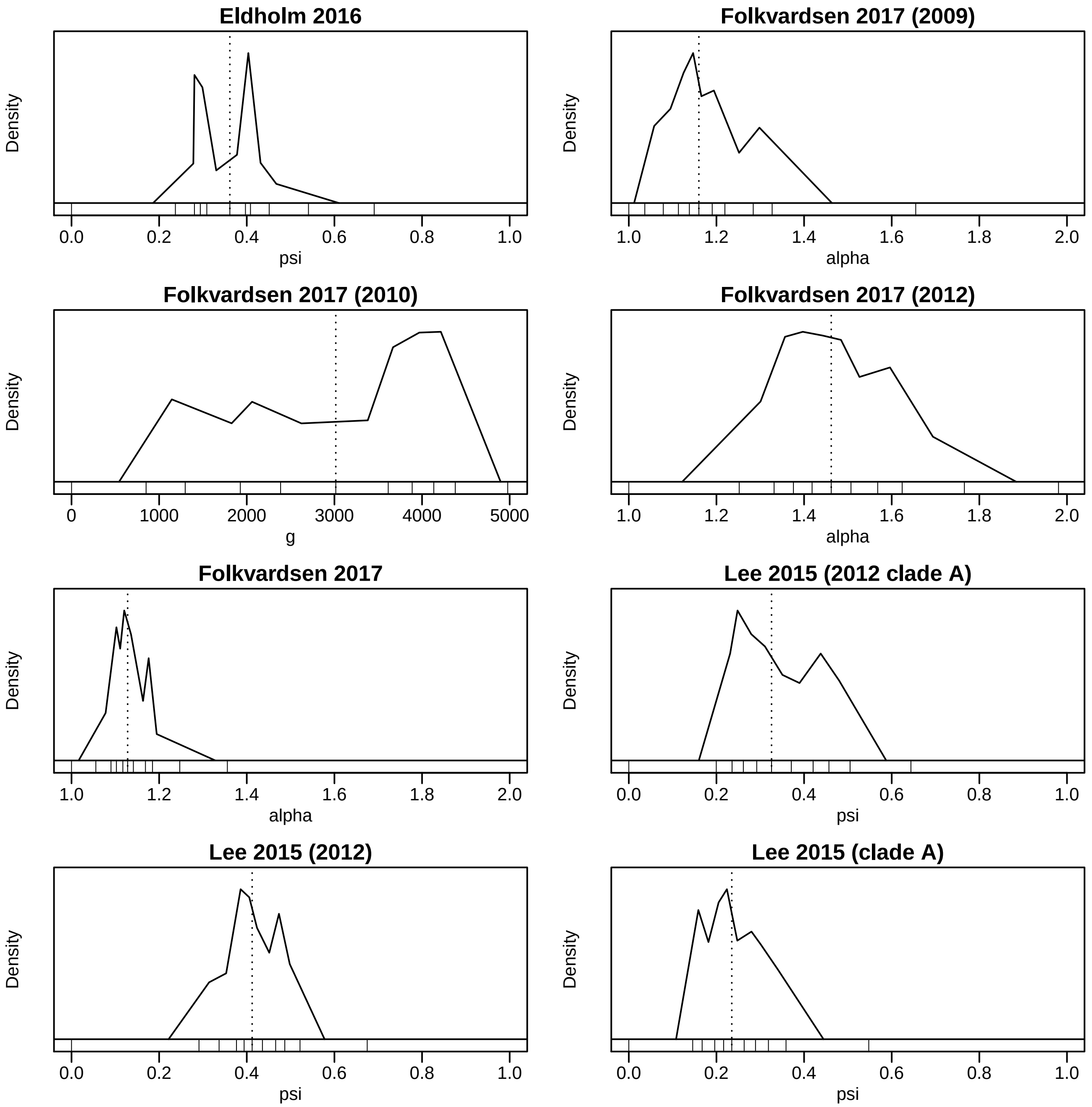


**Supplementary Figures 30-32.** Posterior distribution of the parameter of the best fitting model for all data sets. The plots are approximations of the 95% central posterior density of the parameter obtained from the density of the quantiles resulted from the ABC-RF. The values of the deciles (q 0, q 0.1, q 0.2 … q 0.9, q 1) are plotted at the base of the graph. The script for the plot is available at https://github.com/fabianHOH/mmc_R_gendiv/tree/master/MTB_MMC_repo/plot_posterior_best.R.

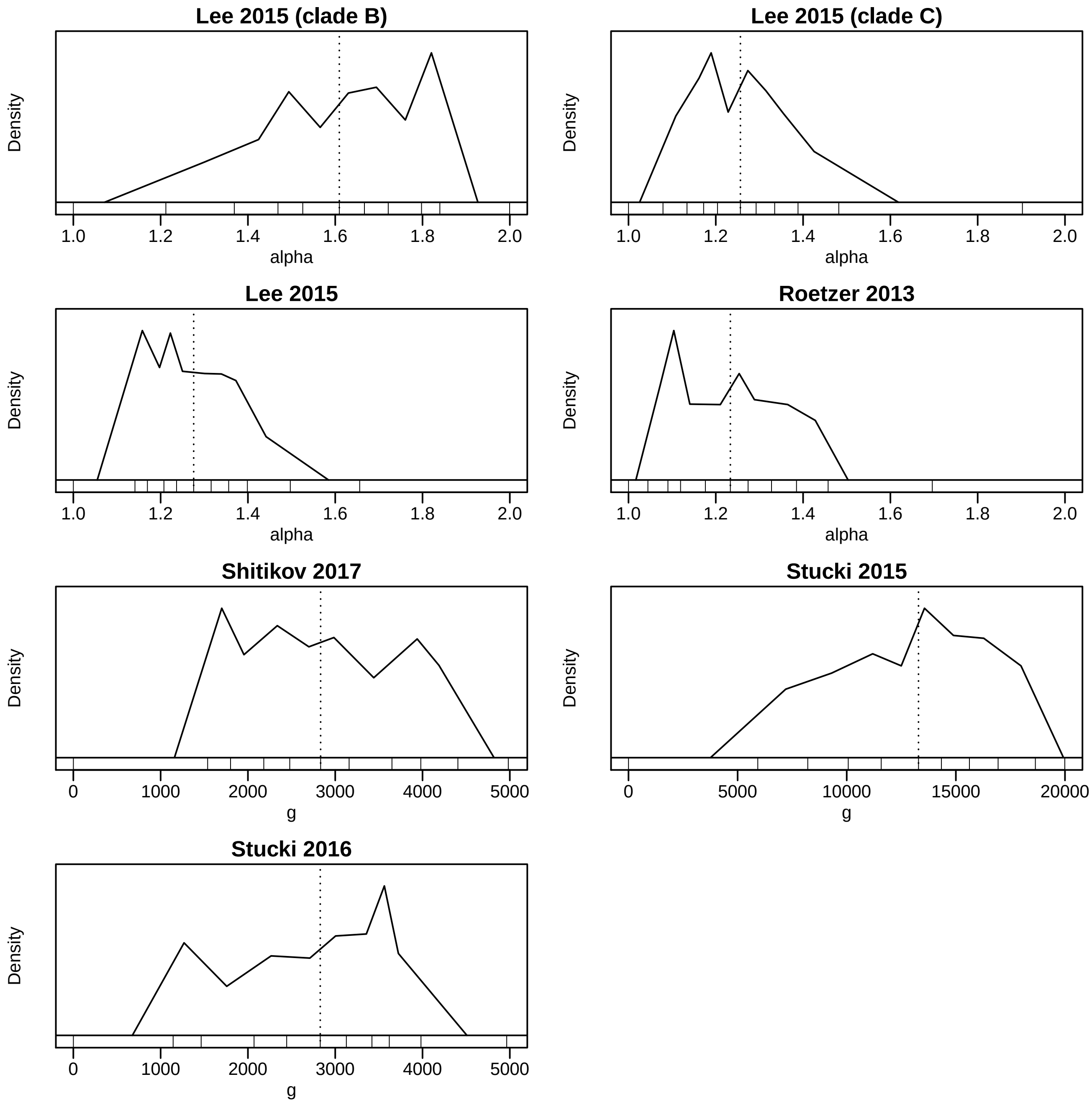


**Supplementary Figures 30-32.** Posterior distribution of the parameter of the best fitting model for all data sets. The plots are approximations of the 95% central posterior density of the parameter obtained from the density of the quantiles resulted from the ABC-RF. The values of the deciles (q 0, q 0.1, q 0.2 … q 0.9, q 1) are plotted at the base of the graph. The script for the plot is available at https://github.com/fabianHOH/mmc_R_gendiv/tree/master/MTB_MMC_repo/plot_posterior_best.R.

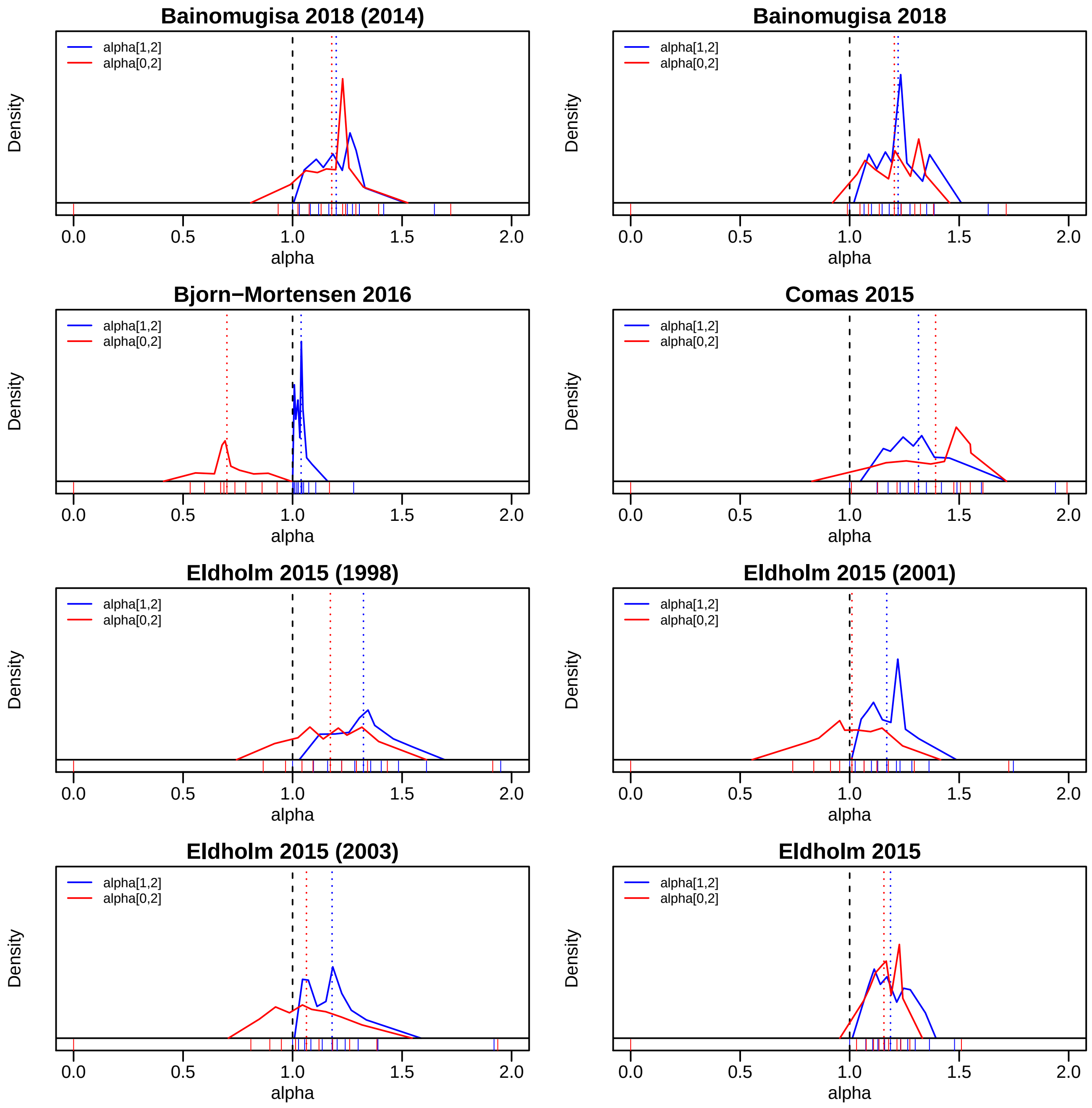


**Supplementary Figure 33-35.** Posterior distribution of the parameter of the Beta coalescent α obtained with two different priors on α: [0,2] in red and [1,2] in blue. The plots are approximations of the 95% central posterior density of the parameter obtained from the density of the quantiles resulted from the ABC-RF. The values of the deciles (q 0, q 0.1, q 0.2 … q 0.9, q 1) are plotted at the base of the graph. The black dashed line indicate α = 1, which correspond to the Bolthausen-Sznitman coalescent. The script for the plot is available at https://github.com/fabianHOH/mmc_R_gendiv/tree/master/MTB_MMC_repo/plot_posterior_alpha.R.

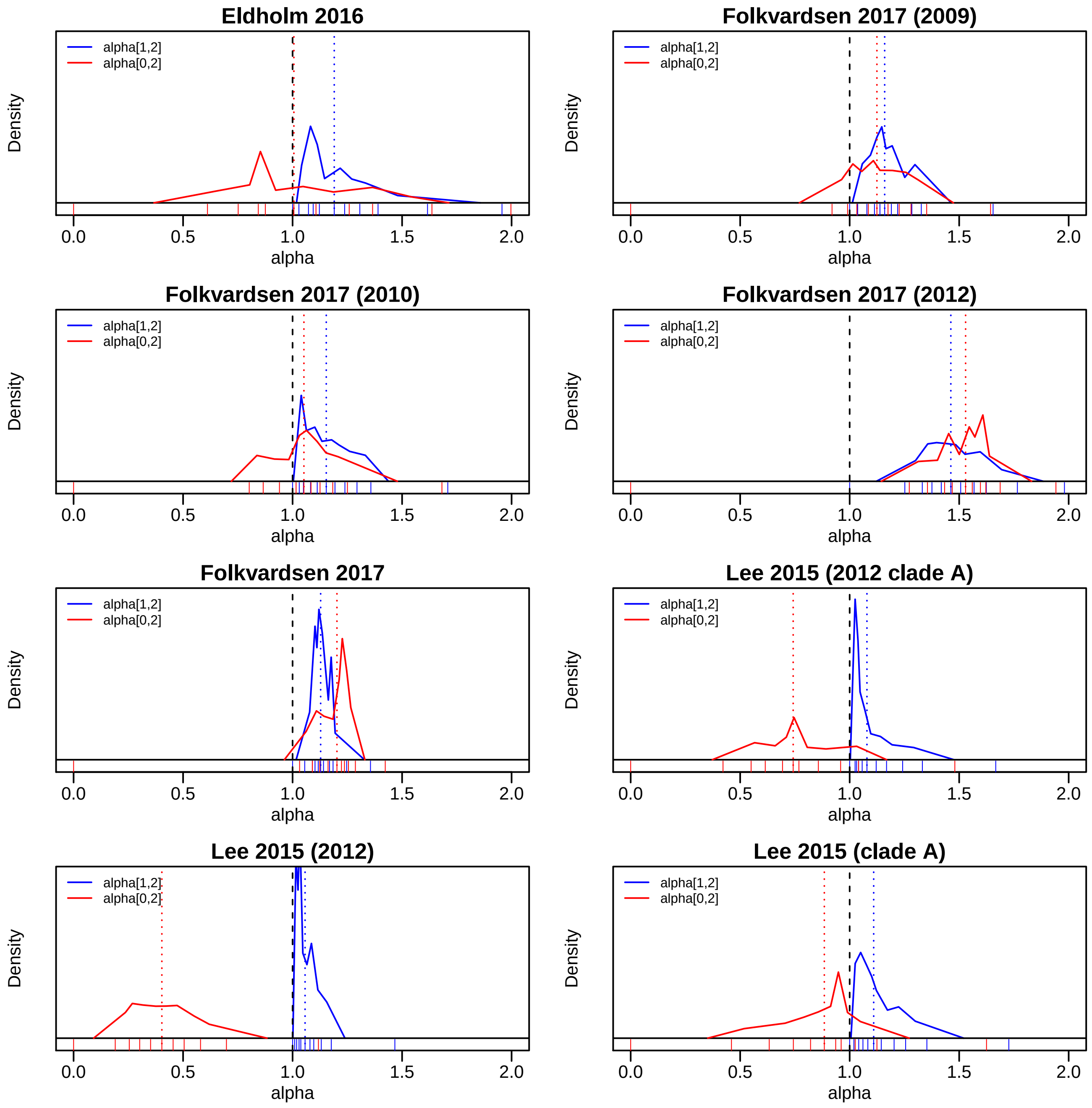


**Supplementary Figure 33-35.** Posterior distribution of the parameter of the Beta coalescent α obtained with two different priors on α: [0,2] in red and [1,2] in blue. The plots are approximations of the 95% central posterior density of the parameter obtained from the density of the quantiles resulted from the ABC-RF. The values of the deciles (q 0, q 0.1, q 0.2 … q 0.9, q 1) are plotted at the base of the graph. The black dashed line indicate α = 1, which correspond to the Bolthausen-Sznitman coalescent. The script for the plot is available at https://github.com/fabianHOH/mmc_R_gendiv/tree/master/MTB_MMC_repo/plot_posterior_alpha.R.

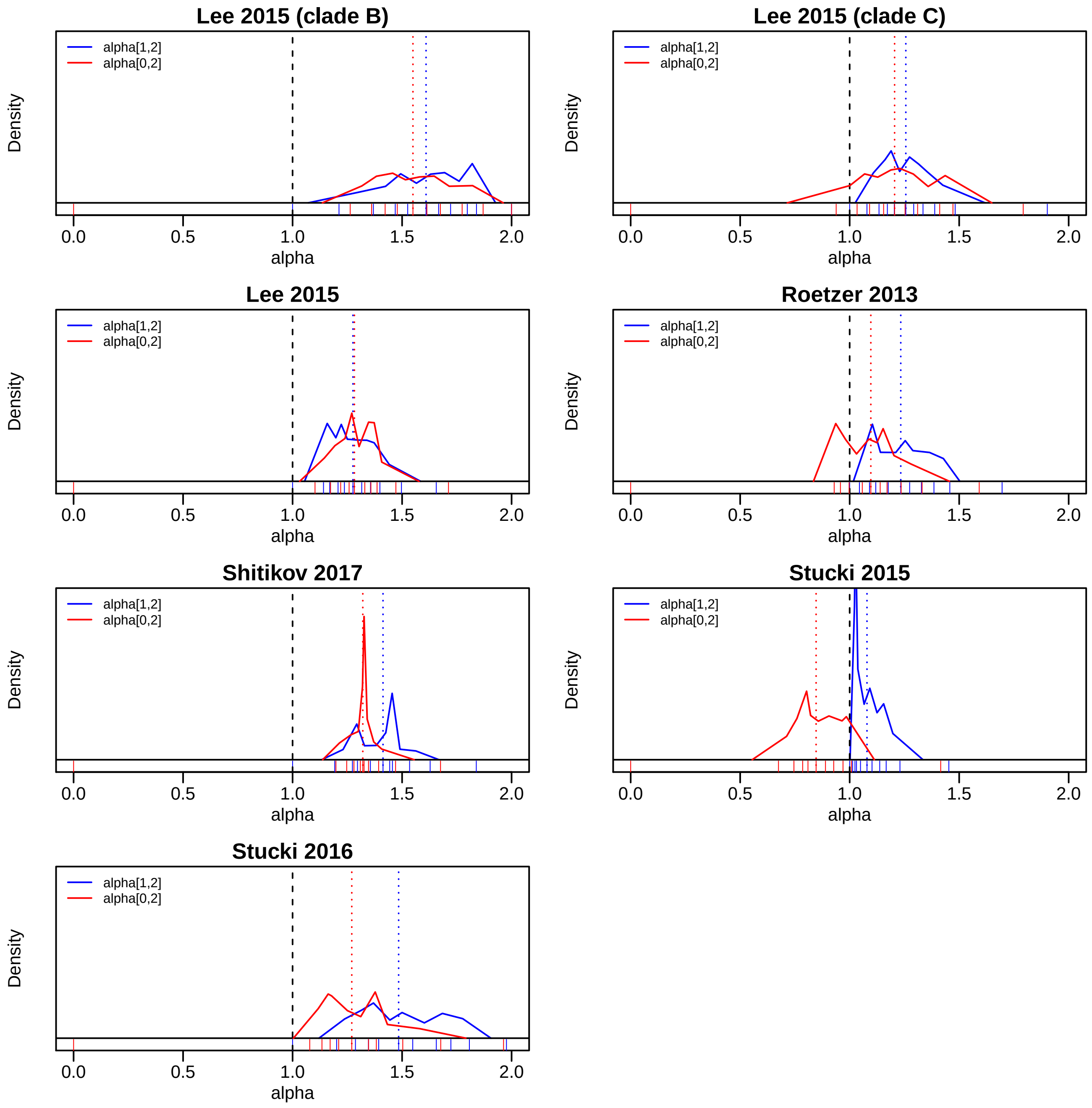


**Supplementary Figure 33-35.** Posterior distribution of the parameter of the Beta coalescent α obtained with two different priors on α: [0,2] in red and [1,2] in blue. The plots are approximations of the 95% central posterior density of the parameter obtained from the density of the quantiles resulted from the ABC-RF. The values of the deciles (q 0, q 0.1, q 0.2 … q 0.9, q 1) are plotted at the base of the graph. The black dashed line indicate α = 1, which correspond to the Bolthausen-Sznitman coalescent. The script for the plot is available at https://github.com/fabianHOH/mmc_R_gendiv/tree/master/MTB_MMC_repo/plot_posterior_alpha.R.

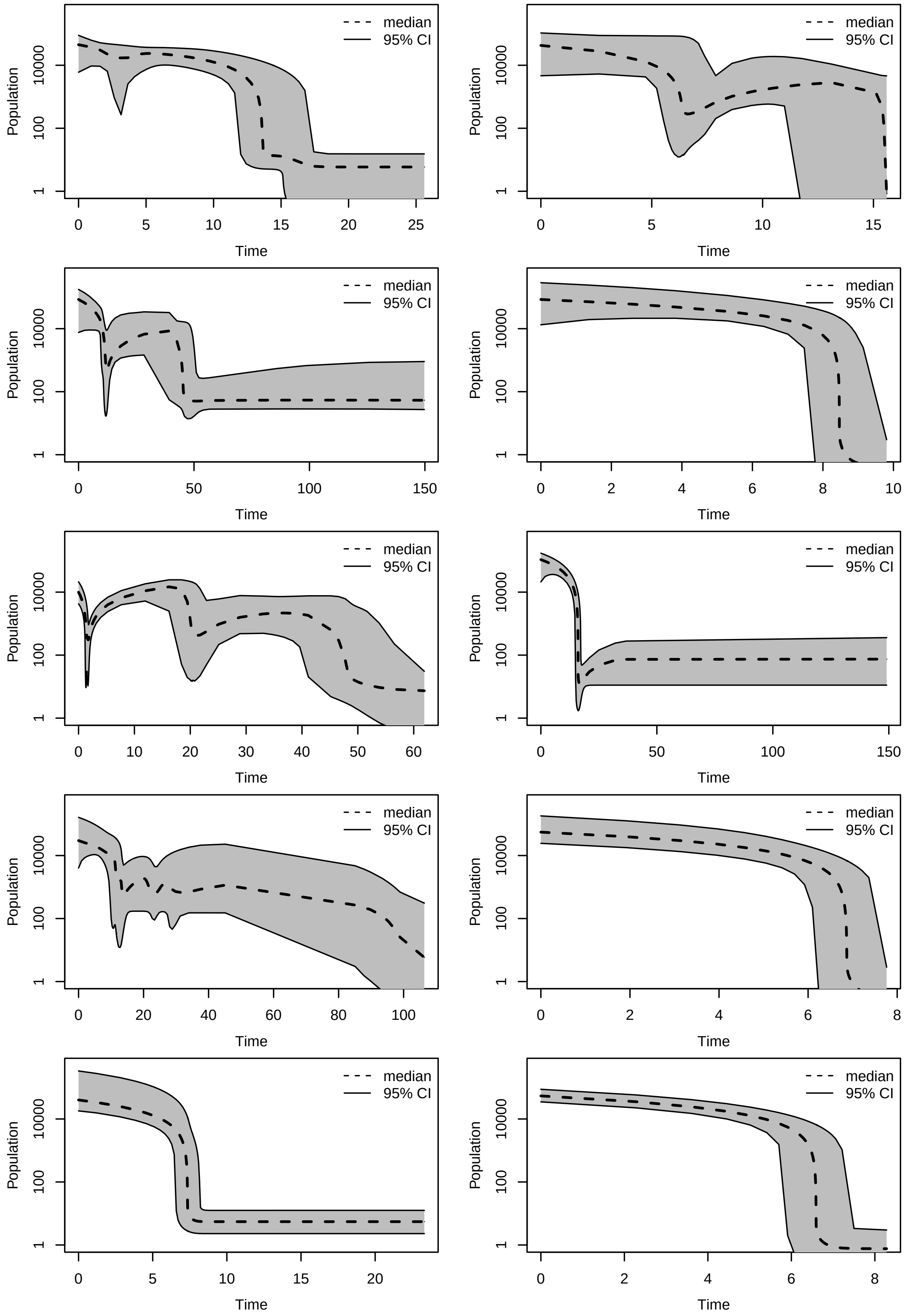


**Supplementary Figure 36.** Bayesian skyline plots of ten different data sets simulated with constant effective population size and α = 0.5. On the y axis the inferred effective population size, on the x axis the time in years before sampling.

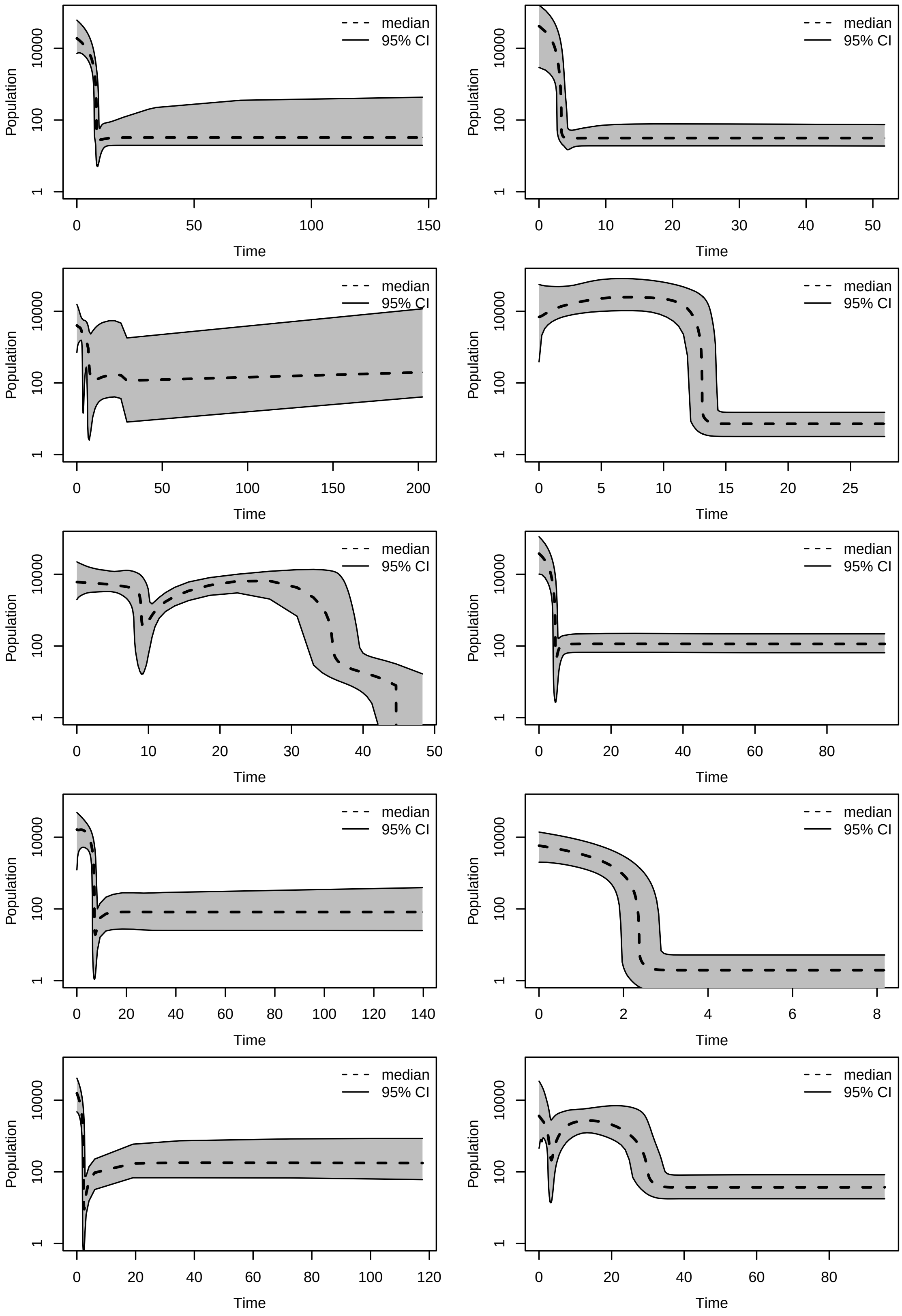


**Supplementary Figure 37.** Bayesian skyline plots of ten different data sets simulated with constant effective population size and α = 0.75. On the y axis the inferred effective population size, on the x axis the time in years before sampling.

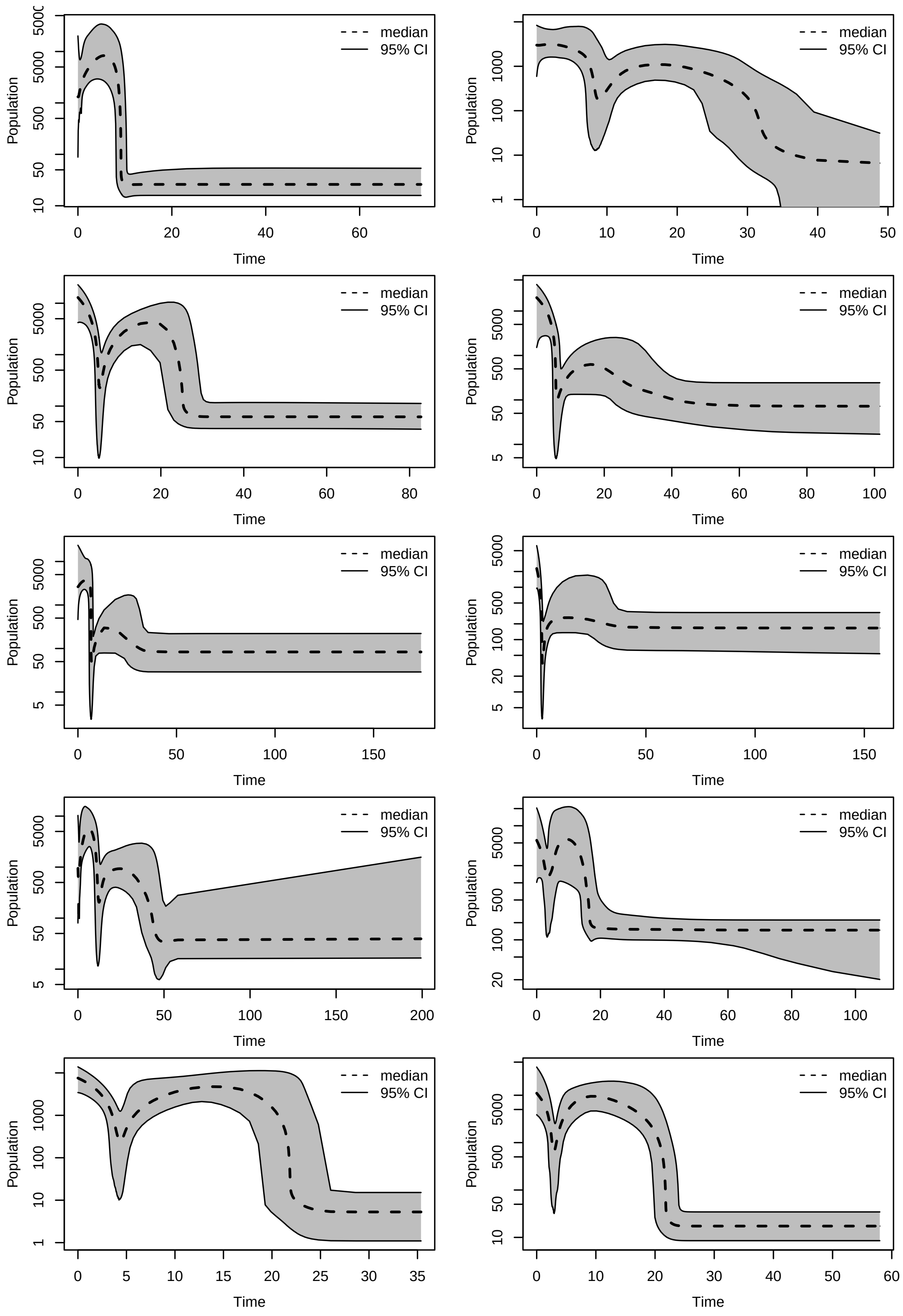


**Supplementary Figure 38.** Bayesian skyline plots of ten different data sets simulated with constant effective population size and α = 1. On the y axis the inferred effective population size, on the x axis the time in years before sampling.

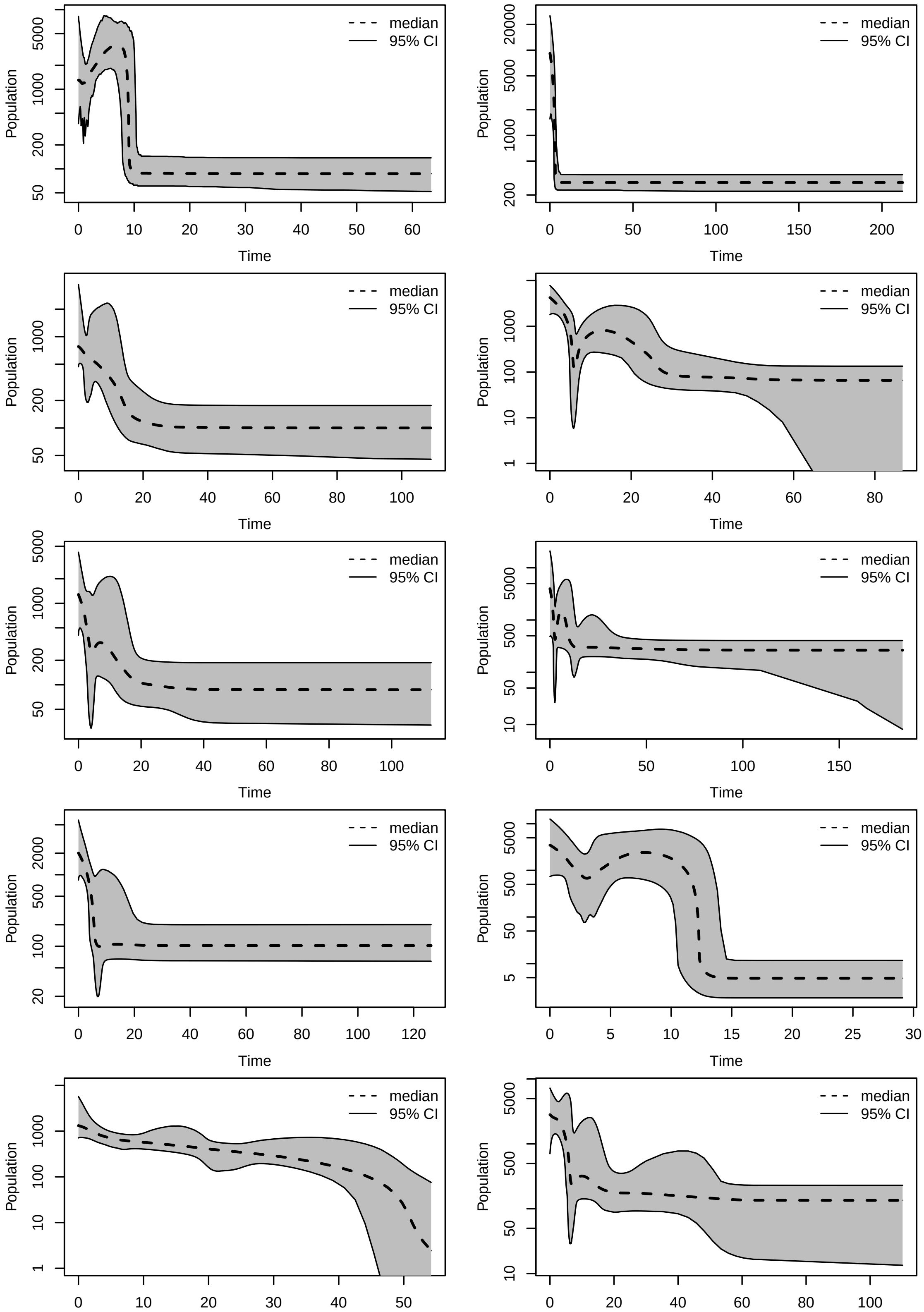


**Supplementary Figure 39.** Bayesian skyline plots of ten different data sets simulated with constant effective population size and α = 1.25. On the y axis the inferred effective population size, on the x axis the time in years before sampling.

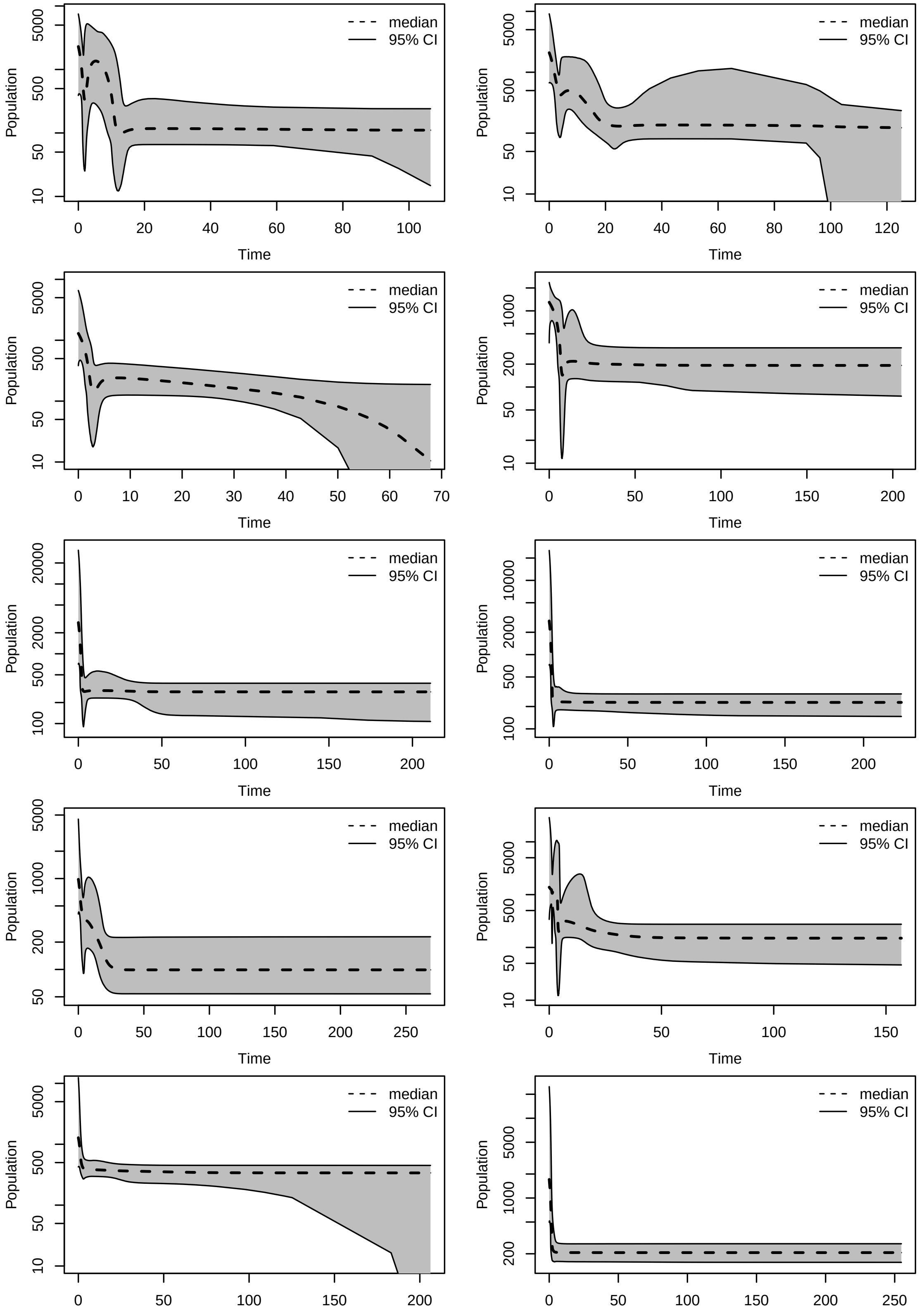


**Supplementary Figure 40.** Bayesian skyline plots of ten different data sets simulated with constant effective population size and α = 1.5. On the y axis the inferred effective population size, on the x axis the time in years before sampling.

Bainomugisa 2018

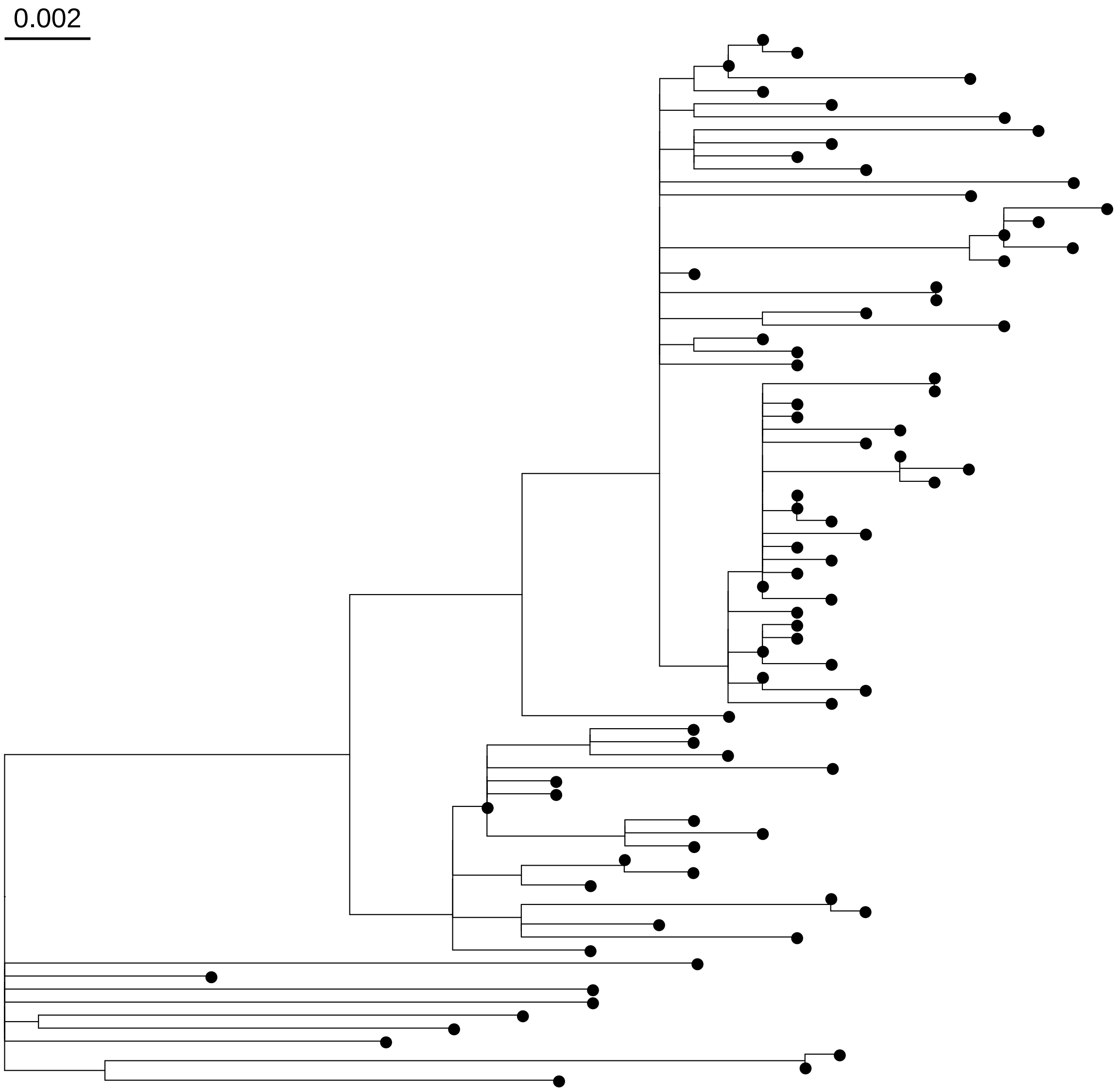


**Supplementary Figure 41.** Phylogenetic tree for the data set Bainomugisa 2018, the scale in expected nucleotide changes per polymorphic site. The tree was rooted with the reconstructed ancestral genome of the *Mycobacterium tuberculosis* complex (not depicted in the tree).

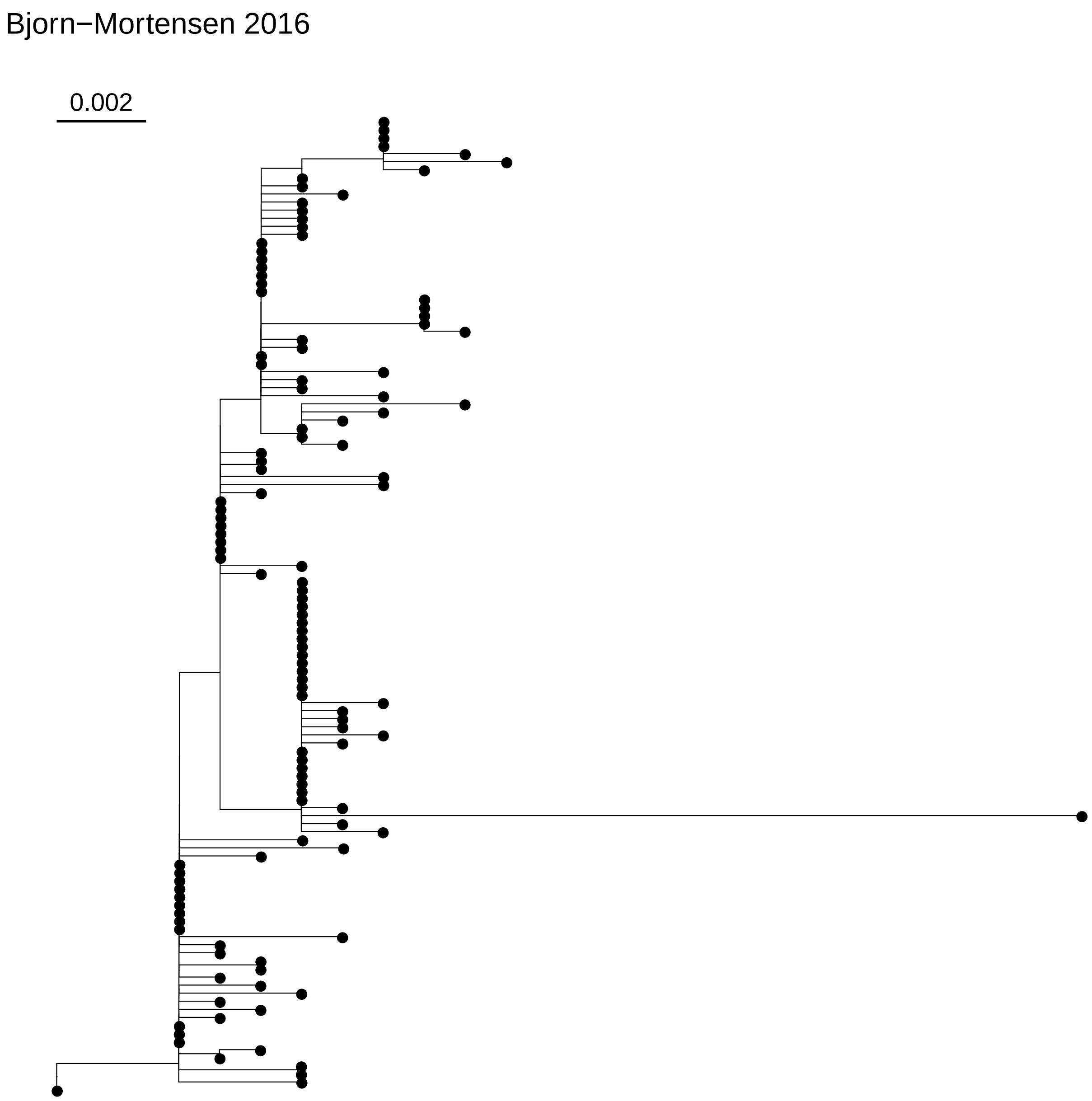


**Supplementary Figure 42.** Phylogenetic tree for the data set Bjorn-Mortensen 2016, the scale in expected nucleotide changes per polymorphic site. The tree was rooted with the reconstructed ancestral genome of the *Mycobacterium tuberculosis* complex (not depicted in the tree).

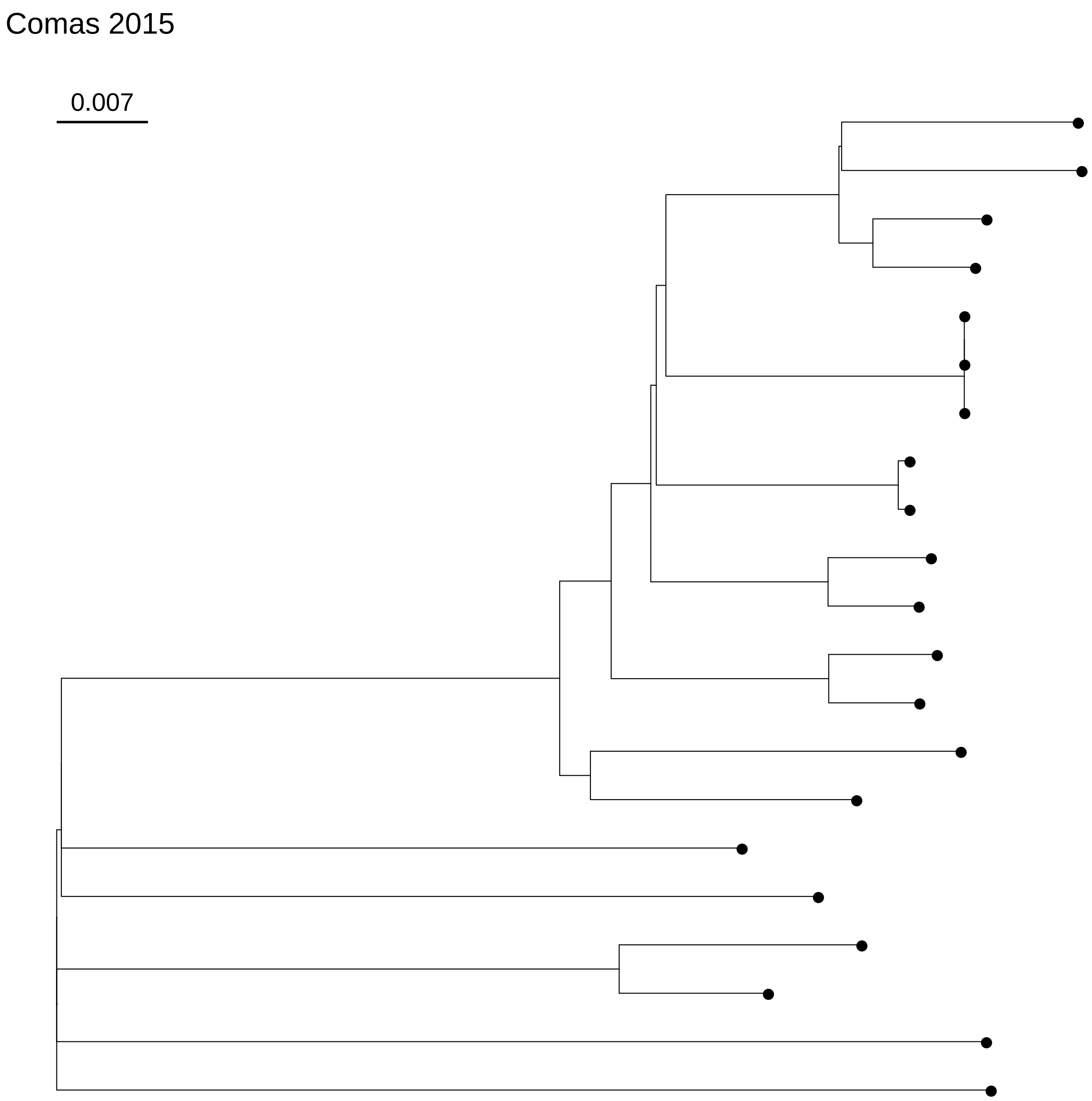


**Supplementary Figure 43.** Phylogenetic tree for the data set Comas 2015, the scale in expected nucleotide changes per polymorphic site. The tree was rooted with the reconstructed ancestral genome of the *Mycobacterium tuberculosis* complex (not depicted in the tree).

Eldholm 2015

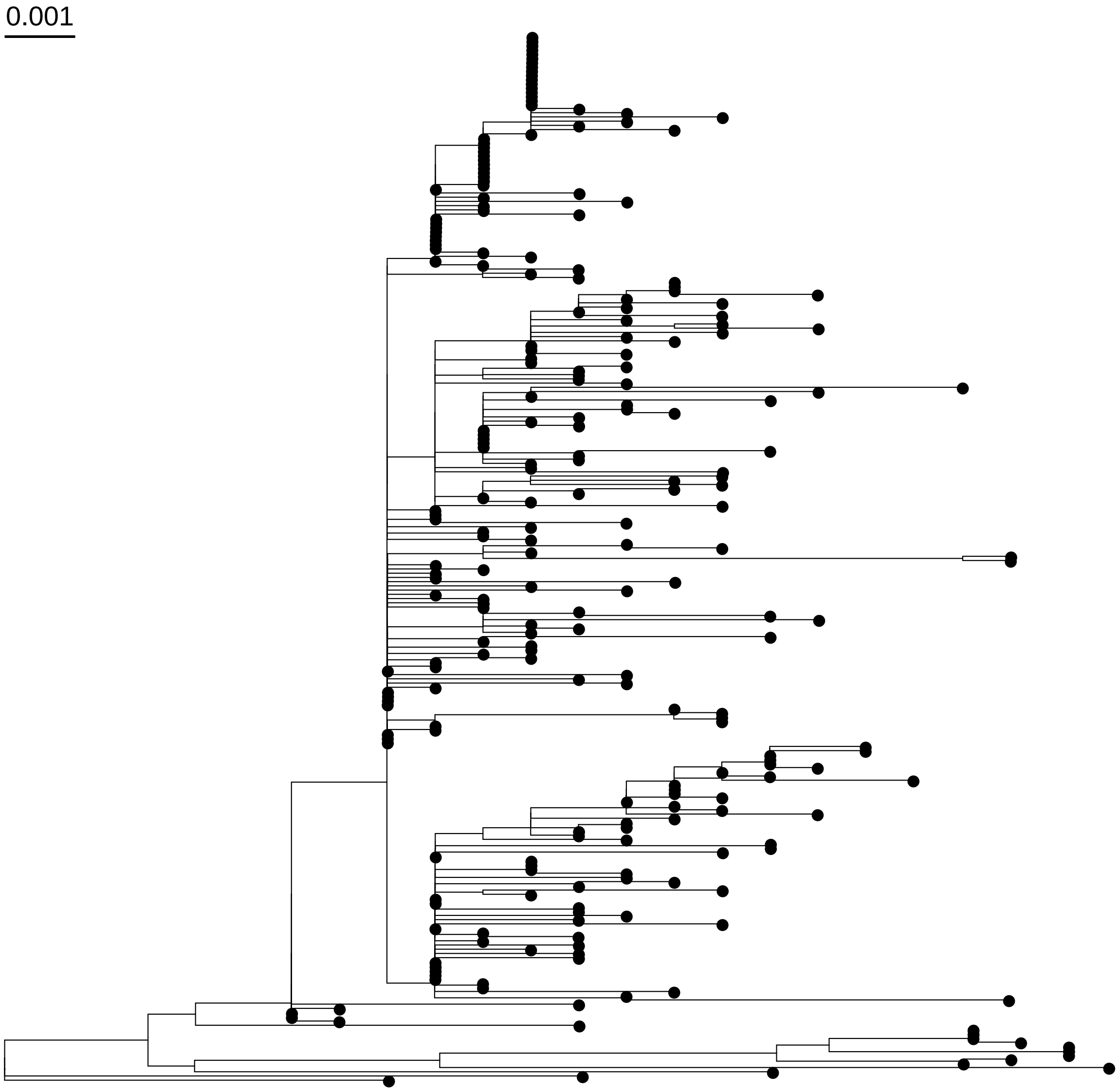


**Supplementary Figure 44.** Phylogenetic tree for the data set Eldholm 2015, the scale in expected nucleotide changes per polymorphic site. The tree was rooted with the reconstructed ancestral genome of the *Mycobacterium tuberculosis* complex (not depicted in the tree).

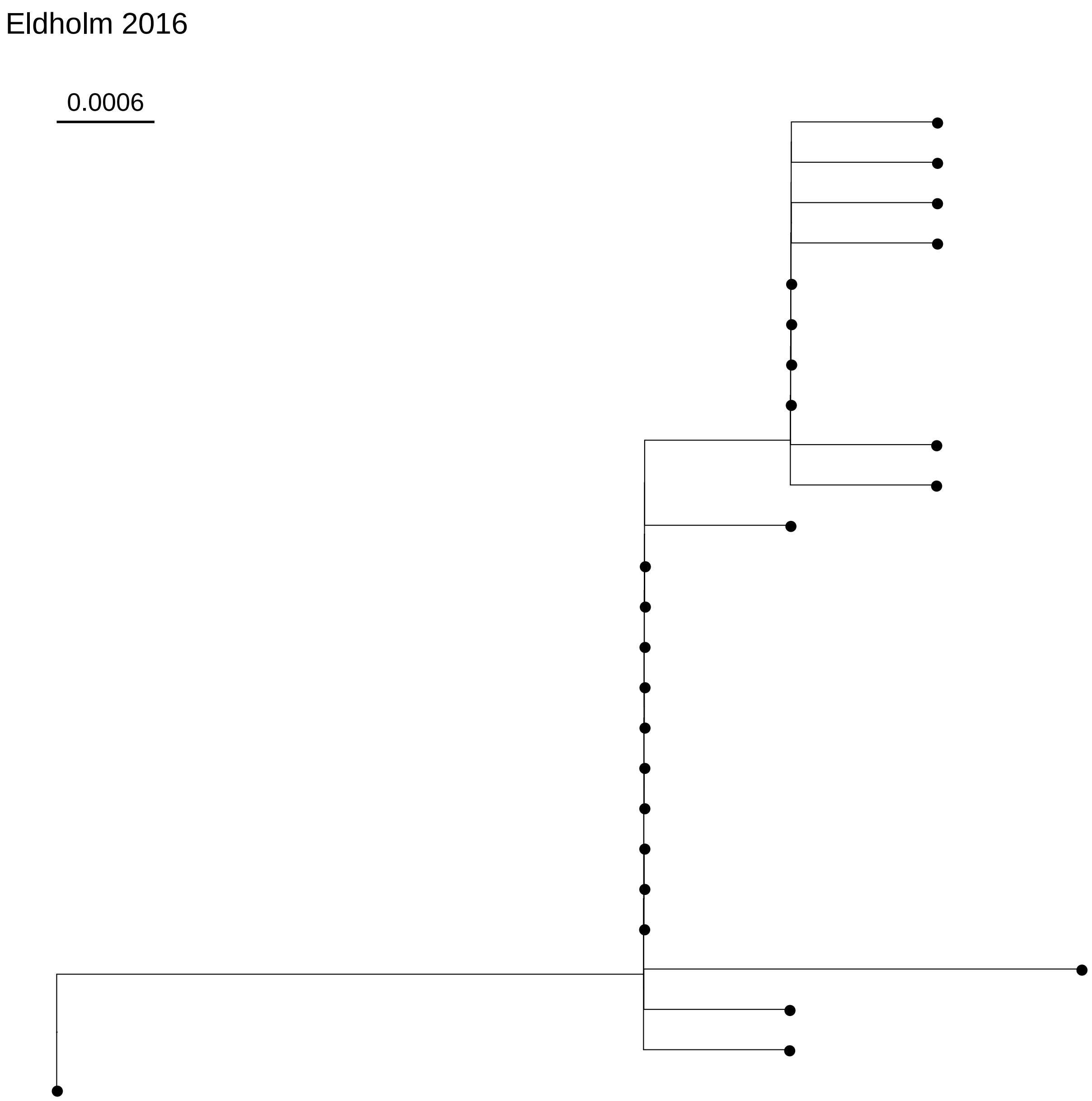


**Supplementary Figure 45.** Phylogenetic tree for the data set Eldholm 2016, the scale in expected nucleotide changes per polymorphic site. The tree was rooted with the reconstructed ancestral genome of the *Mycobacterium tuberculosis* complex (not depicted in the tree).

Folkvardsen 2017

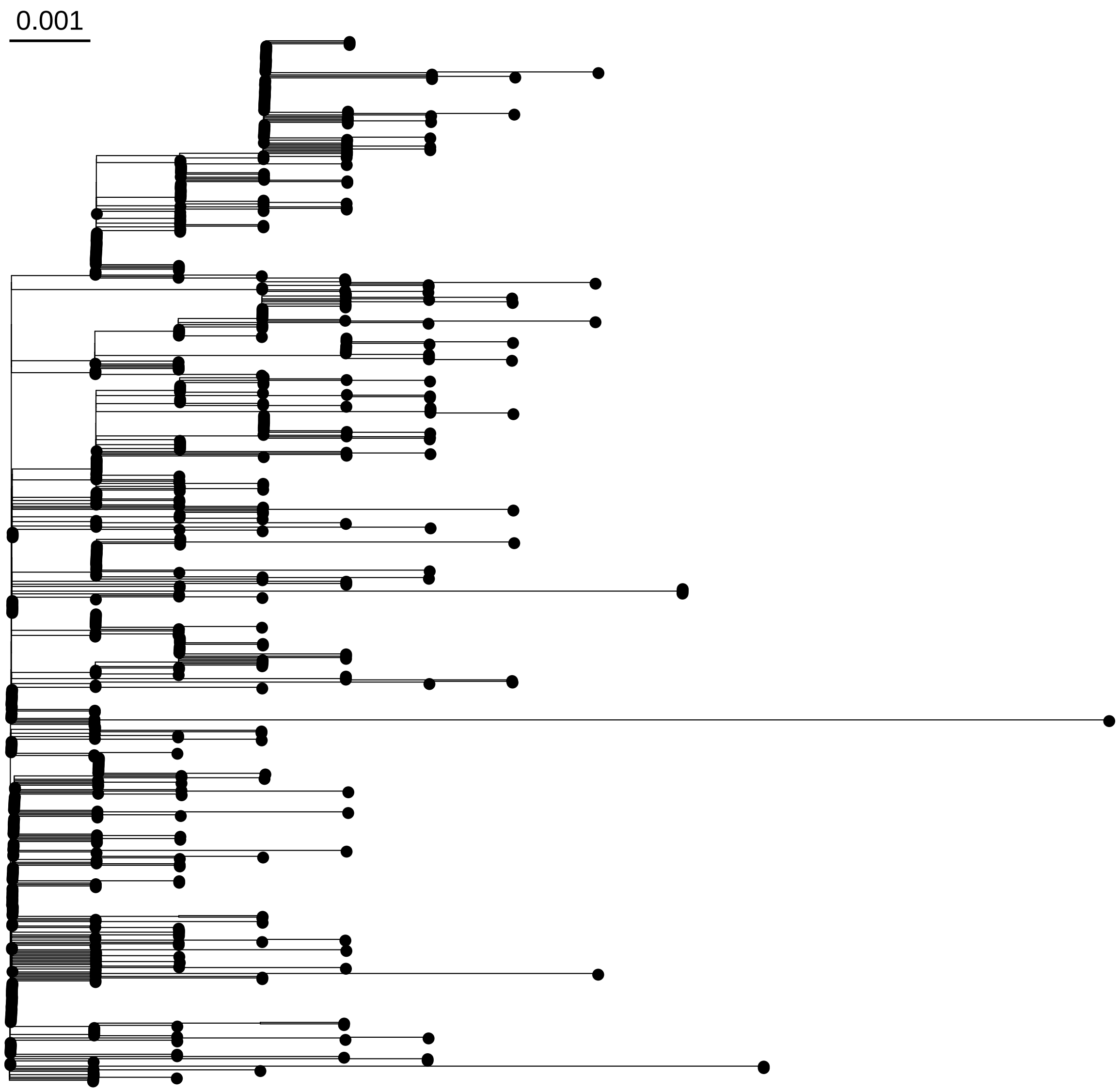


**Supplementary Figure 46.** Phylogenetic tree for the data set Folkvardsen 2017, the scale in expected nucleotide changes per polymorphic site. The tree was rooted with the reconstructed ancestral genome of the *Mycobacterium tuberculosis* complex (not depicted in the tree).

Lee 2015

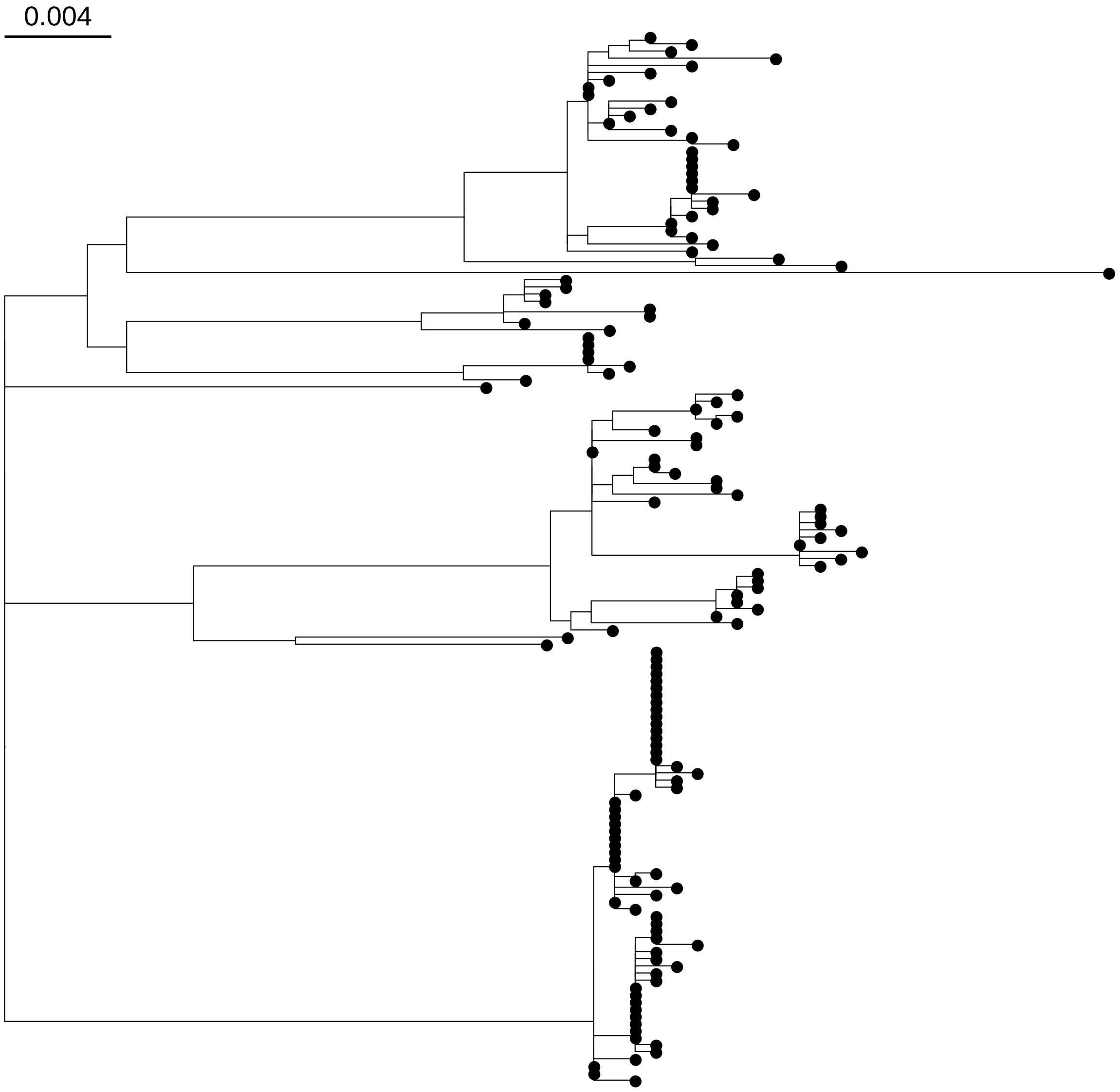


**Supplementary Figure 47.** Phylogenetic tree for the data set Lee 2015, the scale in expected nucleotide changes per polymorphic site. The tree was rooted with the reconstructed ancestral genome of the *Mycobacterium tuberculosis* complex (not depicted in the tree).

Roetzer 2013

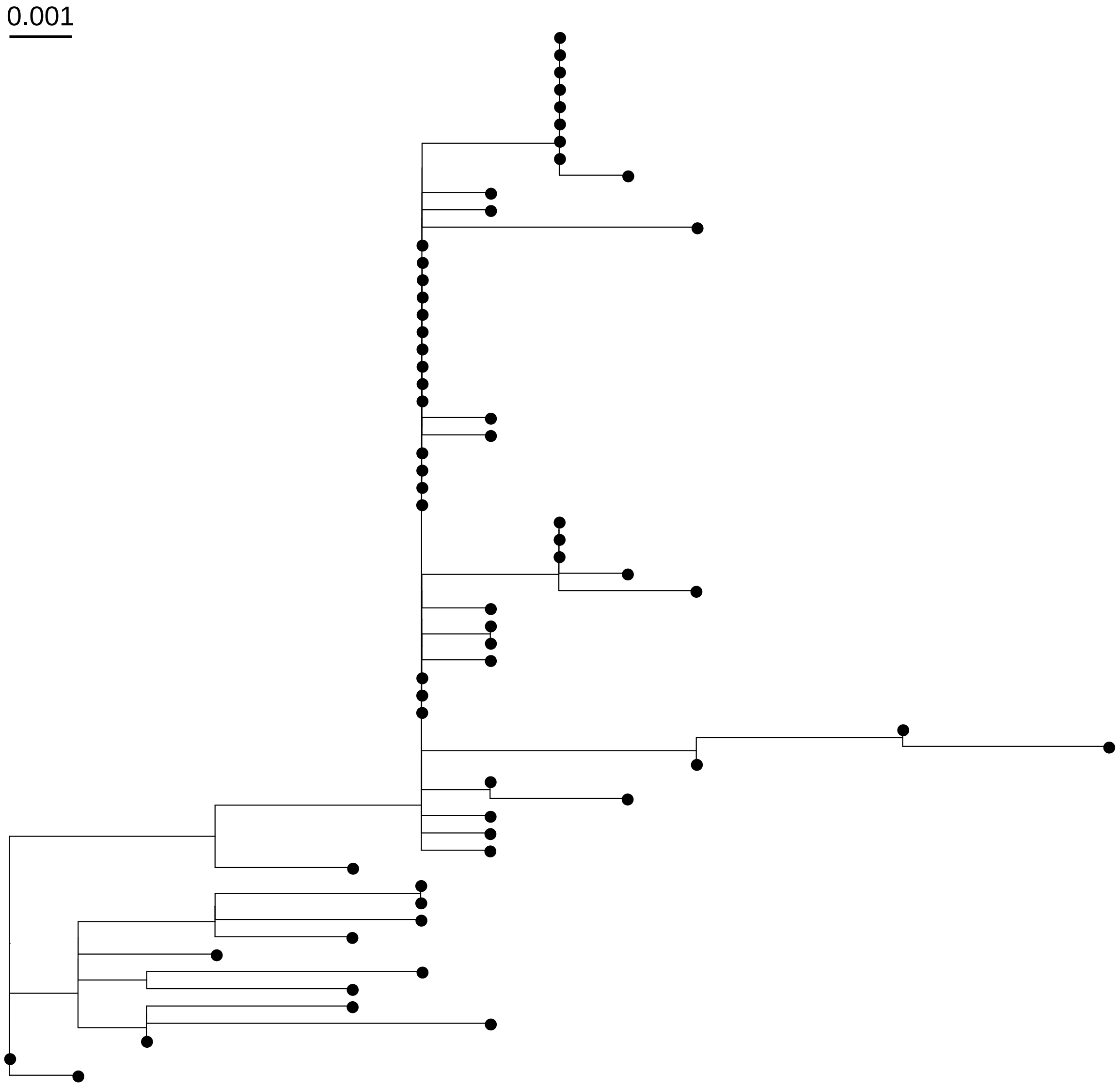


**Supplementary Figure 48.** Phylogenetic tree for the data set Roetzer 2013, the scale in expected nucleotide changes per polymorphic site. The tree was rooted with the reconstructed ancestral genome of the *Mycobacterium tuberculosis* complex (not depicted in the tree).

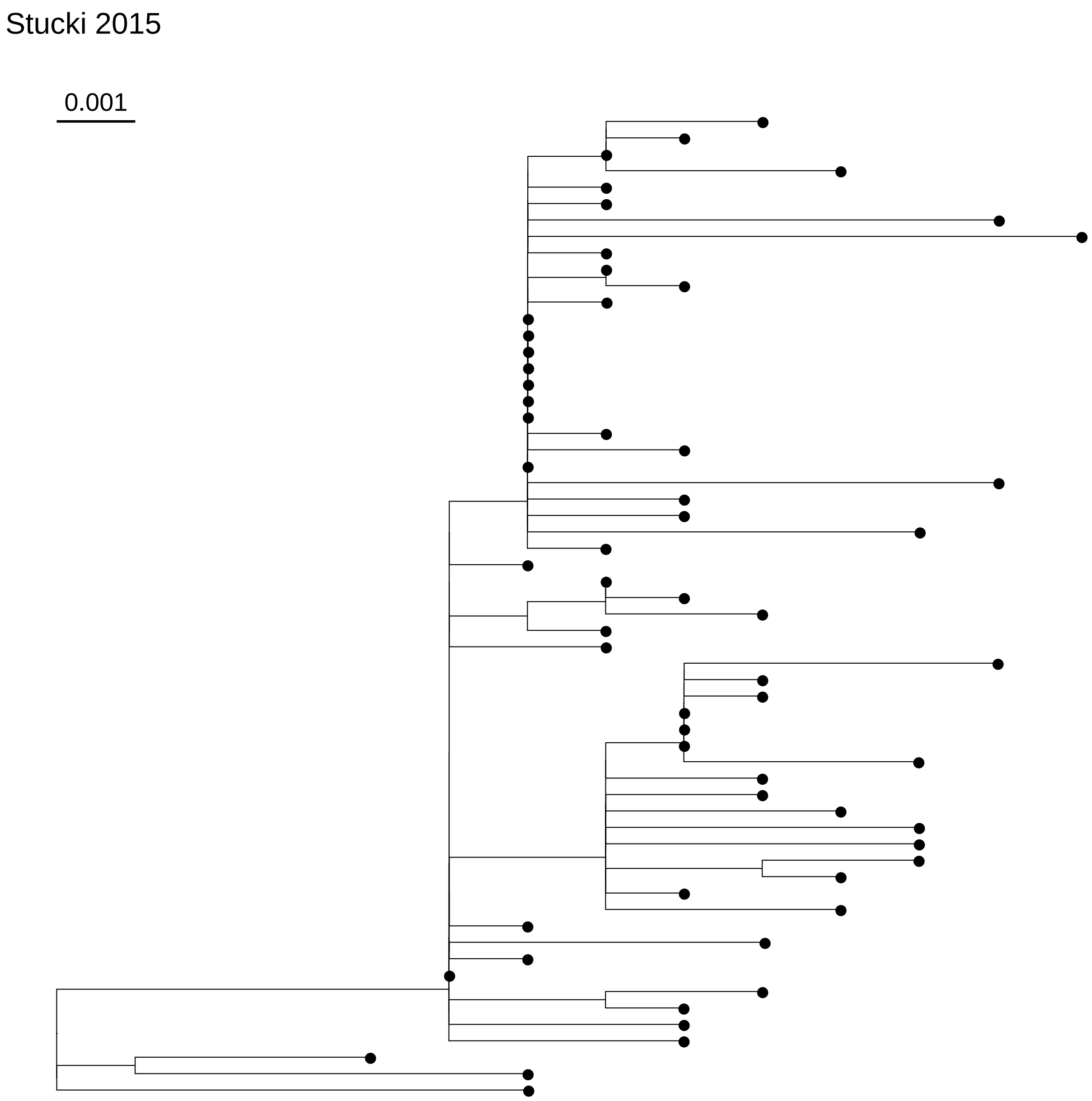


**Supplementary Figure 49.** Phylogenetic tree for the data set Stucki 2015, the scale in expected nucleotide changes per polymorphic site. The tree was rooted with the reconstructed ancestral genome of the *Mycobacterium tuberculosis* complex (not depicted in the tree).

Stucki 2016

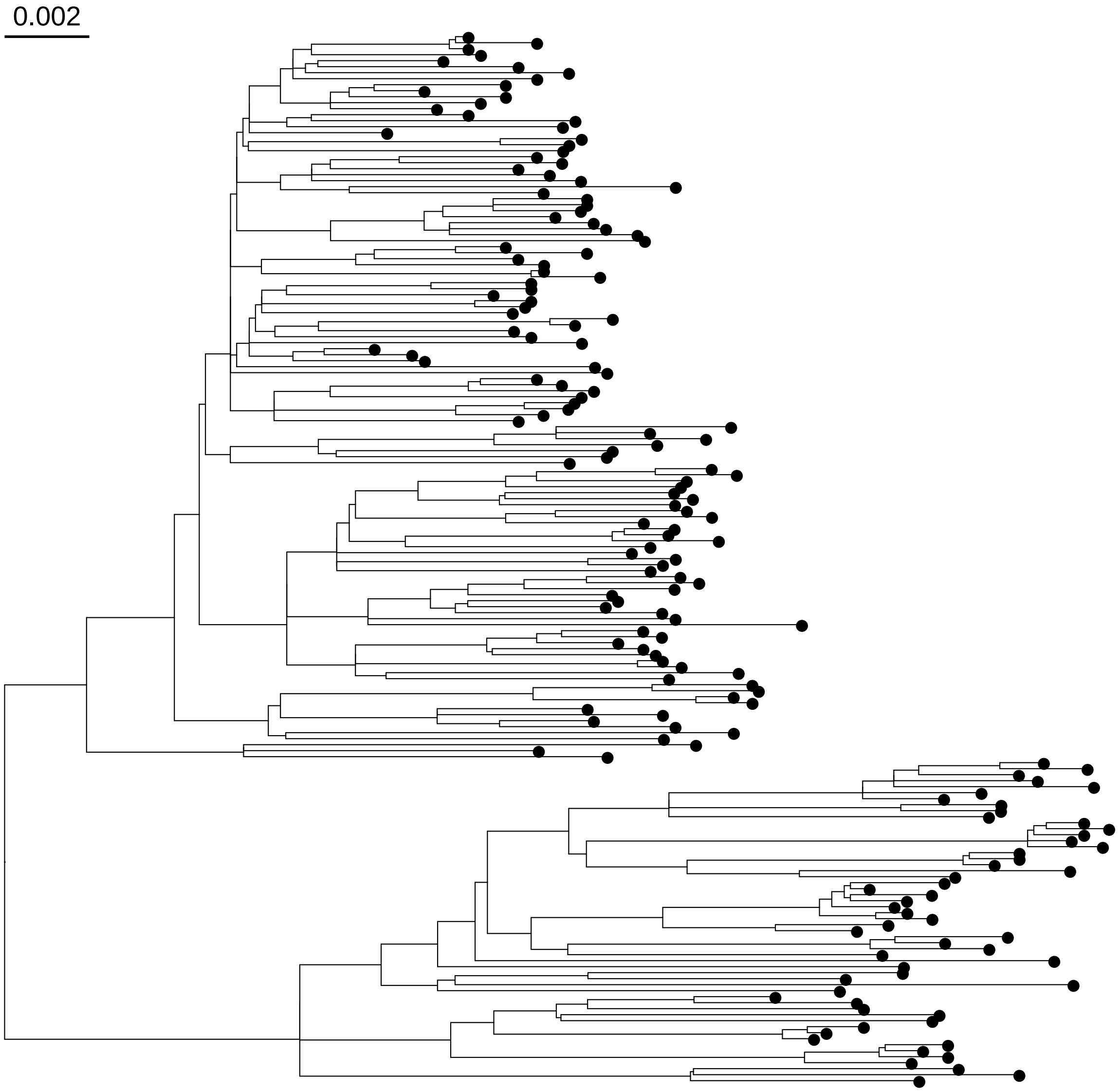


**Supplementary Figure 50.** Phylogenetic tree for the data set Stucki 2016, the scale in expected nucleotide changes per polymorphic site. The tree was rooted with the reconstructed ancestral genome of the *Mycobacterium tuberculosis* complex (not depicted in the tree).

Shitikov 2017

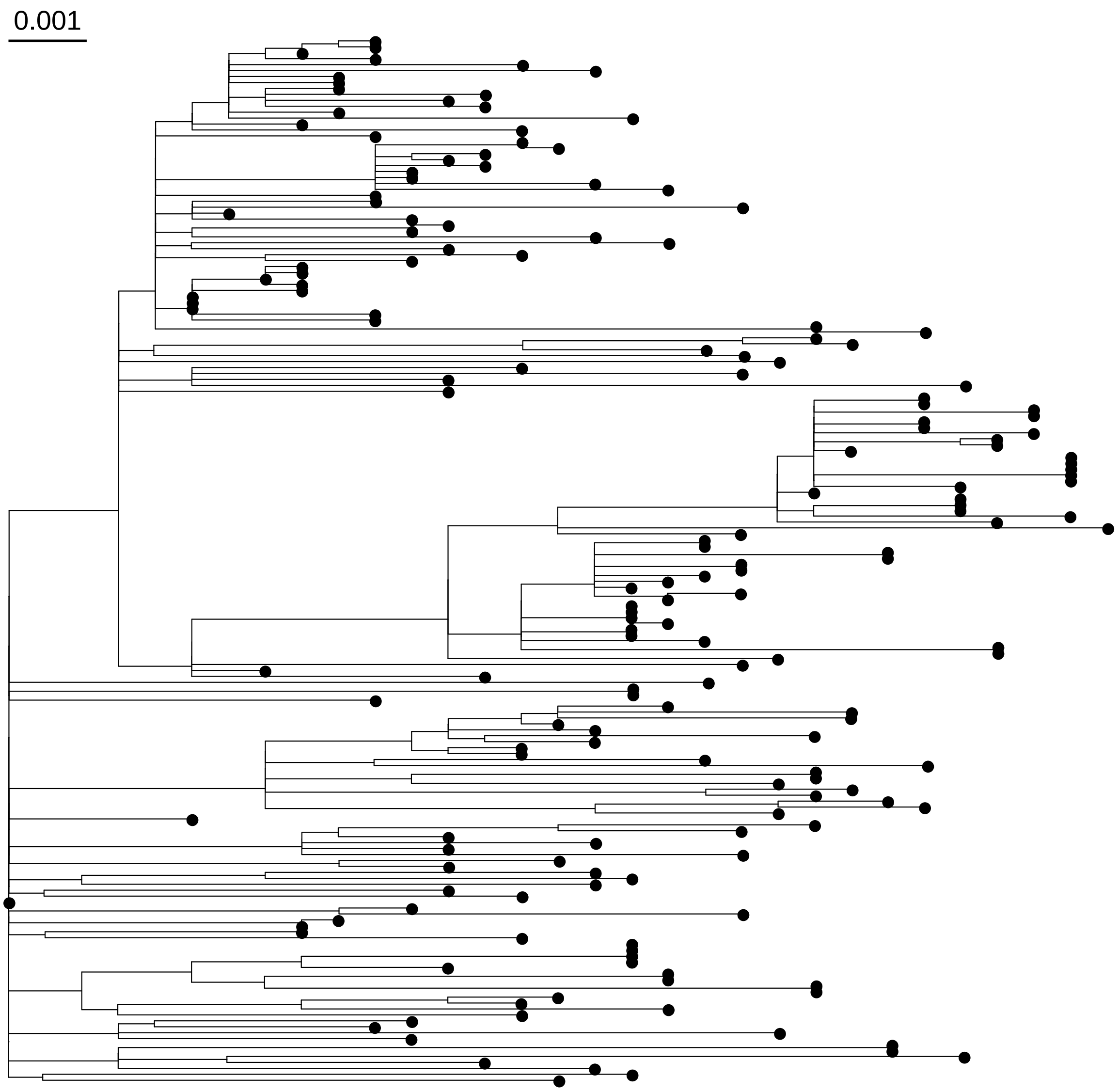


**Supplementary Figure 51.** Phylogenetic tree for the data set Shitikov 2017, the scale in expected nucleotide changes per polymorphic site. The tree was rooted with the reconstructed ancestral genome of the *Mycobacterium tuberculosis* complex (not depicted in the tree).
